# Cell type-specific consequences of mosaic structural variants in hematopoietic stem and progenitor cells

**DOI:** 10.1101/2023.07.25.550502

**Authors:** Karen Grimes, Hyobin Jeong, Amanda Amoah, Nuo Xu, Julian Niemann, Benjamin Raeder, Patrick Hasenfeld, Catherine Stober, Tobias Rausch, Eva Benito, Johann-Christoph Jann, Daniel Nowak, Ramiz Emini, Markus Hoenicka, Andreas Liebold, Anthony Ho, Shimin Shuai, Hartmut Geiger, Ashley D. Sanders, Jan O. Korbel

## Abstract

The functional impact and cellular context of mosaic structural variants (mSVs) in normal tissues is understudied. Utilizing Strand-seq, we sequenced 1,133 single cell genomes from 19 human donors of increasing age, revealing a heterogeneous mSV landscape in hematopoietic stem and progenitor cells (HSPCs). While mSV clonal expansions are confined to individuals over 60, *de novo* mSV formation occurs consistently across age, frequently leading to megabase-scale segmental aneuploidies. Cells harboring subclonal mosaicism show evidence for increased mSV formation. To enable high-resolution cell-typing of each Strand-seq library, we generated single-cell MNase-seq reference datasets for eight distinct HSPCs. Subclonal mSVs frequently exhibit enrichment in myeloid progenitors, and single-cell multiomic analysis suggests that these mSVs result in recurrent dysregulation of pathways related to proliferation and metabolism, including Ras signaling and lipid metabolism. The comprehensive mSV landscape identified in this study implicates mSVs in cell type-specific molecular phenotypes, establishing a foundation for deciphering links between mSVs, aging, and disease risk.

## Introduction

Somatic mutations arise in virtually all tissues and accumulate throughout the human lifespan^1–6^. While important insights into mosaic single nucleotide variants (SNVs) have been unveiled, progress on understanding the cell type-specific impact of mSVs, which include deletions, duplications, balanced inversions, and complex DNA rearrangements, has lagged much further behind^6, 7^. Findings from pan-cancer studies indicate that somatic structural variants more frequently serve as a cancer driver than SNVs^8, 9^. In addition, studies performing bulk and clone-based sequencing in the blood compartment of healthy donors have reported associations between mosaic copy-number aberrations (CNAs) and abnormal blood cell counts, cancer susceptibility, and cardiovascular disease^2, 7, 10–12^. These data suggest a potentially important contribution of mSVs to mosaicism in normal tissues, including hematopoiesis.

However, mSVs represent one of the most cryptic and challenging-to-ascertain classes of human genetic variation^7, 8^: Bulk whole genome sequencing (WGS) typically fails to detect mSVs with a clonal fraction (CF) lower than 30% and does not discriminate between cell types, whereas WGS of single cell-derived clones restricts the analysis to mSVs that are culturable *ex vivo*, which may result in the under-reporting of mSVs which lead to large segmental aneuploidy stretches^8, 13, 14^. While single-cell sequencing can access the widest CF range in principle, the resolution of commonly used methods restricts the reporting to large mosaic CNAs, while other variant classes including balanced and complex mSVs escape detection^15^. These challenges extend to single-cell multi-omics methods capable of concurrent mSV discovery and molecular phenotyping in each cell^8^. As a consequence, the cell type-specific context and functional impact of mSVs in normal tissues is poorly understood^6, 7^.

Here we employ Strand-seq, a haplotype-resolved single-cell sequencing technique^14, 16, 17^, to study the cellular context and functional impact of mSVs in a normal tissue. We focus on the blood compartment, which is maintained by a hierarchically-organized stem cell population. In this compartment, mosaic mutations including SNVs and CNAs are common among older donors^6, 7^. We previously reported that Strand-seq offers access to subclonal balanced, unbalanced, and complex structural variants in cancer cells^14^. Furthermore, Strand-seq simultaneously yields nucleosome occupancy (NO) profiles from the same cell, a readout revealing the functional consequences of somatic chromosomal rearrangements^18^. Our study reports on the first application of this innovative single–cell multiomic technology to investigate somatic mosaicism in a phenotypically normal tissue. By applying Strand-seq to donors of varying age, we unveil a wide spectrum of mSVs classes in human HSPCs. In 1 out of every 43 cells, we detect *de novo* mSVs, which emerge regardless of age. We report hotspots of mSV formation in HSPCs, which coincide with regions exhibiting frequent sister chromatid exchanges^16, 18^ (SCEs). In addition, by constructing an NO reference dataset via single-cell MNase-seq^19^ (scMNase-seq), we resolve the cell-type identity of mSV-bearing cells and show that mSVs are commonly biased for myeloid progenitor cells. We find that while heterogeneous with respect to the loci at which they occur, mSVs are recurrently associated with aberrant cell-cycle pathways, Ras signaling and lipid metabolism, which implies that they may affect common aging-associated molecular pathways.

## Results

### Diverse single-cell resolved mSV landscapes in normal human HSPCs

To allow for profiling of mSVs in human blood cells in a cell-type resolved manner, we devised an experimental workflow for single-cell mSV detection in primary HSPCs (**Fig. 1a**). Our sample cohort consists of 19 normal donors from newborn to 92 years of age, with *N*=3 umbilical cord blood (UCB) and *N*=16 bone marrow (BM) samples (**Methods**). We isolated viable CD34+ HSPCs from these samples (**Fig. S1**), and cultured them *ex vivo* for one cell division to allow for Strand-seq library preparation. Following quality control, we identified 1,133 high-quality single-cell libraries, with a mean of 432,282 uniquely mapped fragments per cell (**Fig. S2**; **Table S1**). We used scTRIP^14^ to perform single-cell discovery of mSVs and chromosomal aneuploidies (collectively referred to as ‘mosaicisms’) in these libraries (**Fig. 1b**). Altogether, we identify 51 independently arisen mosaicisms in our cohort (mean per donor=2.7; range 0–8; mosaicisms detected in 16/19 [84%] donors), which include: 22 deletions, 12 duplications, 3 complex mSV events, a balanced inversion, and 13 sex chromosome losses (**Fig. 1b**, **Table S2**). These mosaicisms affect 17/24 chromosomes, and exhibit no particular enrichment except for the Y chromosome, which shows one or more independent losses of Y (LOYs) in 8/12 (67%) male donors.

**Figure 1:**
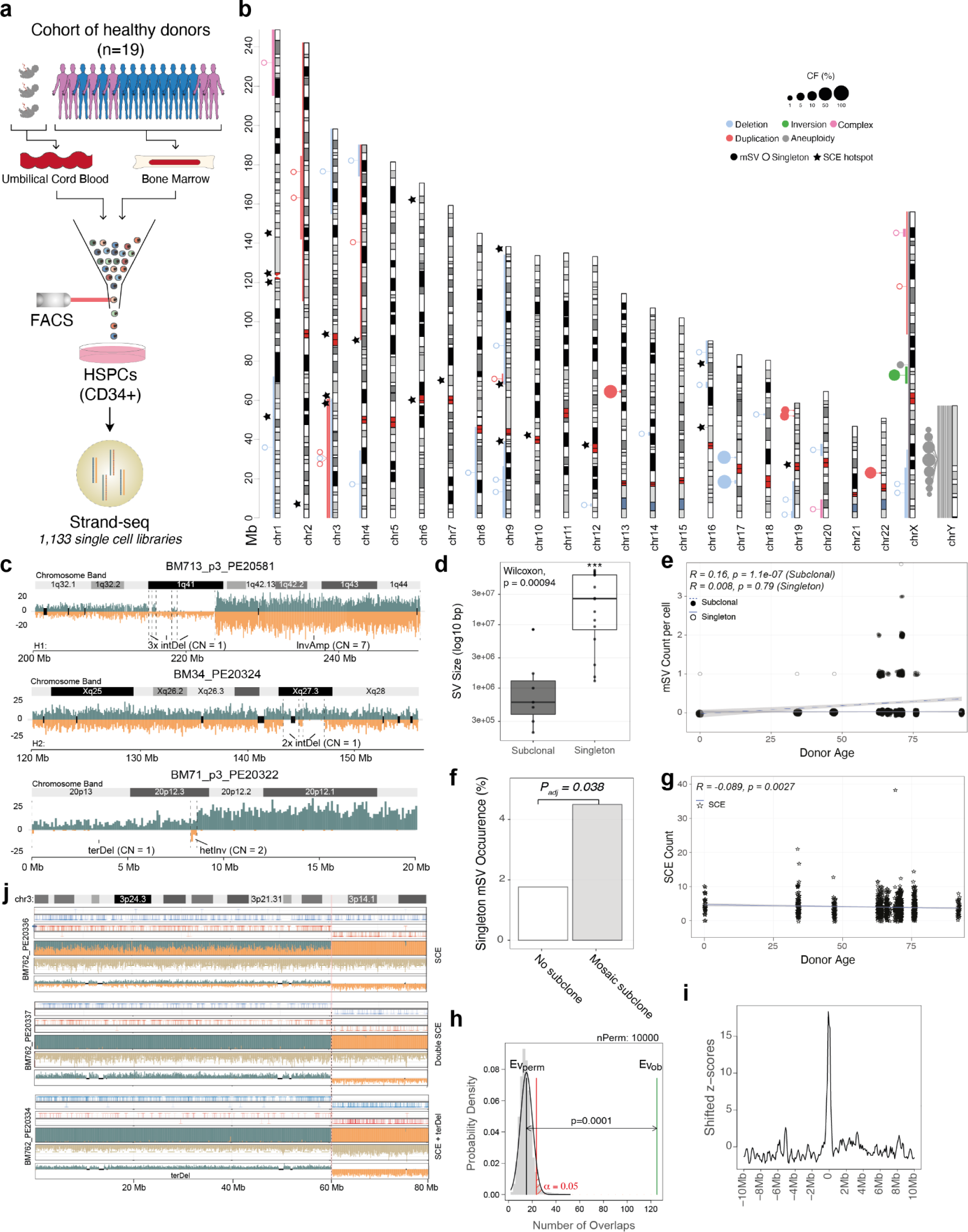
Human HSPCs acquire a wide diversity of mSVs with age, without increased genomic instability. **a**) Cohort and experimental workflow used for profiling somatic mosaicisms in human HSPCs. **b**) Genome-wide karyogram of mSVs identified in human HSPCs. Bars indicate the size of identified mSVs, colour indicates the class, and the relative size of the bubble linked to the middle of each mSV indicates the cellular fraction (CF) of the mSV. Filled circles denote subclonal mSVs, while unfilled are singleton mSVs. Stars indicate bins significantly enriched for SCEs. **c**) Examples of singleton complex rearrangements identified in the cohort. Copy-number estimates in regions affected by complex mSVs are indicated next to the respective segments. **d**) Singleton mSVs are significantly larger, when comparing mean total affected base pairs, than mSVs (Wilcoxon rank-sum test; P = 0.00094). **e,g**) Jitter plots depicting trends in the number of subclonal/singleton mSVs (**e**) and SCEs (**g**) across age in human HSPCs (R = correlation coefficient calculated from the number of mSVs/SCEs given the donor age). **f**) Barplot of the incidence of singleton mSVs (y-axis) in cells with or without mosaicism. Significance is tested with an FE test. **h**) Results of permutation test shuffling singleton mSV breakpoints (100kb confidence interval) and SCE hotspots (200kb bin) genome-wide for 10,000 permutations showing a significantly greater than expected number of overlaps between the two. Adjusted p-value indicates the significance of the difference between the permuted (black line) and actual (green line) number of overlaps between singleton mSV breakpoints and SCE hotspots. **i**) Local Z-score of enrichment of overlaps between singleton mSV breakpoints and SCE hotspots from BM/UCB data. mSV breakpoints are shifted in windows of 100 kb to 10 Mb +/- the bin in which an SCE hotspot is located, and the enrichment Z-score plotted each time. Additional permutations plotted in **Fig. S6**. **j**) Strand-seq data showing recurrent SCE and mSV co-occurrence at the SCE hotspot and common fragile site (CFS) *FRA3B* in donor BM762.

We investigated the subclonal composition in each of the 19 donors (**Table S2)**. Out of all mosaicisms analyzed, 32 are detected in only one cell (’singleton mosaicism’), while the remaining 19 constitute subclones with CFs of 1.6%-56.1% (’subclonal mosaicism’). While we find no singleton or subclonal losses of an autosome, subclones with sex chromosome losses (*N*=12 LOY; *N*=1 losses of X (LOX)) reached CFs up to 46.4%. We focused most of our investigation on the 38 mSVs (*i.e.,* non-LOY/LOX mosaicisms) in our dataset, which we mapped with 200 kb resolution, utilizing the scTRIP^14^ framework. Examination of these mSVs, highlighted notable differences between singleton and subclonal events. First, 21/31 singleton mSVs (67.74%) exhibit terminal gains or losses, whereas all 7 subclonal mSVs are composed of interstitial rearrangements. Second, all of the 3 complex mSV events are singletons, and these harbor a multitude of DNA rearrangements affecting a single chromosomal haplotype – which includes a breakage fusion bridge cycle-mediated^14^ mSV on the chromosome 20p-arm, as well as a terminal sister chromatid fusion-mediated amplification to a copy-number of 7 on 1q (**Fig. 1c**). Third, singleton mSVs are ∼17.6 times larger on average than subclonal mSVs (mean size of 36.9 and 2.1 Megabasepairs (Mb), respectively; *P*=0.0009, Wilcoxon rank-sum test; **Fig. 1d**). These data indicate that singleton mSVs, detected in 1 out of every 43 HSPCs, bear the characteristics of newly arisen, *de novo* mSVs (see **Supplementary Notes**).

We next analyzed the occurrence of mosaicisms with respect to donor age. We detect a significant increase with donor age both for subclonal mSVs (Pearson’s correlation *R=*0.16; *P*=1.1e-07; **Fig. 1e**) and for subclonal whole chromosome losses (*R*=0.087, *P*=0.0034); with subclonal mSVs occurring exclusively in donors aged 63 or older. These data align with prior reports of age-related CNA mosaicism in the blood of healthy individuals^2, 7, 20, 21^. However, we find that the occurrence of singleton mSVs – a type of mosaicism not previously amenable to detection – is not significantly correlated with donor age (*R*=0.008; *P*=0.79; **Fig. 1e**), suggesting that HSPCs acquire mSVs regardless of age. We also examined the full dataset for co-occurrence of mSVs. Notably, we find evidence for an elevated number of *de novo* mSVs (*i.e.* singleton mSVs) in cells that already contain mosaicisms, compared to WT cells (Fisher’s exact (FE) test; 4.76% vs. 1.96%; *P*=0.038; 2.4-fold elevated; **Fig. 1f**). These data suggest that while SV formation in HSPCs occurs consistently across age, it remains possible that cells harboring mosaicisms are ‘predisposed’ to accumulate further mSVs.

### Investigation of mSV breakpoint regions unveils hotspots of mSV formation in HSPCs

The distinctive capabilities of Strand-seq to detect sister-chromatid exchanges (SCEs) allows accurate mapping of DSBs that were subject to DNA repair and resolved by a crossover event^16^. Since DSBs play a crucial role in triggering DNA rearrangement formation^8, 22^, we therefore examined the correlation between DSB acquisition and donor age. We mapped SCEs across all 1,133 single cells (**Methods**), and identified 4,528 in total, averaging 4 SCEs per single cell (**Fig. S3**), consistent with prior reports^23^. We find an inverse correlation between SCE abundance and age (*R*=-0.089; *P*=0.0027; **Fig. 1g**), with on average 4.6 SCEs/cell in individuals < 60 years of age compared to 3.9 SCEs/cell in individuals >60, independent of tissue of origin (**Fig. S4**). Collectively the data imply that, despite the accumulation of subclonal mSVs with age, HSPCs from older donors do not exhibit increased mSV or SCE formation; with mSVs rather arising consistently over an individual’s lifetime.

Since DNA rearrangement formation can be mediated by the local genomic context^8^, we analysed the location of SCE and mSV breakpoints. We find SCEs are unevenly distributed in the genome of HSPCs (**Fig. 1b**), with 6.67% (302/4,528) clustering into 20 SCE ‘hotspots’ (**Methods**, **Fig. S5, Table S3**). Only 25% (5/20) of the SCE hotspots coincide with a previously reported common fragile site (CFS) (**Table S4**) – genomic regions vulnerable to DSB formation during replication stress^24^. We also find that 3% (133) of the 4,528 SCEs intersect with the breakpoint regions of an interstitial mSV, with these breakpoints showing significant overlap with SCEs (*P*<0.0001, derived from 10,000 permutations; **Fig. 1h,i**, **Fig. S6, Table S3**) and CFSs (*P*<0.0002) (**Fig. S6**). Notably, we identify recurrent mSVs at SCE hotspots – not all of which correspond to known CFS (*e.g.*, *FRA3B*) (**Fig. 1b,j; Fig. S7; Table S2, S3, S4**) – suggesting that SCE hotspots (*e.g.,* chromosome 9:68,300,000-68,500,000) likely represent DNA rearrangement hotspots in HSPCs. These findings highlight the association between SCEs and mSV formation in HSPCs, suggesting that SCEs and mSVs can arise through shared processes such as break-induced replication and non-homologous end-joining^22^.

### High-resolution cell typing of HSPCs using indexed scMNase-seq-based NO reference datasets

The cell-type-specific molecular impact of mSVs is yet to be established. We set out to examine this in the context of the blood compartment, using the scNOVA tool designed for cell-type classification based on single-cell NO data, which we previously showed to be effective for cell-typing of Strand-seq libraries into various cell lines and tissue types^18^. In spite of the availability of various multiomics data from the blood compartment^25^, NO reference datasets are lacking. We thus generated single-cell NO reference profiles for both UCB- and BM-derived HSPCs, for 8 distinct HSPC cell types. These cell types include: hematopoietic stem cells (HSCs), multipotent progenitors (MPPs), lymphoid-primed multipotent progenitors (LMPPs), common lymphoid progenitors (CLPs), plasmacytoid-dendritic cells (pDC), common myeloid progenitors (CMPs), granulocyte-macrophage progenitors (GMPs), and megakaryocyte–erythroid progenitors (MEPs) (**Fig. 2a, Fig. S8**). Using predefined immunophenotypes (**Table S5; Fig. S8**) we index-sorted HPSCs, and devised a preamplification-free scMNase-seq protocol to generate single-cell NO profiles for each cell-type separately (**Methods**).

**Figure 2:**
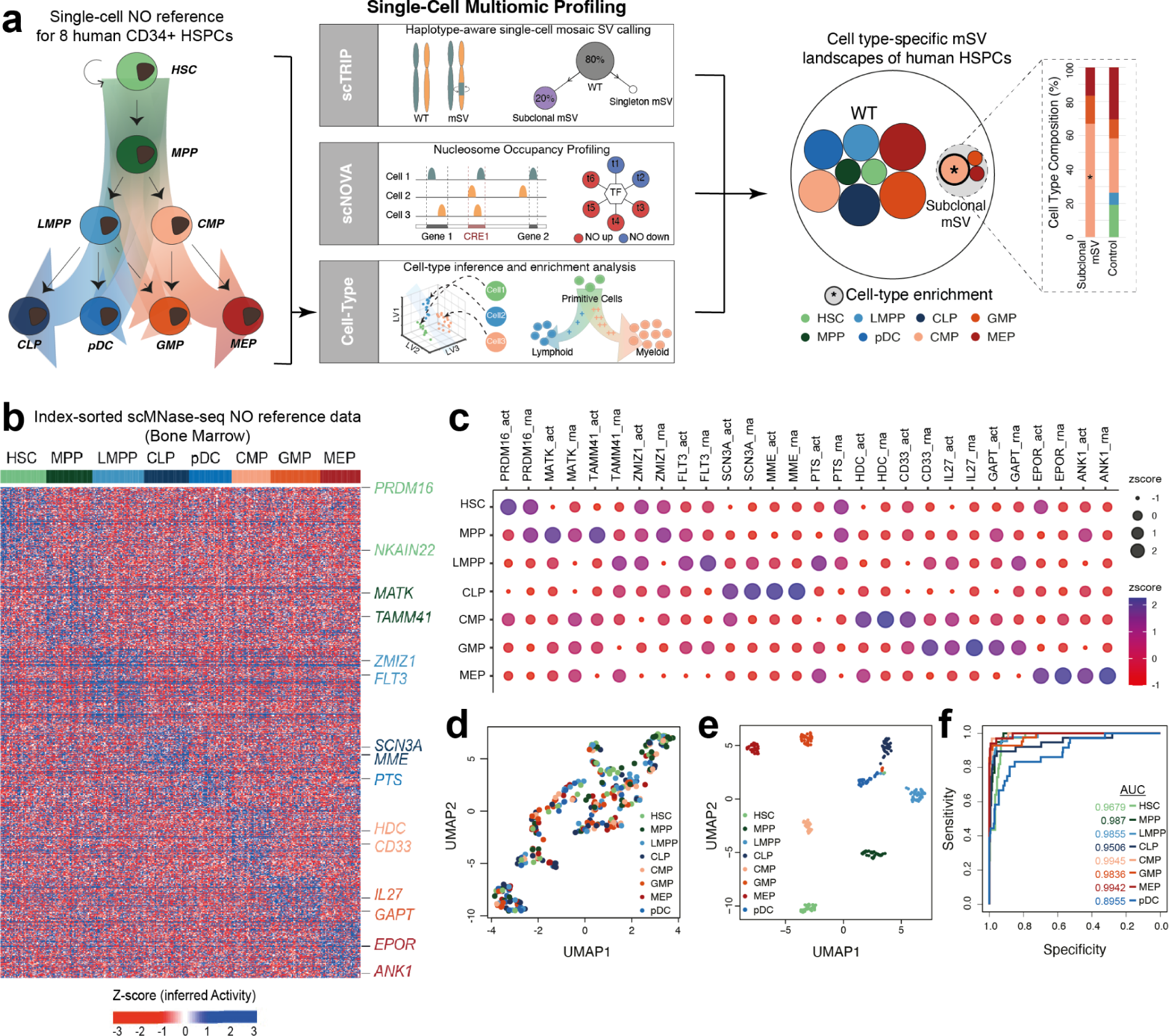
scMNase-seq atlases for 8 distinct HSPCs enable cell type-aware single-cell multiomic profiling of mSV landscapes **a**) Computational single-cell multiomic profiling workflow used to characterize cell-type specific mSV landscapes in HSPCs using Strand-seq, which involves single-cell mSV discovery (scTRIP^14^), single-cell NO analysis to identify mSV functional effects (scNOVA^18^), and single-cell cell-typing (informed by scMNase-seq-derived classifiers). **b**) Construction of BM and UCB-specific NO reference datasets, based on subjecting eight distinct HSPC cell-types (BM-based reference is depicted here; the UCB-based reference is shown as **Fig. S7**) to index sorting using FACS followed by scMNase-seq. Heatmap of single cell NO of gene bodies of 305 single BM HSPCs. The 819 signature genes depicted (rows) allow for discrimination between 8 distinct HSPC cell-types (columns). Cells are grouped and colour-coded by immunophenotype, determined by FACS. Example marker genes for each cell type are shown to the right of the heatmap, colour-coded by the defined cell type. Differential NO of marker genes is represented by Z-scores. **c**) Comparison of inferred gene activity^18^ (act; inverse NO, using scMNase-seq) and gene expression (RNA-seq) for the representative classifier genes from the BM scMNase-seq reference. Publicly available RNA-seq data was used for this analysis^28^. Gene activity at gene bodies was inferred using the NO Z-score multiplied by (-1). Color and the dot sizes reflect the Z-score of inferred gene activity and RNA expression, respectively. **d**) Unsupervised UMAP dimensionality reduction of BM HSPC scMNase-seq data. **e**) Supervised UMAP dimensionality reduction of BM HSPC scMNase-seq data using the BM cell-type classifier. **f**) ROC curve showing leave-one-out cross-validation of the BM cell-type classifier’s performance using single cell NO patterns.

We obtained 480 high-quality scMNase-seq reference libraries (**Table S6**): 305 from BM-derived cells (1 donor) and 175 from UCB-derived cells (5 donors) (**Table S1**). Using scNOVA, we identify 899 and 819 genes exhibiting cell-type-specific NO patterns in their gene bodies in the UCB- and BM-derived datasets, respectively, indicative for cell-type specific gene activity^18^ (**Fig. 2b**, **Fig. S9**). These included several previously identified as marker genes in HSPCs (**Table S7A,B**): For instance, using the BM-derived NO reference dataset, we infer that the canonical marker *MME* (CD10) shows increased activity in CLPs^26^, while *HDC* exhibits increased gene activity in CMPs, reflecting its involvement in myeloid lineage priming^27^. We also observe differential NO at several genes not previously reported as HSPC markers – such as *SH2D4B* and *FAT3* (**Table S7A**) – indicative for potentially novel cell-type-specific gene activity. Overall, the gene activities inferred from NO are broadly consistent with previously reported RNA-seq expression data^28^ (**Fig. 2c**), supporting the applicability of our NO reference datasets for cell typing.

Harnessing these NO marker gene sets, we built novel cell-type classifiers to enable accurate cell-typing of discrete HSPCs from Strand-seq data. We utilized NO measurements within the gene bodies of marker genes as features for developing a supervised model using partial linear square discriminant analysis (**Fig. 2d-f**, **Fig. S9, Table S7A, B; Methods**). The classifiers provide excellent accuracy, with an average area under the curve (AUC) of 0.97 for BM and 1.00 for UCB, as estimated by leave-one-out cross-validation (**Fig. 2d, Fig. S9**). UMAP projections of the latent variables further corroborate the discriminatory power of these classifiers in comparison to an unsupervised classification (**Fig. 2e,f**, **Fig. S9**). These data establish that both NO based classifiers offer robust cell-typing of single-cell NO data, providing a new means for unraveling the cellular context of mSVs in HSPCs identified using Strand-seq.

### Subclonal mSVs commonly exhibit a lineage-bias towards myeloid HSPCs

Using the trained cell type classifiers, we performed cell-typing of each Strand-seq library (**Fig. 3a, Table S8**). We find that tissue-level cell abundances show consistency with previous studies^28–31^, such as an expanded HSC frequency amongst BM donors with age (from 8.1% to 80%; FDR-adjusted *P* (*P_adj_*)=0.013; mixed linear model analysis), and a greater abundance of MPPs in UCB donors vs BM^28, 29^ (37% in UCB vs. 0.1% in BM; *P_adj_*=2.45e-33; FE test; **Fig. S10**). MPPs, CLPs, and pDCs exhibit a lower prevalence than other cell-types, likely reflecting their natural scarcity in HSPCs^28^ and known challenges with sustaining these cells *in vitro*.^32^

**Figure 3:**
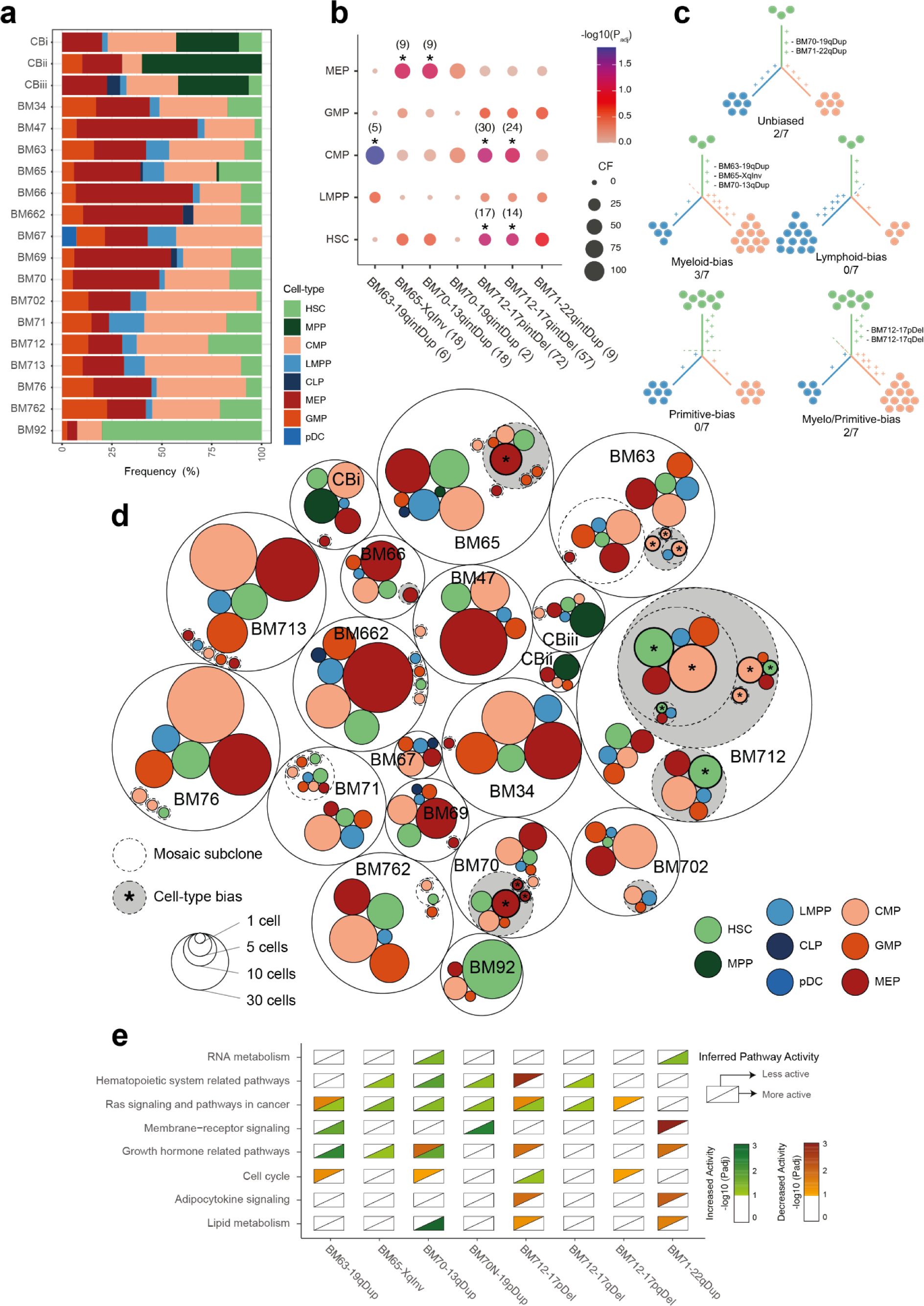
mSVs in human HSPCs frequently exhibit lineage-bias, in a donor-specific context. **a**) Predicted cell type composition per donor, in order of increasing age. **b**) Dotplot of results of the cell-type enrichment analysis for each mSV identified, showing the CF, enrichment and significance of each cell type per mSV sub-clone vs an idealized control. The number in brackets indicates the number of single cells of a given cell-type in a given sub-clone used to calculate the enrichment of that cell-type. Data here shows enrichment for single genotypes, for combined enrichments see **Fig. S12**. **c**) Summary of lineage biases observed in all subclonal mSVs across the cohort. **d**) Circle packing plot summarizing the mSVs and inferred cell type composition of each sub-clone for each of the 19 donors in this study. Each transparent circle with a solid outline represents a distinct sample. Transparent inner circles with dashed outlines represent distinct mSV sub-clones within that individual, while coloured circles denote the cell types contributing to that clone. Each circle is proportional to the total number of single cells composing that cell-type/sub-clone. A grey background identifies sub-clones which contain a significant (FDR 10 %) cell type-enrichment with respect to a control population made from randomly sampling karyotypically normal cells from donors over 60 years. **e**) Enrichment analysis of pathways grouped by Jaccard similarity, for subclonal mSVs across the cohort. Only groups of pathways enriched in 2 or more mSVs are shown. For all individual pathways, see **Fig. S14.** For all groups of pathways and details on Jaccard similarity-based grouping see **Fig. S32**.

Using these cell type classifications, we explored the cellular context of mSVs in normal cells, by testing subclonal mosaicisms for cell-type enrichment in each donor (**Methods**). Of 19 subclonal mosaicisms in our dataset, we find 8 (47%) display significant subclone-specific cell-type enrichment (FDR 10%; **Fig. 3b; Fig. S11, S12**). Out of these, we find that 5/7 (71%) subclonal mSVs exhibit significant cell-type-bias. Notably, while diverse in their affected locus and CF, subclonal mSVs are predominantly enriched in myeloid, or a combination of myeloid and primitive, cell-types: with either myeloid or myelo-primitive enrichment observed in all 5 cell-type biased mSV subclones in our study (**Fig. 3c**). These lineage-biased subclonal mSVs include: a 10 Mb inversion (Inv) on chromosome Xq enriched in MEPs (BM65); a 1 Mb duplication (Dup) on chromosome 13q enriched in MEPs (BM70); a 300 kb Dup on chromosome 19q enriched in CMPs (BM63); and two sequentially arisen deletions (Del) on chromosome 17p (1.2 Mb) and 17q (500 kb) enriched in both CMPs and HSCs (BM712).

LOY/LOX events exhibit more variability, with donor-specific enrichments seen in only 3/12 (25%) mosaicisms. Furthermore each shows bias for a different lineage (**Fig. S13**), with LOY enriched in MEPs in BM66, LMPPs in BM702, and HSCs in BM712. The reduced cell-type bias and high recurrence of sex chromosome loss suggest that the functional consequences of LOY may be less impactful and/or more context-specific^33^ compared to subclonal mSVs. By comparison, singleton mSVs do not show cell-type bias across donors (**Fig. S12)**, suggesting that the principal driver for cell type-specific subclonal expansion of these mosaicisms in HSPCs is likely linked to a specific impact of the subclonal mSVs on cell function, rather than their biased acquisition in certain cell-types. Altogether, these data suggest a potentially appreciable functional effect of mSVs, which is cell-type- and locus-specific.

Despite the diversity of genomic loci impacted by mosaicism, we find evidence for recurrent molecular phenotypic effects. We observe substantial overlap in dysregulated genes and pathways across donors (**Fig. 3c, d, e**; **Fig**. **S13**, **S14**, **S15; Table S9, S10, S17**). These recurrent pathways include Ras and JAK/STAT signaling, both of which are commonly associated with clonal hematopoiesis, proliferation, and hematological malignancy^34^. These data directly link subclonal mSVs to commonly observed changes in aging-related pathways in HSPCs. As such, we expand on the cellular context and functional consequences of two particularly interesting examples in the following sections.

### Cell-type-specific consequences of a mosaic inversion

The molecular consequences of mosaic inversions in normal tissues are under-appreciated, reflecting prior difficulties in subclonal inversion discovery^7, 8^. We therefore investigated the effects of the subclonal balanced inversion identified on chromosome Xq12-Xq21.1 (‘Xq-Inv’), seen in 22.6% (19/84) of cells from a 65-year old female donor (BM65; **Fig. 4a**). Analysis of the chromosome X NO profiles^18^ confirms the inversion arose on the active X-homolog (**Fig. S16**), supporting its potential for driving a functional consequence. We refined the inversion breakpoints from the Strand-seq data^35^, and fine-mapped the 10 Mb inversion to chrX:66753519-76960327, with breakpoint confidence intervals of ∼10 kb and ∼18 kb, respectively. While neither breakpoint directly overlaps a gene, they fuse two annotated topologically-associating domains (TADs) (**Fig. 4b**), potentially reorganizing the local gene regulatory environment^36^ in these regions.

**Figure 4:**
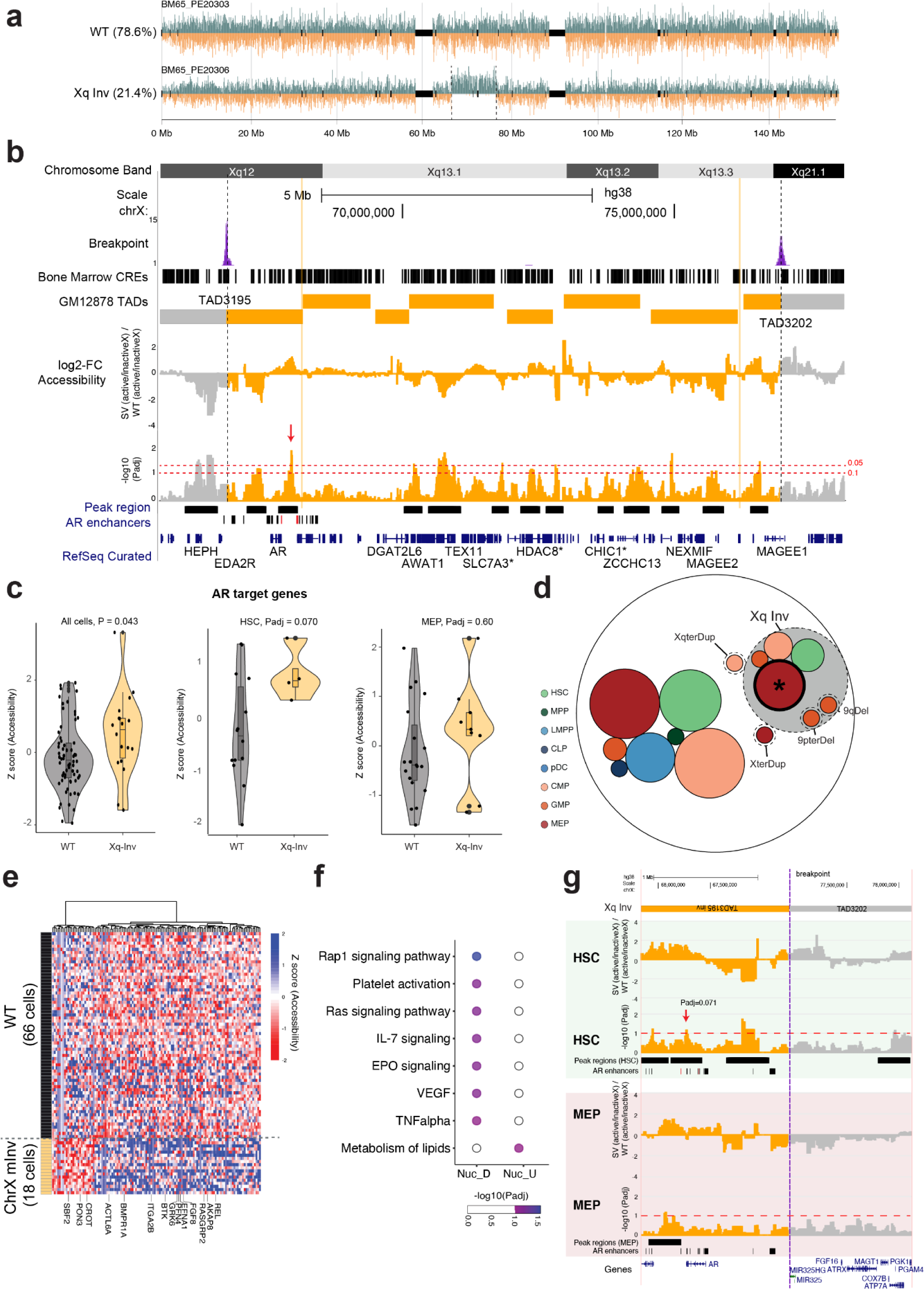
**Somatic mosaic inversion driving MEP-biased cell fate and subclonal expansion of HSPCs through *cis* and *trans* effects. a**) Strand-seq data of X chromosomal homologs from BM65 depicting the unaffected haplotype 2 (also denoted ‘WT’; top) and the Xq-Inv on haplotype 1 (bottom). **b**) Genome browser track showing the confidence interval of inversion breakpoints and the TAD boundaries^74^ around them. Below, NO differences at CREs between Xq-Inv and WT cells are shown as log2 fold changes. Permutation adjusted *P*-values were computed using a sliding window approach as previously described^18^. The most significant signal out of 13 peaks representing patterns of haplotype-specific NO is a region with inferred increased chromatin accessibility, which overlaps with annotated AR enhancers^75^ residing 386 kbp apart from the *AR* gene. Three annotated *AR* enhancers intersecting with the most significant peak are highlighted in red. **c**) Boxplot of NO of known AR target genes, which exhibit an AR-binding motif in their promoter based on MSigDB^76^, in Xq Inv and WT cells in all cell-types (left), HSCs only (middle) and MEPs (right). **d**) Circle-packing plot depicting cell type-resolved mSV landscape in BM65. Dotted lines denote mSVs, and the grey coloured background denotes a measured cell type enrichment. **e**) Heatmap of differential NO (diffNO) genes identified for the Inv subclone, compared to the WT cells, generated after regressing out the contribution of individual cell types. The Y-axis represents single cells analyzed using scTRIP, and diffNO genes are plotted on the X-axis. Change in NO is coloured from red (increased NO) to blue (decreased NO). **f**) Pathways over-represented by the genes with differential NO (FDR 10%). **g**) Cell-type specific analysis of NO differences at CREs between the mSV subclone and WT cells. *P_adj_* values of significant peak regions (FDR<10%) are highlighted. A red arrow indicates the HSC-specific significant peak region containing two AR enhancers, in which we infer increased chromatin accessibility (the two enhancers are highlighted in red in **Fig. S20**).

To investigate the regulatory landscape of the inversion, we interrogated the haplotype-resolved NO profiles at *cis*-regulatory elements (CREs) to infer chromatin accessibility^18^ on each homolog. Using a haplotype-aware sliding window analysis (**Methods**), we normalized NO between the active and inactive X, and compared Xq-Inv cells with those lacking the mSV (WT cells). We identify 13 peak regions with significantly altered NO (10% FDR; **Fig. 4b**), with four (31%) located within one of the disrupted TADs. The most significant peak fell into an intergenic region within a disrupted TAD, and showed decreased NO on the inverted haplotype, indicative for increased chromatin accessibility^18^. This peak is located adjacent to the androgen receptor gene (*AR*), which was previously identified as a clonal hematopoiesis driver by analyzing normal blood for somatic mutations with evidence of positive selection^37^. Closer analysis shows 3 annotated *AR* enhancers fall within this peak of increased chromatin accessibility (**Table S11**), all of which reside in the fused TAD (**Fig. 4b**, **Fig. S16**). These data suggest *AR* as a potential target of gene dysregulation and contributor to subclonal expansion, through disruption of TADs by a balanced inversion. Indeed, androgens are widely used to treat bone marrow failure syndromes owing to their ability to induce increased HSPC proliferation, although the mode of action is incompletely understood^38^.

We performed a genome-wide search for differential gene activity^18^, by comparing the NO of gene bodies between Xq-Inv and WT cells (**Methods**), in order to identify potential downstream effects of the Xq-Inv outside the of affected Xq chromosomal region. We find 123 genes displaying differential NO (**Fig. 4c**, **Table S10**) – all of which reside outside the inversion locus – suggesting a strong *trans* effect of Xq-Inv. Gene-set over-representation analysis (**Methods**) reveals dysregulation of several pathways including Rap1 signaling, platelet activation and metabolism of lipids (10% FDR; **Fig. 4d**; **Table S12**). Several of these pathways, notably, have previously been reported to be modulated by AR activity, such as Ras signaling and erythropoietin (Epo) signaling (**Table S12)**. Epo signaling, in particular, has been shown to contribute to an erythroid-bias of HSCs in association with elevated AR activity^39, 40^. Finally, TF-target enrichment analysis^18^ reveals 3 TFs with differential activity in Xq-Inv cells: EGR1, RUNX1, and IKZF1 - all of which are linked to AR signaling (**Fig. S17**). These data therefore provide independent support that AR activation represents an important functional consequence of the mosaic Xq-Inv.

Notably, the three TFs identified above (EGR1, RUNX1 and IKZF1) also play critical roles in the MEP lineage^41–43^. This suggests AR activation may directly drive the striking enrichment of MEPs seen in the Xq-Inv subclone (**Fig. 4e**). To explore this, we performed a cell-type-resolved analysis of NO in the AR gene-body, revealing elevated AR activity from the rearranged homolog specifically in HSCs with Xq-Inv, but not in MEPs (10% FDR; **Fig. S18**). Moreover, upon testing AR target genes (**Table S13**) we find increased activity in Xq-Inv HSCs, but not Xq-Inv MEPs (10% FDR, **Fig. 4f**, **Fig. S17**) – suggesting HSC-specific AR overactivation as consequence of the mosaic inversion. Consistent with this, Xq-Inv HSCs contain several unique differential NO peaks not found in MEPs (10% FDR), including at 2 AR enhancers (**Fig. 4g**, **Fig. S19)**. These enhancers, which contain binding sites for EGR1, RUNX1 and IKZF1, are more accessible, suggesting cell-type specific enhancer activity, in Xq-Inv HSCs (**Fig. S20**). Finally, where Xq-Inv HSCs show regulatory changes consistent with elevated AR signaling (with 3/4 differential NO genes representing annotated AR targets), the Xq-Inv myeloid cells (CMPs and MEPs) show a more diffuse signal (with 23/105 and 12/55 differential NO genes being AR targets, respectively; **Table S9**, **Fig. S17**). Among the MEP-specific genes, we infer high activity of *RIT1* (FDR 10%, P_adj_=0.0057), a gene whose overexpression has been implicated in clonal hematopoiesis leading to MEP expansion^44^. Altogether, our data strongly imply an increased AR activity in the mosaic subclone which is HSC-specific, and suggest a ‘priming’ role of the Xq-Inv that biases differentiation towards the megakaryocyte-erythroid lineages.

### Stepwise accumulation of mosaic deletions drives HSPC clonal expansion

The data we obtained from BM65 indicate that subclonal mSVs can have a considerable impact on molecular phenotypes that arise in a cell-type specific manner. Yet, how subclonal expansions are facilitated in cells showing more than one co-existing mSV remains unclear. We explored such complex subclonal dynamics in BM712. This 71-year-old male donor shows five distinct subclones, three of which exhibit a cell-type bias (FDR 10%; **Fig. 5a**). Of the 123 cells sequenced, 103 (84%) harbor at least one subclonal mosaicism. The mSVs include two interstitial deletions affecting distinct chromosome 17 loci and three independently-arisen LOYs (**Fig. 5a,b**). We tracked the subclonal evolution^18^ of BM712 using shared, haplotype-resolved mSVs. One subclone (26% CF) shows LOY as the only mSV event and is enriched for HSCs. The four other subclones trace back to a ∼1.2Mb deletion at 17p11.2 (17p-Del), seen in 56% of cells, that was followed by the progressive acquisition of additional mosaicisms, including a ∼500kb deletion at 17q11.2 (17q-Del) and two independent LOYs (**Fig. 5c,d,e**). Using bulk WGS of sorted CD34- cells from BM712, we verified the presence of both subclonal 17q-Del and 17p-Del events, allowing us to refine the 17q-Del breakpoints at high resolution, and to confirm that the mosaicisms are also detectable in mature peripheral blood cells (**Methods; Fig. 5e, Fig. S21**). These data show that HSPCs can sequentially accumulate mSVs that are carried into mature blood cells.

**Figure 5:**
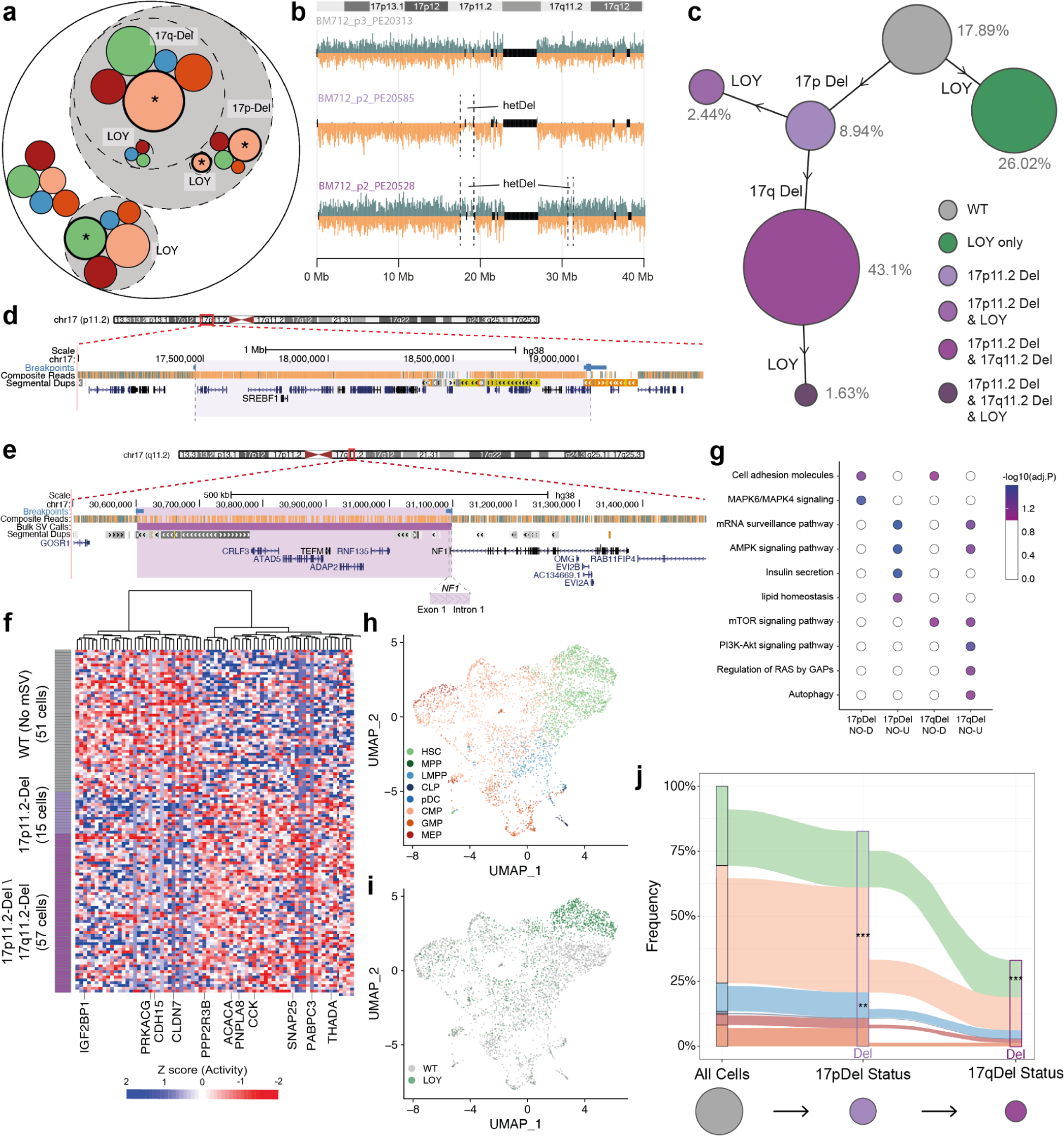
mSV accumulation in a single donor driving clonal expansion. **a**) Circle-packing plot of mSVs found in BM712. **b**) Strand-seq karyograms of unmutated (WT; upper), 17p-Del only (middle) and 17p-Del and 17q-Del (bottom) somatic genotypes. **c**) Bubble hierarchy plot of mSVs identified in BM712. Bubbles are coloured by somatic genotype, and scaled proportional to each subclone’s frequency within the donor. CF is noted beside each bubble, and the distinguishing mSV acquired by each subclone are indicated on the adjoining arm from the parent population. d-e) UCSC genome browser tracks for the 17p-Del (**d**) and 17q-Del (**e**) genomic segments. Tracks for both panels include composite read data and BreakpointR^35^ based breakpoint calls, and highlight relevant genes. In (**e**), the high confidence deletion call from bulk sequencing is also displayed (the inferred VAF from the Delly2 deletion call is 28.5%). **f**) Heatmap of genes showing differential NO between WT, and the union of 17p-Del and 17pq-Del cells. **g**) Pathway over-representation analysis using ConsensusPathDB^67^ for the genes identified in the pairwise comparison of 17p-Del and 17pq-Del subclones to WT cells. Significant pathways were identified with an FDR of 10%. In the x-axis, NO-U and NO-D indicate increased and decreased nucleosome occupancy, respectively. **h**-**i**) UMAP plot of scRNA-seq of CD34+ cells from BM712, with inferred cell-type from reference data^51^ overlaid (**h**) and inferred LOY status (**i**). **j**) Cell-type composition and enrichment analysis for 17p-Del and 17pq-Del subclones in scRNA-seq from BM712.

To explore the functional impact of the initiating mSV, the 17p-Del, we compared gene-body NO of 17p-Del cells with cells bearing a WT chromosome 17 using scNOVA, identifying 76 dysregulated genes (10% FDR; **Fig 5f**). TF-target over-enrichment analysis^18^ shows enrichment for targets of 7 TFs, with the most-significant TF gene being *SREBF1* (*P_adj_*=0.0047) (**Fig. S22**). This gene is hemizygously deleted by the 17p-Del (whereas the other TFs fall outside of the deletion region), suggesting a potential role of *SREBF1* loss in mediating the molecular phenotype of 17p-Del cells and influencing their subsequent expansion (**Fig. 5d**). Protein-protein interaction (PPI) mapping of all 7 dysregulated TFs using STRING^45^ (**Supplementary Methods)** reveals a significant PPI network connecting all TFs (*P*=3.57e-08; **Fig. S22)**, highlighting a functional relationship between these TF pathways (**Supplementary Notes**). Pathway enrichment analysis shows that the PPI network is enriched for components of the MAPK signaling pathway (*P_adj_=*0.0028), which has been previously linked to cell-cycle activation of aging HSCs^46^, and which may thus mediate clonal expansion. Gene-set enrichment analysis of the 76 genes with differential NO further supports MAPK activation in the 17p-Del subclone (**Fig. 5g**), and also identifies dysregulation of lipid homeostasis - a well-known contributor to increased myelopoiesis^47^. Taken together, these data suggest that the 17p-Del mSV triggers increased MAPK activity that likely drives myeloid-biased clonal expansion via hemizygous loss of *SREBF1*.

We next examined the consequences of the second mSV: the 0.49 Mb 17q-Del, which arose in the 17p-Del subclone and expanded to a CF of 43.1%. Our breakpoint mapping shows the deletion results in the hemizygous loss of the *NF1* tumor suppressor via loss of protein-coding exon 1 (**Fig. 5e, Fig. S23, S24)**. Notably, *NF1* exon 1 deletions were previously reported to cause the neurofibromatosis 1 hereditary cancer syndrome^48^. Furthermore, *NF1* has been proposed as a clonal hematopoiesis driver gene based on an SNV analysis, which annotated 175 mosaic SNVs arising within *NF1*.^37^ We find 26/175 (15%) of these SNVs are predicted loss-of-function (pLoF) variants, including 17 stop gains and 9 frameshifts. Additionally, 21/175 (12%) are predicted to affect splicing and thus potentially result in irregular *NF1* transcripts (**Supplementary Notes**). Collectively, these data suggest that *NF1* hemizygous loss could fuel clonal expansion.

To examine the downstream consequences of 17q-Del, we utilized scNOVA to identify dysregulated genes between 17q-Del cells and WT cells. We find 112 dysregulated genes, and pathway over-representation analysis shows altered metabolism and upregulated mTOR signaling in the 17q-Del subclone (**Fig. S25**). Given the known critical role of *NF1* in mTOR signaling^49^, and the role of mTOR signaling in cell proliferation and HSPC differentiation^50^, these findings indicate the 17q-Del results in the hemizygous disruption of *NF1* to induce mTOR dysregulation, potentially leading to accelerated subclonal expansion of the 17q-Del mSV.

To further characterize these subclonal dynamics, we generated 4,109 scRNA-seq libraries from CD34+ cells isolated from BM712 (**Fig. S26**), which we assigned into HSPC cell types using a transcriptome reference of human blood^51^ (**Fig. 5h**). While the resolution of scRNA-seq data is thought to be inadequate for mSV discovery, Strand-seq-derived DNA rearrangement calls can be utilized to perform targeted re-calling of CNAs, allowing transcriptomic analysis of some CNAs across a wide dynamic expression range^18^. Therefore, leveraging the mSV breakpoint assignments from scTRIP, we conducted targeted re-calling of each mosaic CNA (**Methods; Table S14**), which infers 2,571 (63%) 17p-Del, 1,841 (45%) 17q-Del, and 995 (24%) LOY scRNA-seq cells in the dataset. Co-occurrence analysis of these mosaicisms corroborates the intricate subclonal structure of BM712, initially constructed from Strand-seq (**Fig. S27**). Moreover, and also in line with our Strand-seq findings, cell-type enrichment analysis of the scRNA-seq data verify all 3 mosaicisms show significant lineage biases with the 17p-Del enriched for CMPs and LMPPs (P_adj_=2.0e-11; 0.0064; FE test), and both 17q-Del and LOY clones enriched for HSCs (P_adj_=2.6e-14, P_adj_=1.0e-56; FE test; **Fig 5i, j**; **Fig. S26**).

We next performed a transcriptomic analysis of each BM712 mosaic subclone, to obtain a more nuanced view of the molecular phenotypes of the identified subclones. Gene ontology analysis of the differentially expressed genes between HSCs with and without LOY reveals dysregulation of pathways linked with HSC quiescence^52, 53^ (10% FDR; **Table S15, S16)**, potentially explaining the observed HSC enrichment of LOY in this donor, which was not seen for any other LOY in the cohort. Furthermore, consistent with our NO analyses, 17q-Del cells show a highly distinct transcriptional profile compared to WT cells, with differential gene activity seen for 16 pathways (MSigDB Hallmark; T**able S15**; **Table S16**; **Fig. S28**) including those related to HSPC proliferation, differentiation and metabolism. Notably, these pathways include *MYC* and mTOR signaling through *mTORC1* – two known downstream effectors of somatic *NF1* inactivation^49, 54^ – which have been linked to HSC expansion and inhibition of differentiation^55, 56^. Interestingly, we find a significant enrichment of HSCs in 17q-Del cells compared to 17p-Del cells (*P*_adj_=2.1e-05; **Fig. 5j**), potentially mediated through the upregulation *MYC* and/or *mTORC1*^55, 56^. Finally, the 17q-Del subclone shows evidence for an altered DNA damage response, with decreased expression of genes such as *BRCA1*, *BRCA2*, *FANCI* and *BLM* – suggesting that 17q-Del cells might be prone to acquire further somatic mutations. Together, these data suggest that BM712 underwent a stepwise acquisition to a potentially ‘higher-risk’ molecular phenotype; firstly by enabling HSCs to exit quiescence and biasing their differentiation; and secondly, by inducing a proliferative and more HSC-like molecular phenotype which could be more permissive to acquiring further mosaicism.

### Functional effects of mSVs in blood samples based on targeted re-analysis in the UK Biobank

Based on our in-depth characterization of subclonal mSVs found in two donors, we identified target genes that are dysregulated and appear to drive or influence HSPC subclone expansion. To explore whether these findings can be extrapolated to a larger cohort, and test whether they also have implications for circulating blood cells, we interrogated the UK Biobank^57^. The extensive phenotypic data paired with whole-exome sequencing (WES) data for 469,792 blood donors^57^ provide the opportunity to study somatic mutations in relation to blood counts in the normal human population. Focusing on the genes that showed direct functional evidence within the 17p-Del, 17q-Del and Xq-Inv subclones (respectively *NF1*, *SREBF1* and *AR*), we extracted rare (MAF<1%) SNVs and small InDels from the WES data, which we classified based on their potential for functional impact (**Methods**; **Table S18**). Since CNA losses affecting both the 17p-Del and 17q-Del regions have been observed in in UK Biobank samples^2, 58^, we additionally made use of WES-based CNA loss calls recently generated for 200,624 UK Biobank donors^58^ which we analyzed by burden testing (**Methods**). We first concentrated our analyses on the 17p and 17q regions, analyzing variants leading to gene disruption. We find a bimodal somatic variant allele frequency (VAF) distribution for *NF1* and *SREBF1* pLoF SNVs, but not for synonymous variants (**Fig. 6a; Fig. S29**) – indicating pLoF SNVs, but not other SNVs, exhibit mosaicism at these loci. These data hence underpin the link between gene-disrupting mosaicisms affecting *SREBF1* and *NF1* and clonal expansions in normal blood.

**Figure 6:**
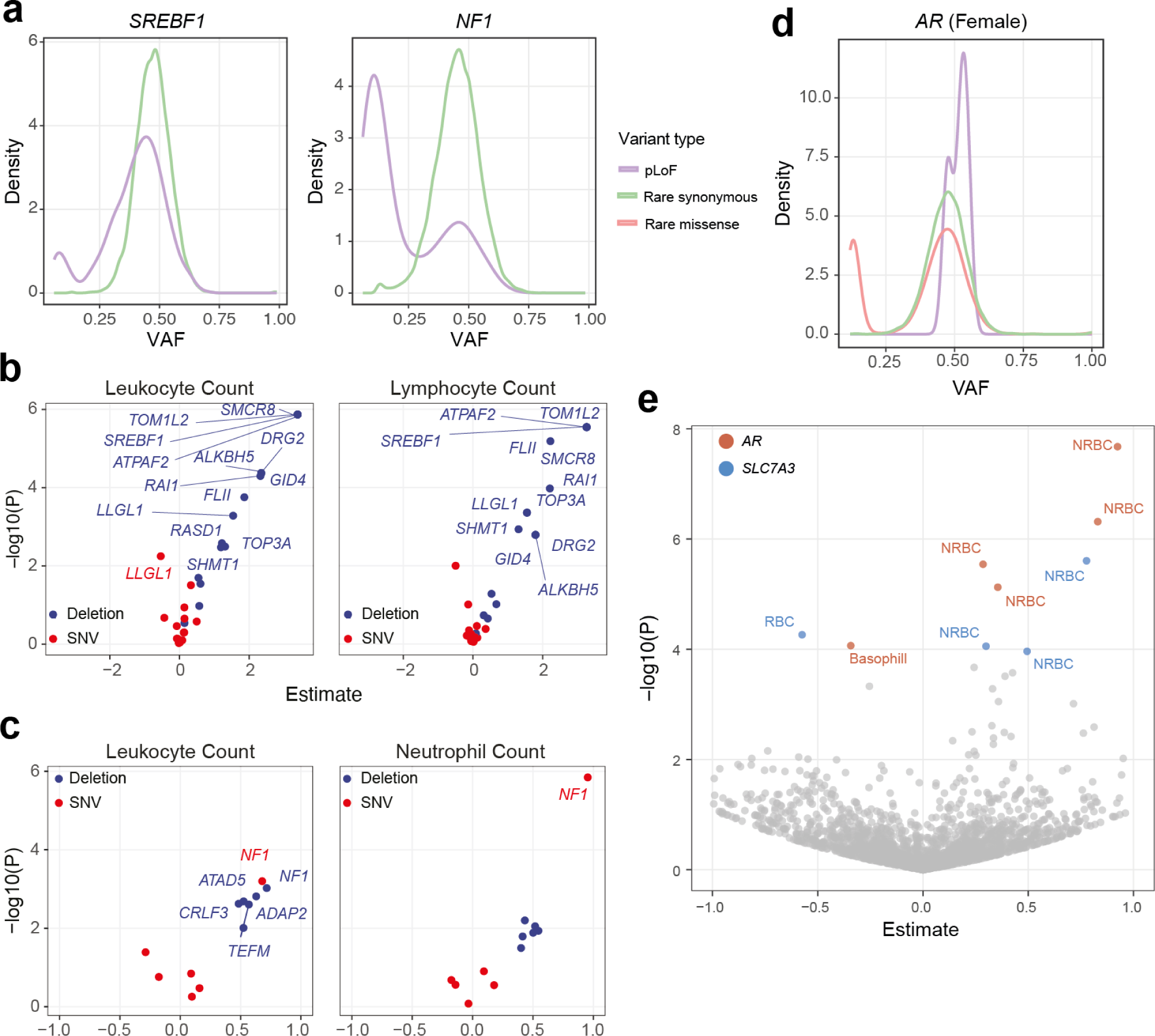
Functional effects of mSVs are supported by informed re-analysis of large-scale bulk dataset. **a**) VAF plot for mutations in left) *SREBF1* and right) *NF1*, separated by mutation type, from the UK Biobank. **b-c**) Volcano plot showing burden test results for 17p-Del (**b**) and 17q-Del (**c**). Genes with *P_adj_*< 0.05 are labeled. Only selected leukocyte count traits are shown. See **Fig. S29** for all blood count traits. Y-axes show nominal *P*-values. **d**) VAF plot for mutations in *AR*, separated by mutation type. See **Fig. S30** for male data. **e**) Volcano plot showing association test results of single rare missense variant at the Xq-Inv locus for all 11 blood count traits (generated from female donors; see **Methods**). The full respective list of missense variants analyzed is available from **Table S18**. Variants with *P_ad_*_j_<0.05 are colored by gene and labeled by trait: NRBC, nucleated red blood cell count; basophil, basophil count. Variants with *P_adj_* >= 0.05 are colored in gray. Y-axis in (**b**), (**c**) and (**e**) depicts nominal *P*-values.

At the *SREBF1* locus, we find both CNA losses and pLOF SNVs are associated with altered blood counts (n=2 losses and n=74 pLOF SNVs; **Table S18**), with this gene being the most significant hit within the 17p-Del region for several categories, including elevated total leukocytes (*P_adj_*=0.00012; loss), elevated lymphocytes (*P_adj_*=0.00013; loss) and elevated nucleated red blood cell count (*P_adj_*=0.039; pLOF) (**Fig. 6b; Fig. S29**). While these findings do not formally rule out potential effects from other genes in this region, they bolster support for our findings that disruption of *SREBF1* can contribute to a cell-type skewing in healthy blood. Delineating the causal relationship will require further study. When repeating the same analysis for all genes in the 17q-Del region, we find losses at 5/6 are associated with elevated total leukocytes – yet, only for the *NF1* locus we find that both loss and pLoF SNVs are significant (*P_adj_*=0.042 for both; **Table S18**; **Fig. 6c; Fig. S29**). This supports the notion that both 17p-Del and 17q-Del contribute to cell-type skewing and potentially clonal expansion within the blood. Interestingly, pLOF in *NF1* alone exhibits a highly significant increase in neutrophil counts (*P_adj_*=0.00019) – which strongly implicates this gene in myeloid-skewed hematopoiesis.

Lastly, we analyzed rare missense SNVs at the Xq-Inv locus (n=5 genes), motivated by prior reports of activating somatic missense mutations at the *AR* locus in cancer^59^ – which we reasoned could potentially mirror the AR activation-based molecular phenotype we observe in the female BM65 donor. In females, we observe a bimodal VAF for missense, but not pLoF, SNVs, which affirms that *AR* missense SNVs, but not *AR* pLoF SNVs, exhibit somatic mosaicism in normal blood (**Fig. 6d; Supplemental Notes**). Furthermore, we find that 5 rare *AR* missense, but not *AR* pLoF SNVs, are associated with altered blood cell counts (*P_adj_*<0.05, for all 5 SNVs; **Fig. 6e**). These fall into exon 1 (n=3), exon 2 (n=1), and exon 4 (n=1), all which also harbor somatic missense SNVs in cancer which on AR function^59^. We observe association with increased nucleated red blood cell count for n=4 missense SNVs (*P_adj_*<0.05, for all 4 SNVs), and decreased basophil cell count for the remaining SNV (*P_adj_*=0.043). These findings strongly support our above analyses, associating activation of *AR* with altered blood cell counts – specifically in the megakaryocyte-erythroid lineages – in a large cohort of healthy blood donors.

## Discussion

Our study provides a systematic cell-type-resolved exploration of distinct mosaicism classes, illuminating the functional impact and cellular context of mSVs in HSPCs for the first time. Earlier studies reporting on mosaic CNAs in the blood highlighted the discovery of subclonal CNAs as a feature of the normal aging population, seen in sample subsets^2, 6, 7, 20, 21^. In comparison, the high mapping resolution of Strand-seq^14^ (**Fig. S31**) identifies mSVs in the majority of samples, spanning the donor age spectrum. Our data imply that mSVs may impact both gene activity and the locus-specific *cis*-regulatory environment of genes in a cell-type-specific manner. Despite arising within distinct loci, we find subclonal mSVs converge on aberrant growth and/or developmental signaling pathways and most frequently associate with the myeloid lineage. This cell-type bias is notable given the commonly observed age-related myeloid bias of HSPCs^30^ and disease pathology involving myeloid lineage cells, including leukemia development^60^. By comparison, LOYs exhibit more variability, with cell-type enrichments differing in each donor.

The close association of SCEs and mSVs suggests that mSV formation is frequently triggered by abnormal DSB repair, either through unequal sister chromatid recombination^61^ or non-homology associated repair processes^22^. Our observations indicate that mSV formation in HPSCs occurs irrespective of age, suggesting a pattern akin to some point mutation processes which show a consistent rate throughout life^62^. However, our data imply that cells harboring mosaicisms are more prone to accumulate additional mSVs, potentially owing to increased propensity to generate or tolerate additional mutations, which could foster cumulative functional effects of mSVs in certain subclones. While longitudinal data would be needed to corroborate this, we note that our data hint at a potential similarity between mSVs and clonal hematopoiesis associated with SNVs, whereby the individuals with multiple mosaic SNVs are considered at higher risk for malignant transformation^63^. Furthermore, newly formed mSVs frequently result in large terminal gains and losses, whereas all clonally expanded mSVs in our dataset represent interstitial events. These large terminal gains and losses may not (or only rarely) reach appreciable CF, perhaps due to the detrimental consequences of autosomal aneuploidy^13^ inhibiting cell proliferation in normal HSPCs. Moreover, these data imply that factors other than an elevated mSV formation rate contribute to mSV subclonal expansion during aging. Depletion or exhaustion of the HSC pool resulting in decreased clonal diversity over time^12^, or changes to the BM microenvironment such as increased inflammation that may favor cells bearing particular large scale mosaicisms, could mediate mSV subclonal expansion during aging.

Prior reports have associated mosaic CNAs in blood with clonal hematopoiesis^2, 7, 20, 21^, an age-related phenomenon where HSPCs contribute to genetically distinct blood cell subpopulations. Our findings imply that subclonal mSVs, seen in 36% of donors over 60 years, commonly impact HSPC function by affecting diverse genomic loci, including genes with known or suspected roles in clonal hematopoiesis^37^. The prevalence of this class of mosaicism implies that the cumulative phenotypic impact of mSVs on specific tissues or organs could potentially at least parallel that of SNVs, a finding that underscores the necessity for future studies in larger cohorts. Given known challenges in detecting mSVs present with a low CF^8^, an important consideration will be the choice of technology for coupling genomic and functional readouts. High-throughput production of Strand-seq libraries in nanoliter vessels may be a promising path forward to enable more large-scale studies, given the unique capabilities of this technology to enable single-cell multiomics of a wide variety of structural variant classes, and at appreciable scale^18, 64^. While Strand-seq could in principle access any dividing cell, post-mitotic tissues as well as mSVs smaller than 200 kb in size (such as mobile element insertions) are not currently amenable to profiling with this technology. Finally, an unaddressed question of interest is whether mSVs may synergise with SNV mosaicism associated with clonal hematopoiesis^4, 5^, a question that was not the focus of this study. We caution, however, that single-cell multiomic methods capturing both mSVs and SNVs sensitively in single cells, and at scale, are presently missing^8^, highlighting an important need for further technology development, before such investigations could be effectively conducted.

In conclusion, the heterogeneous mSV landscape we unveil in the blood compartment has implications for understanding how mSVs impinge on the molecular phenotypes of HSPCs over life. The single-cell multiomic framework applied in this study, which is guided by a novel scMNase-seq-based NO reference for HSPCs, paves the way for systematically linking mSVs to molecular phenotypes related to clonal expansions in diverse normal human tissues.

## Data and materials availability

We have made all genomics data generated in this study (Strand-seq, scMNase-seq, scRNA-seq, bulk WGS) available under the following accession: EGAS00001006567.

## Code availability

Our study has publicly made available a novel scMNase/NO-based classifier for cell-typing Strand-seq libraries in HSPCs. This classifier can be accessed and downloaded from GitHub, to facilitate its use in further studies: https://github.com/jeongdo801/NO_based_HSPC_classifier

## Authorship contributions

Study design: K.G., H.J., A.D.S., J.O.K.; Contribution of BM samples: H.G., D.N., J-C.J, A.A., J.N., J.N., R.E., M.H., A.L., A.H.; Contribution of UCB samples: A.H.; Strand-seq experiments: K.G.; Strand-seq library preparation: B.R., P.H., C.S., E.B.; Plate-based scMNase-seq protocol development: K.G.; Cell-type classification models: H.J., mSV calling: K.G., H.J.; SCE calling: K.G.; scNOVA analysis: H.J., K.G.; WGS based mosaicism calling: K.G., T.R.; scRNA-seq data generation: K.G.; scRNA-seq data analysis K.G., H.J.; Cell type classification and enrichment testing: H.J., K.G.; data interpretation: K.G., H.J., A.D.S., J.O.K; UK biobank analyses: N.X., S.S.; Paper writing: K.G., H.J., A.D.S, J.O.K., with contributions from all authors.

## Acknowledgements

We acknowledge EMBL core facilities and services for support in computing (IT), sequencing (GeneCore) and cell sorting (FACS). We thank Jiaxuan Li from Southern University of Science and Technology for assisting with the analysis of the Pan-Cancer Analysis of Whole Genomes dataset. Principal funding from this work came from an ERC consolidator grant (MOSAIC) to J.O.K. This work was additionally supported by the Health + Life Science Alliance Heidelberg Mannheim which receives state funds approved by the State Parliament of Baden-Württemberg in Germany. N.X. and S.S were supported by the National Natural Science Foundation of China (32200487) and the Center for Computational Science and Engineering at Southern University of Science and Technology.

## Ethics declarations

For samples from the department of Hematology and Oncology, Medical Faculty Mannheim, Heidelberg University, the use of primary human materials for research purposes was approved by the Medical Ethics Committee II of the Medical Faculty Mannheim of the Heidelberg University. The Ethics approval number is 2013-509N-MA. For samples from Ulm University Hospital, collection and investigation was approved by the Internal Review Board (Ethikkomission) at Ulm University (392/16). Healthy samples used in this study were obtained from waste bone fragments obtained from endoprosthetic surgery and cardiovascular surgery. Recruitment was based on availability and written informed consent. The status “healthy” was defined as being negative for HIV, Hepatitis B and C, having a normal blood count and no history or currently active malignancy. For samples from the Department of Medicine V, Hematology, Oncology and Rheumatology, University of Heidelberg, bone marrow samples were harvested from the posterior iliac crest. The studies on aging of bone marrow HSPCs have been approved by the Ethics Committee for Human Subjects at the University Heidelberg. Healthy human subjects were recruited through an announcement published in the Department’s Newsletter for patients and their family. Before donation, healthy subjects were examined and screened by an internist and blood examinations (complete blood count, routine panel of laboratory examinations) were performed to assure their “healthy” status. UCB was collected after informed consent of the mother using the guidelines approved by the Ethics Committee on the use of Human Subjects. All donors provided written informed consent and all interventions were performed in accordance with the Declaration of Helsinki.

## Methods

### Human samples

Healthy donor human umbilical cord blood and bone marrow were obtained either as frozen aliquots of mononuclear cells (MNCs) or freshly isolated from Heidelberg University Hospital, Ulm University Hospital, Mannheim University Hospital, and ATCC (ATCC PCS-800-013™), and were cryopreserved in liquid nitrogen until processing. Strand-seq library generation was initiated from cultures obtained either from freshly isolated or freshly thawed MNCs. For scMNase-seq and 10X scRNA-seq library generation, freshly thawed MNCs were used.

### Statistical testing and multiple test correction

All significance tests used are reported, where applied, in the main text. Multiple test correction was utilized as required, indicated by P_adj_, with a false discovery rate (FDR) of 10%. Our targeted analysis of UK Biobank data employed a more stringent significance threshold of *P_adj_* < 0.05. FE test is used as an abbreviation for Fisher’s exact test.

### HSPC culturing and Strand-seq

UCB samples were obtained from Heidelberg University Hospital. BM from healthy donors was isolated either from donor BM aspirations (N=2), discarded pelvis from hip replacement surgeries (N=6), or sternum removed during routine heart surgeries (N=8) (**Table S1**). Cells were stained on ice in the dark for 30 mins with CD34-APC (clone 581; Biolegend; 1:100), CD38-PE/Cy7 (clone HB7; eBioscience), CD45Ra-FITC (clone HI100; eBioscience), CD90-PE (clone 5E10; eBioscience), and LIVE/DEAD Fixable Near-IR Dead Cell Stain (Thermofisher). Single, viable, CD34+ cells (gating as per **Fig. S1**) were FACS-sorted (BD FACSMelody, 100 μM nozzle, single-cell mode) directly into ice-cold complete medium (Stemspan serum-free expansion medium (SFEM) supplemented with 100 ng/ml SCF and Flt3 (Stem Cell Technologies) and 20 ng/ml IL-3, IL-6, G-CSF and TPO (Stem Cell Technologies). Cells were seeded into Corning® Costar® Ultra-Low Attachment 96-well plates (Sigma-Aldrich) at a density of 1-2x10^5^ cells/ml and cultured for 42h in the presence of BrdU. BrdU-containing nuclei were sorted into 96-well plates and subject to Strand-seq library preparation. All Strand-seq libraries were generated using a Biomek FXP liquid handling robotic system, as previously described^16, 23^. Libraries were sequenced on an Illumina NextSeq 500 sequencing platform (MID-mode, 75 base pair paired-end sequencing protocol). Somatic structural variant calling and NO profiling was pursued using the scTRIP^14^ and scNOVA^18^ workflows, as previously described.

### Single-cell micrococcal nuclease sequencing (scMNase-seq)

HSPCs from a healthy BM donor were obtained from ATCC (ATCC PCS-800-013™), whereas normal UCB samples were obtained from Heidelberg University Hospital. Frozen MNCs were thawed and stained as per **Table S5** with antibodies outlined in **Table S19** in order to distinguish the 8 distinct HSPC populations outlined in **Fig. S8.** Single, viable HSPCs (gating strategy **Fig. S8**) were index-sorted using a BD FACSAria™ Fusion Cell Sorter (100 μM nozzle, single-cell mode) into 96-well plates containing 5 μl modified freeze buffer (0.1% NP-40, 7.5. % DMSO, 42.5 % 2X Profreeze-CDM (Lonza) in PBS) and frozen. ScMNase-seq libraries were generated from sorted, frozen single cells as per Strand-seq library preparation^23^, with the following modification: the Hoechst/UV treatment step was omitted (with scMNase-seq requiring no BrdU incorporation). Following single cell sequencing, each cell had an average sequencing coverage of 613,483 uniquely mapped fragments.

### Building NO reference set cell type classifiers

The scNOVA framework enables cell-typing in single cell NO datasets based on supervised machine learning classification approach. While previously applied to distinguish cell lines from distinct tissues^18^, here we sought to apply scNOVA to classify closely related HSPC cell-lineages, based on generating single-cell NO reference profiles from FACS-sorted HSPCs subjected to scMNase-seq. We constructed distinct NO based classifiers for cells derived from each source. We index-sorted both the BM-derived and UCB-derived CD34+ cells from eight HSPC cell types using previously-defined immunophenotypes^28^ (**Fig. S8**, **Table S5**). Indexed scMNase-seq libraries were used as the ground-truth input for cell type classifiers. In the case of BM HSPCs, gene body NO profiles were extracted for 305 high quality single cells and normalized by library size to obtain reads per million (RPM). These normalised values were log2-transformed and standardised, before being subject to supervised partial linear square discriminant analysis^65^ (PLS-DA) to (1) identify informative feature sets, and subsequently (2) build a classification model. To identify informative feature (gene) sets for each cell type, we used variable autosomal genes to build an X-matrix (305 cells x 18,851 genes) and Y-matrix (305 cells x 8 cell types). These X and Y variables were passed to the PLS-DA, which output variance importance in the projection (VIP) for each feature. The 1,904 genes with a VIP value >90 % of the null distribution from the permutation test were retained for the second stage of feature selection. In the second feature selection stage, a second X-matrix (305 cells x 1,904 genes) and Y-matrix (305 cells x 1 cell type; with cell type in this case being binary information for each cell either belonging to that cell type (1) or not (0), based on FACS indexes) were passed to the PLS-DA, and features with a VIP value >95 % of the null distribution from permutation test were retained. This process was repeated for each cell type, resulting in a final informative feature set of 819 genes for BM HSPCs. We repeated these steps for 175 high quality single-cells obtained from UCB HSPCs, which resulted in 899 genes as significant feature sets for cell-type classification. Our study has openly released this classifier (see Code Availability) to facilitate its use in other research studies.

### Cell type enrichment test for donor specific mSVs

We devised cell-type enrichment tests for each of the identified subclones exhibiting specific mSVs, using a control group consisting of all individuals over the age of 60 who were not affected by mSVs. We performed a binomial test to determine if the number of cells in a particular cell type within the subclone was greater than expected, based on the single cell based cell type composition of the control group. We then calculated permutation-based adjusted p-values for each subclonal mSV by randomly sampling the same number of HSPCs from the entire single-cell population 100,000 times and tallying the number of cells from given cell types in question belonging to that subclone.

### Single-cell multiomic analysis of differential gene activities in HSPC subclones

Differentially active genes in subclones affected by mSVs were identified using scNOVA^18^. We used scNOVA’s infer altered gene activity module, using the PLS-DA option, which is recommended for the investigation of low CF subclones^18^. To regress-out cell-type effects in the identification of differential gene activity, we considered predicted cell-type for each single cell as a confounding factor when we executed scNOVA’s infer altered gene activity module. Genes within the deleted region were masked from this analysis, given that the correlation of CNAs with the expression of the affected genes does not always directly correlate^66^. Genes with significantly altered gene activity (10% FDR) were subjected to gene set over-representation analysis using ConsensusPathDB^67^. Significantly overrepresented pathways (10% FDR) were visualized as dot plots. When comparing 17p-cells and WT cells in BM712 in **Fig. 5f**, we considered all cells carrying the 17p-Del including those harboring other mosaicisms in addition to 17p-Del as ’17p-cells’.

### Investigation of potential *cis*-effects of a balanced inversion

To investigate the local effects of Xq-Inv in BM65, we employed scNOVA^18^. We utilized a sliding window approach previously used to uncover *cis-*effects of balanced somatic DNA rearrangements in leukemia in a haplotype-aware manner^18^. We focused on the Xq-Inv-affected segment, including both of its rearranged TADs. We firstly defined CREs based on a prior study utilizing ATAC-seq in HSPCs^28^. We used a sliding window (300kb in size, moving 10kb each)^18^ analyzing CREs along chromosome X, to infer chromosome-wide haplotype-specific NO for the mSV subclone and WT cells, which is predictive for chromatin accessibility^18^. For each sliding window, haplotype-specific NO values at CREs from the mSV subclone (NO in the active X chromosome / NO in the inactive X) and WT cells (NO in the active X / NO in the inactive X) were compared using likelihood ratio tests to obtain nominal *P*-values *[P real]*. As a multiple testing correction to control the type I error, we performed a permutation test by randomly shuffling genotype labels of each single cell (mSV or WT), in the single-cell RPM matrix 1000 times. For each permutation we performed likelihood ratio tests to compare NO between randomly shuffled mSV subclones and WT cells. We computed the number of incidences we observed the same, or a lower, *P*-value than *[P real]* from 1000 permutations, and divided this value by the number of trials (N=1000) to estimate the permutation-adjusted *P*-value. Sliding windows with permutation-adjusted P-value lower than 0.1 were identified as significantly altered windows, and were assigned to the nearest genes within the same topologically associated domain (TAD) boundaries.

### Single-cell RNA sequencing (scRNA-seq)

Bone marrow MNCs were thawed and stained as previously described, with the following antibodies: CD34-AF488 (clone 561; BioLegend; 1:20), CD38-PE/Cy7 (clone HB7; eBioscience; 1:100). Cells were washed and resuspended as above, and stained for 5 mins with DAPI prior to sorting. The gating strategy as described in **Fig. S1** was used to sort CD34+ cells and CD34- cells respectively into ice cold 0.04 % BSA in PBS using a BD FACSMelody cell sorter. For each donor, 2 samples were prepared: one sample of CD34+ cells and one sample a 50:50 mixture of CD34+ and CD34- cells. scRNA-seq libraries for each sample were generated as per the standard 10x Genomics Chromium 3′ (v.3.1 chemistry) protocol. Completed libraries were sequenced on a NextSeq5000 sequencer (HIGH-mode, 75 bp paired-ends).

### scRNA-seq data processing, unsupervised clustering and cell type annotation

Transcripts were aligned to the human reference genome (GRCh38) and quantified into count matrices using Cellranger mkfastq and count workflows (10X Genomics, V 3.1.0, default parameters). Seurat^68^ (V 3.2.2) was used for QC of single cells and unbiased clustering of the data. Briefly, cells with < 1000 UMIs and cells with > 6 % of mitochondrial reads were removed as ‘low quality’. Normalisation, feature selection, scaling and dimensionality reduction were carried out using default settings. To annotate cell-types, previously reported scRNA-seq data from HSPCs^51^ were used as a reference for cell-type labeling using SingleR^69^. Differential expression analysis to identify cluster-/genotype-specific marker genes was carried out using the FindMarkers function from Seurat.

### Targeted CNA re-calling in scRNA-seq data

Single cell RNA-seq data was normalized to counts per million (CPM) and transformed into log2(CPM/10+1) using Seurat^68^ (V3.2.2). These values were then subject to targeted CNA recalling using the CONICSmat package^70^, as described previously^18^. For the analysis of donor BM712, all three subclonal mosaicisms were investigated: 17p-Del, 17q-Del, and LOY. By default, the CONICSmat ‘plotChrEnrichment’ function considers genomic loci which have more than 100 expressed genes for CNA discovery. Since we performed targeted re-calling variants which were previously identified in the donor cells, and given the small size of the CNAs in question, regions with 5 or more expressed genes were considered in this analysis. The number of expressed genes detected per mSV were as follows: 17p-Del: 24 genes, 17q-Del: 5 genes, LOY: 17 genes (**Table S14**). To profile the mSV regions with expressed genes, CONICSmat generates distributions of average expression levels across single cells in the given regions, and then fits 1-component and 2-component mixture models to these distributions. It further compares the likelihood ratios of being 1-component (unimodal; i.e. absence of CNAs) and 2-component (bimodal; i.e. presence of CNAs), to determine the most-likely state in those regions based on the Bayesian information criterion (BIC). Candidate CNA regions identified as likely to be bimodal within a 1 % FDR criterion (based on a Chi-squared likelihood ratio test) were considered further for downstream analysis. Once the region was inferred to have bimodality, the posterior probability for each single cell to belong to the normal clone or CNA subclone was calculated. A posterior probability cutoff of 0.8 was used to assign single cells into one of the two clones. This analysis was repeated for each subclonal mosaicism event.

### Construction of genome-wide SCE maps in single cells and locus-specific SCE enrichment

We constructed genome-wide maps of SCEs in each single cell by subjecting the Strand-seq data of single cells to the scTRIP computational pipeline^14^, followed by manual inspection and curation of each call yielding SCE positional coordinates for each cell. These coordinates were padded by 1 bp upstream and downstream. The human reference genome (GRCh38) was divided into 500 kb bins using the bedtools makewindows command^71^, and overlaps between these 500 kb bins and our SCE callset were generated using bedtools intersect; giving the number of times each bin is hit by an SCE. A genomic bin was considered to be hit if the majority of an SCE confidence interval fell within that bin, and each SCE was only counted in a single bin. To compute significance of the calculated SCE counts per bin, the count data per bin genome-wide was then fit to a negative binomial distribution using the fitdist function from fitdistrplus^72^, and p-values were calculated using the qnbinom function (with size=1.2506716, mu=0.4823156), with Benjamini-Hochberg correction. To compute overlap of mSV breakpoints with SCEs, we considered 200kb-sized breakpoint regions (reported breakpoints +/- 100 kb).

### Bulk DNA isolation and mSV breakpoint refinement in whole genome sequencing data

Bulk genomic DNA was isolated from CD34- cells (viable cells from the donors which were not put into culture to be used for Strand-seq library preparation) using the QIAamp DNA Blood Maxi Kit as per the manufacturer’s instructions. Samples were sequenced using a NextSeq 5000 (HIGH mode, 75 bp paired end). Raw whole genome sequencing reads were aligned to the human reference genome, sorted, marked for duplicates and indexed. Structural variants were called using Delly2 (default parameters), combining split read, paired-end and read depth analysis^73^. Unfiltered mSV calls were compared to our mSV callset. Since split read analysis failed to identify the precise breakpoints of the 17p-Del which resides in a repeat-rich area on chromosome 17, we generated a single, directional composite bam file^35^ of the region based on our Strand-seq data to allow for 17p-Del breakpoint refinement with BreakpointR^35^.

### UK Biobank analysis

*Data collection.* The UK Biobank is a large-scale population database of approximately half a million participants aged 40-69 years in the UK. For SNVs and INDELs, we used the population level exome OQFE variants for 469,792 individuals (UK Biobank field ID 23157). For autosomal large deletions, we used CNA loss calls on WES data that were recently generated by subjecting 200,624 individuals from the UK Biobank to the CNest copy-number caller^58^. We considered CNA calls >1 kb. Additionally, we obtained phenotypic data for 11 blood count traits (UK Biobank category ID 100081), containing count for white blood cells, basophils, eosinophils, monocytes, neutrophils, lymphocytes, red blood cells, nucleated red blood cells, platelets, reticulocytes and high light scatter reticulocytes. This research was conducted under the application number 83497. The UK Biobank has ethics approval from the North West Multi-centre Research Ethics Committee (21/NW/0157).

#### Variant annotation

We annotated SNVs/INDELs from WES data using Variant Effect Predictor (VEP v1.0.3) with Loss-Of-Function Transcript Effect Estimator (LOFTEE v0.3-beta) plugin. Variant annotation was performed using Hail v.0.2. According to annotation results, we grouped variants into rare loss of function variants (“high confidence” identified by LOFTEE with a minor allele frequency (MAF) < 1%) and rare missense variants (missense variants annotated by VEP with MAF < 1% in UK Biobank cohort). In the case of CNA losses, we considered deletions overlapping coding exons with MAF < 1%.

#### Association testing

The blood count data were rank normalized using the ‘RNOmni’ package in R. Linear regression models (blood count ∼ genotype + covariates) were used to assess the association between three loci of interests (17p-Del, 17q-Del and Xq-Inv) and blood count adjusted for several covariates including age, sex, and the first five principal components derived from genotype arrays. For all genes at the respective 17p and 17q loci, we used gene rare pLoF burden and rare large CNA loss burden as genotype in the regression model. For all genes at the X chromosomal locus of interest, we used gene burden for rare pLoF variants and rare missense variants in the model. Moreover, since missense variants can have distinct functional impact, we also performed single variant association analysis for rare missense mutations at the Xq-Inv locus by sex. The volcano plot in **Fig. 6e** presents nominal P-values derived solely from female donors only, generated since we made the observation of sex-biased VAF distributions at the *AR* locus in UK Biobank samples. For all data, see **Fig. S30**. A minimum of three individuals with relevant variants was required for association tests of a given gene, with the exception of the 17p-Del CNA seen in only two UK Biobank donors based on WES. *P*-values were obtained using the Wald test and the Benjamini and Hochberg method was used to correct for multiple hypothesis testing.

## References

1. Martincorena, I. et al. Tumor evolution. High burden and pervasive positive selection of somatic mutations in normal human skin. Science 348, 880–886 (2015).

2. Loh, P.-R. et al. Insights into clonal haematopoiesis from 8,342 mosaic chromosomal alterations. Nature 559, 350–355 (2018).

3. Lee-Six, H. et al. Population dynamics of normal human blood inferred from somatic mutations. Nature 561, 473–478 (2018).

4. Jaiswal, S. et al. Age-related clonal hematopoiesis associated with adverse outcomes. N. Engl. J. Med. 371, 2488–2498 (2014).

5. Genovese, G. et al. Clonal hematopoiesis and blood-cancer risk inferred from blood DNA sequence. N. Engl. J. Med. 371, 2477–2487 (2014).

6. Kakiuchi, N. & Ogawa, S. Clonal expansion in non-cancer tissues. Nat. Rev. Cancer 21, 239–256 (2021).

7. Forsberg, L. A., Gisselsson, D. & Dumanski, J. P. Mosaicism in health and disease - clones picking up speed. Nat. Rev. Genet. 18, 128–142 (2017).

8. Cosenza, M. R., Rodriguez-Martin, B. & Korbel, J. O. Structural Variation in Cancer: Role, Prevalence, and Mechanisms. Annu. Rev. Genomics Hum. Genet. (2022) doi:10.1146/annurev-genom-120121-101149.

9. ICGC/TCGA Pan-Cancer Analysis of Whole Genomes Consortium. Pan-cancer analysis of whole genomes. Nature 578, 82–93 (2020).

10. Sano, S. et al. Hematopoietic loss of Y chromosome leads to cardiac fibrosis and heart failure mortality. Science 377, 292–297 (2022).

11. Jaiswal, S. et al. Clonal Hematopoiesis and Risk of Atherosclerotic Cardiovascular Disease. N. Engl. J. Med. 377, 111–121 (2017).

12. Mitchell, E. et al. Clonal dynamics of haematopoiesis across the human lifespan. Nature 606, 343–350 (2022).

13. Tang, Y.-C. & Amon, A. Gene copy-number alterations: a cost-benefit analysis. Cell 152, 394–405 (2013).

14. Sanders, A. D. et al. Single-cell analysis of structural variations and complex rearrangements with tri-channel processing. Nat. Biotechnol. 38, 343–354 (2020).

15. Gawad, C., Koh, W. & Quake, S. R. Single-cell genome sequencing: current state of the science. Nat. Rev. Genet. 17, 175–188 (2016).

16. Falconer, E. et al. DNA template strand sequencing of single-cells maps genomic rearrangements at high resolution. Nat. Methods 9, 1107–1112 (2012).

17. Porubsky, D. et al. Recurrent inversion polymorphisms in humans associate with genetic instability and genomic disorders. Cell (2022) doi:10.1016/j.cell.2022.04.017.

18. Jeong, H. et al. Functional analysis of structural variants in single cells using Strand-seq. Nat. Biotechnol. (2022) doi:10.1038/s41587-022-01551-4.

19. Lai, B. et al. Principles of nucleosome organization revealed by single-cell micrococcal nuclease sequencing. Nature 562, 281–285 (2018).

20. Forsberg, L. A. et al. Age-related somatic structural changes in the nuclear genome of human blood cells. Am. J. Hum. Genet. 90, 217–228 (2012).

21. Jacobs, K. B. et al. Detectable clonal mosaicism and its relationship to aging and cancer. Nat. Genet. 44, 651–658 (2012).

22. Liu, P., Carvalho, C. M. B., Hastings, P. J. & Lupski, J. R. Mechanisms for recurrent and complex human genomic rearrangements. Curr. Opin. Genet. Dev. 22, 211–220 (2012).

23. Sanders, A. D., Falconer, E., Hills, M., Spierings, D. C. J. & Lansdorp, P. M. Single-cell template strand sequencing by Strand-seq enables the characterization of individual homologs. Nat. Protoc. 12, 1151–1176 (2017).

24. Glover, T. W. & Stein, C. K. Induction of sister chromatid exchanges at common fragile sites. Am. J. Hum. Genet. 41, 882–890 (1987).

25. Ye, F., Huang, W. & Guo, G. Studying hematopoiesis using single-cell technologies. J. Hematol. Oncol. 10, 27 (2017).

26. Hennrich, M. L. et al. Cell-specific proteome analyses of human bone marrow reveal molecular features of age-dependent functional decline. Nat. Commun. 9, 4004 (2018).

27. Chen, X. et al. Bone Marrow Myeloid Cells Regulate Myeloid-Biased Hematopoietic Stem Cells via a Histamine-Dependent Feedback Loop. Cell Stem Cell 21, 747–760.e7 (2017).

28. Corces, M. R. et al. Lineage-specific and single-cell chromatin accessibility charts human hematopoiesis and leukemia evolution. Nat. Genet. 48, 1193–1203 (2016).

29. Bunis, D. G. et al. Single-Cell Mapping of Progressive Fetal-to-Adult Transition in Human Naive T Cells. Cell Rep. 34, 108573 (2021).

30. Pang, W. W. et al. Human bone marrow hematopoietic stem cells are increased in frequency and myeloid-biased with age. Proc. Natl. Acad. Sci. U. S. A. 108, 20012–20017 (2011).

31. Amoah, A. et al. Aging of human hematopoietic stem cells is linked to changes in Cdc42 activity. Haematologica 107, 393–402 (2022).

32. Ichii, M., Oritani, K. & Kanakura, Y. Early B lymphocyte development: Similarities and differences in human and mouse. World J. Stem Cells 6, 421–431 (2014).

33. Dumanski, J. P. et al. Immune cells lacking Y chromosome show dysregulation of autosomal gene expression. Cell. Mol. Life Sci. 78, 4019–4033 (2021).

34. van Zeventer, I. A. et al. Evolutionary landscape of clonal hematopoiesis in 3,359 individuals from the general population. Cancer Cell (2023) doi:10.1016/j.ccell.2023.04.006.

35. Porubsky, D. et al. breakpointR: an R/Bioconductor package to localize strand state changes in Strand-seq data. Bioinformatics 36, 1260–1261 (2020).

36. Weischenfeldt, J. et al. Pan-cancer analysis of somatic copy-number alterations implicates IRS4 and IGF2 in enhancer hijacking. Nat. Genet. 49, 65–74 (2017).

37. Pich, O., Reyes-Salazar, I., Gonzalez-Perez, A. & Lopez-Bigas, N. Discovering the drivers of clonal hematopoiesis. Nat. Commun. 13, 4267 (2022).

38. Zhang, Q.-S. et al. Oxymetholone therapy of fanconi anemia suppresses osteopontin transcription and induces hematopoietic stem cell cycling. Stem Cell Reports 4, 90–102 (2015).

39. McManus, J. F. et al. Androgens stimulate erythropoiesis through the DNA-binding activity of the androgen receptor in non-hematopoietic cells. Eur. J. Haematol. 105, 247–254 (2020).

40. Grover, A. et al. Erythropoietin guides multipotent hematopoietic progenitor cells toward an erythroid fate. J. Exp. Med. 211, 181–188 (2014).

41. Behrens, K. et al. Runx1 downregulates stem cell and megakaryocytic transcription programs that support niche interactions. Blood 127, 3369–3381 (2016).

42. Yoshida, T., Ng, S. Y.-M., Zuniga-Pflucker, J. C. & Georgopoulos, K. Early hematopoietic lineage restrictions directed by Ikaros. Nat. Immunol. 7, 382–391 (2006).

43. Desterke, C., Bennaceur-Griscelli, A. & Turhan, A. G. EGR1 dysregulation defines an inflammatory and leukemic program in cell trajectory of human-aged hematopoietic stem cells (HSC). Stem Cell Res. Ther. 12, 419 (2021).

44. Chen, S. et al. Impaired Proteolysis of Noncanonical RAS Proteins Drives Clonal Hematopoietic Transformation. Cancer Discov. 12, 2434–2453 (2022).

45. Szklarczyk, D. et al. The STRING database in 2021: customizable protein-protein networks, and functional characterization of user-uploaded gene/measurement sets. Nucleic Acids Res. 49, D605–D612 (2021).

46. Ito, K. et al. Reactive oxygen species act through p38 MAPK to limit the lifespan of hematopoietic stem cells. Nat. Med. 12, 446–451 (2006).

47. Dragoljevic, D., Westerterp, M., Veiga, C. B., Nagareddy, P. & Murphy, A. J. Disordered haematopoiesis and cardiovascular disease: a focus on myelopoiesis. Clin. Sci. 132, 1889–1899 (2018).

48. Imbard, A. et al. NF1 single and multi-exons copy number variations in neurofibromatosis type 1. J. Hum. Genet. 60, 221–224 (2015).

49. Johannessen, C. M. et al. The NF1 tumor suppressor critically regulates TSC2 and mTOR. Proc. Natl. Acad. Sci. U. S. A. 102, 8573–8578 (2005).

50. Zou, Z., Tao, T., Li, H. & Zhu, X. mTOR signaling pathway and mTOR inhibitors in cancer: progress and challenges. Cell Biosci. 10, 31 (2020).

51. Xie, X. et al. Single-cell transcriptomic landscape of human blood cells. Natl Sci Rev 8, nwaa180 (2021).

52. Singh, S. K. et al. Id1 Ablation Protects Hematopoietic Stem Cells from Stress-Induced Exhaustion and Aging. Cell Stem Cell 23, 252–265.e8 (2018).

53. Kovtonyuk, L. V. et al. Hematopoietic Stem Cells Increase Quiescence during Aging. Blood 134, 2484 (2019).

54. Zhang, P. et al. Chromatin regulator Asxl1 loss and Nf1 haploinsufficiency cooperate to accelerate myeloid malignancy. J. Clin. Invest. 128, 5383–5398 (2018).

55. Laurenti, E. et al. Hematopoietic stem cell function and survival depend on c-Myc and N-Myc activity. Cell Stem Cell 3, 611–624 (2008).

56. Fernandes, H., Moura, J. & Carvalho, E. mTOR Signaling as a Regulator of Hematopoietic Stem Cell Fate. Stem Cell Rev Rep 17, 1312–1322 (2021).

57. Sudlow, C. et al. UK biobank: an open access resource for identifying the causes of a wide range of complex diseases of middle and old age. PLoS Med. 12, e1001779 (2015).

58. Fitzgerald, T. & Birney, E. CNest: A novel copy number association discovery method uncovers 862 new associations from 200,629 whole-exome sequence datasets in the UK Biobank. Cell Genom 2, 100167 (2022).

59. Gottlieb, B., Beitel, L. K., Nadarajah, A., Paliouras, M. & Trifiro, M. The androgen receptor gene mutations database: 2012 update. Hum. Mutat. 33, 887–894 (2012).

60. Bowman, R. L., Busque, L. & Levine, R. L. Clonal Hematopoiesis and Evolution to Hematopoietic Malignancies. Cell Stem Cell 22, 157–170 (2018).

61. Johnson, R. D. & Jasin, M. Sister chromatid gene conversion is a prominent double-strand break repair pathway in mammalian cells. EMBO J. 19, 3398–3407 (2000).

62. Alexandrov, L. B. et al. Clock-like mutational processes in human somatic cells. Nat. Genet. 47, 1402–1407 (2015).

63. Weeks Lachelle D., et al. Prediction of Risk for Myeloid Malignancy in Clonal Hematopoiesis. NEJM Evidence 2, EVIDoa2200310 (2023).

64. Hanlon, V. C. T. et al. Construction of Strand-seq libraries in open nanoliter arrays. Cell Rep Methods 2, 100150 (2022).

65. Boulesteix, A.-L. & Strimmer, K. Partial least squares: a versatile tool for the analysis of high-dimensional genomic data. Brief. Bioinform. 8, 32–44 (2007).

66. Roszik, J. et al. Somatic Copy Number Alterations at Oncogenic Loci Show Diverse Correlations with Gene Expression. Sci. Rep. 6, 19649 (2016).

67. Kamburov, A. & Herwig, R. ConsensusPathDB 2022: molecular interactions update as aresource for network biology. Nucleic Acids Res. 50, D587–D595 (2022).

68. Stuart, T. et al. Comprehensive Integration of Single-Cell Data. Cell 177, 1888–1902.e21 (2019).

69. Aran, D. et al. Reference-based analysis of lung single-cell sequencing reveals a transitional profibrotic macrophage. Nat. Immunol. 20, 163–172 (2019).

70. Müller, S., Cho, A., Liu, S. J., Lim, D. A. & Diaz, A. CONICS integrates scRNA-seq with DNA sequencing to map gene expression to tumor sub-clones. Bioinformatics 34, 3217–3219 (2018).

71. Quinlan, A. R. & Hall, I. M. BEDTools: a flexible suite of utilities for comparing genomic features. Bioinformatics 26, 841–842 (2010).

72. Delignette-Muller, M. L. & Dutang, C. fitdistrplus: An R Package for Fitting Distributions. J. Stat. Softw. 64, 1–34 (2015).

73. Rausch, T. et al. DELLY: structural variant discovery by integrated paired-end and split-read analysis. Bioinformatics 28, i333–i339 (2012).

74. Rao, S. S. P. et al. A 3D map of the human genome at kilobase resolution reveals principles of chromatin looping. Cell 159, 1665–1680 (2014).

75. Fishilevich, S. et al. GeneHancer: genome-wide integration of enhancers and target genes in GeneCards. Database 2017, (2017).

76. Liberzon, A. et al. Molecular signatures database (MSigDB) 3.0. Bioinformatics 27, 1739–1740 (2011).

