## Supplemental Data File 1 for "Cell type-specific consequences of mosaic structural variants in hematopoietic stem and progenitor cells"

### Supplemental Data File1

Strand-seq plots for the Singleton SVs

**Supplemental Data File1 : Strand-seq plots for the singleton SVs which shows the *de novo* mSV formation in HSPCs.** In total 32 singleton SV events were identified from the 1133 single-cell libraries included in this study, among which 3 were depicted in the Fig. 1c, and the rest of 29 were depicted in this file. terDel, terminal deletion; terDup, terminal duplication; intDel, interstitial deletion; LOY, loss of Y chromosome; Chr, chromosome.

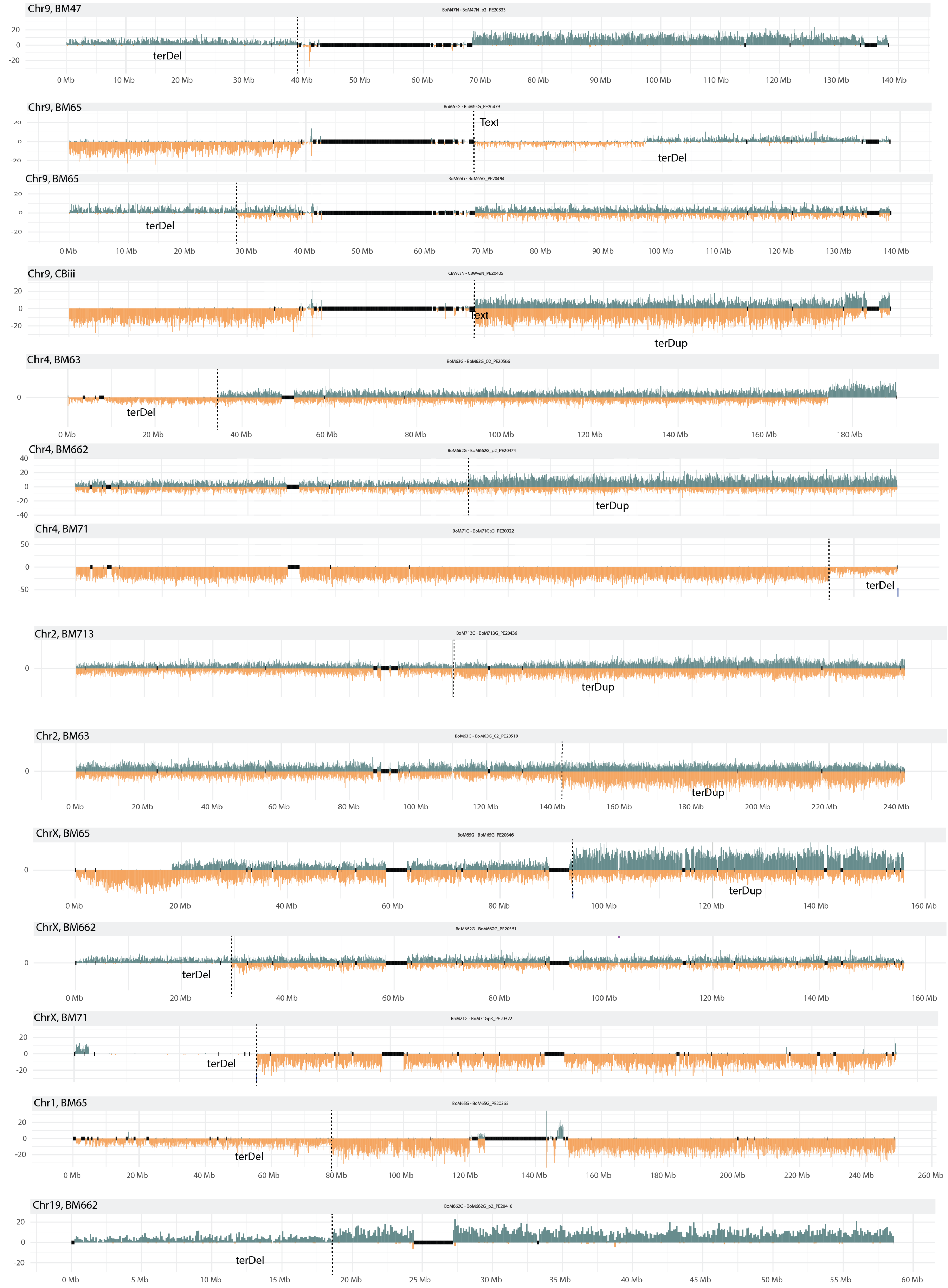

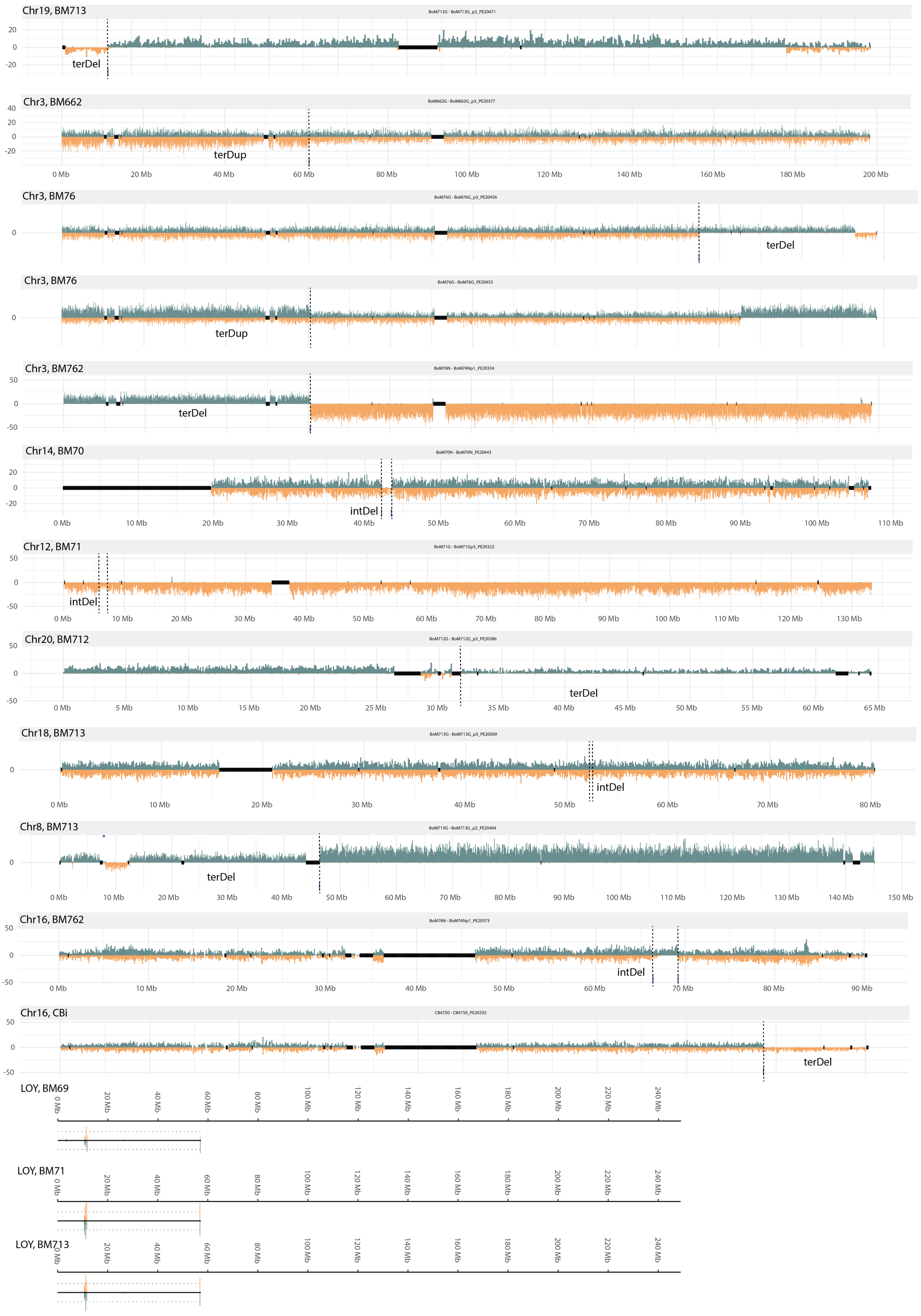
Chr19, BM713
terDel
Chr3, BM662
terDup
0 Mb
20 Mb
40 Mb
60 Mb
80 Mb
100 Mb
120 Mb
140 Mb
160 Mb
180 Mb
200 Mb
Chr3, BM76
terDel
Chr3, BM76
terDup
Chr3, BM762
terDel
Chr14, BM70
intDel
Chr12, BM71
intDel
Chr20, BM712
terDel
Chr18, BM713
intDel
Chr8, BM713
terDel
Chr16, BM762
intDel
Chr16, CBi
terDel
LOY, BM69
LOY, BM71
LOY, BM713
