## Supplementary Materials for "Cell type-specific consequences of mosaic structural variants in hematopoietic stem and progenitor cells"

**Supplementary Figures**

**Supplementary Tables**

**Supplementary Methods**

1. Identification of active X chromosome from the female genome
2. Protein-protein interaction (PPI) network analysis using STRING
3. Similarity analysis for over-represented pathways of dysregulated genes in mSV subclones

**Supplementary Notes**

1. Characteristics of de novo mSVs in HSPCs
2. Comparison with prior surveys of mosaic copy-number alterations and mSVs
3. Investigation of genes associated with local effect of subclonal inversion in BM65
4. Investigation of functional links between dysregulated TFs in the 17p-Del subclone in BM712
5. Potential small deletions at regions of recurrent SCE/mSV formation
6. Analysis of somatic SNVs from the IntoGen Clonal Hematopoiesis Mutation Browser
7. Analysis of somatic SNVs affecting the AR gene in the UK Biobank

**References**

### Supplementary Figures

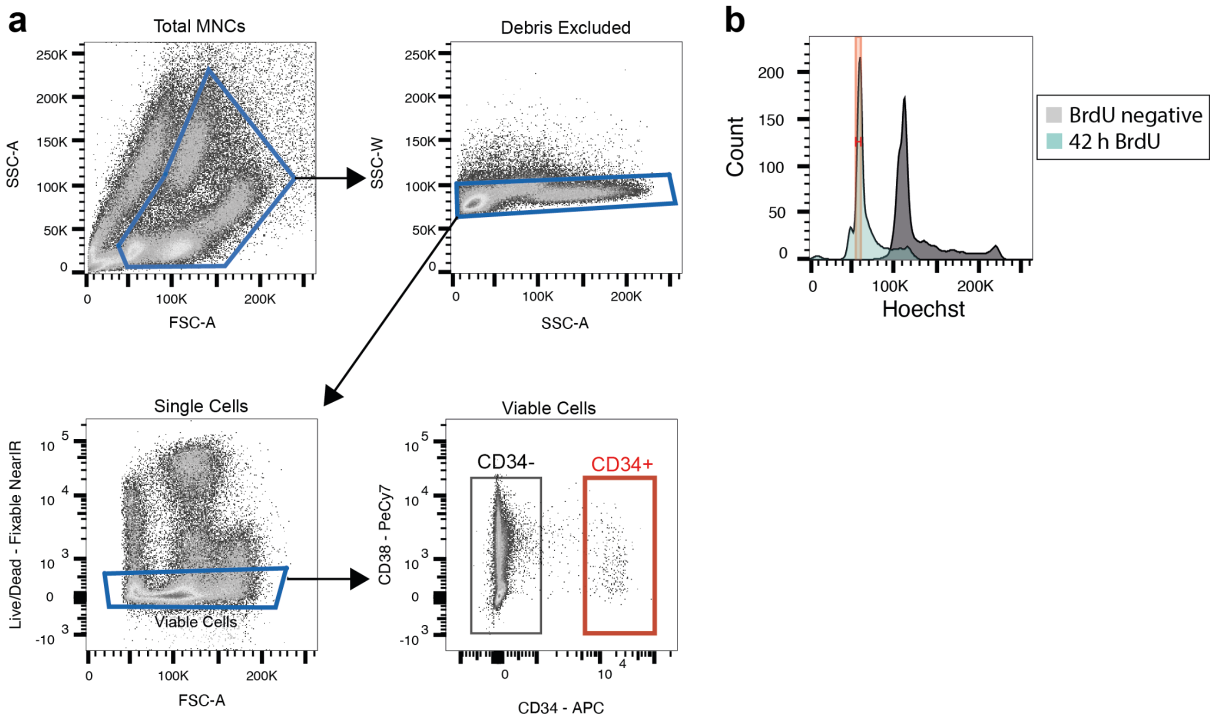


**Figure S1: Sorting strategies used to generate Strand-seq libraries from HSPCs.**
**a**) Gating strategy for isolation of viable CD34+ cells from human bone marrow/umbilical cord blood mononuclear cells. **b**) Gating strategy for sorting of single, BrdU-containing nuclei from cultured HSPCs.

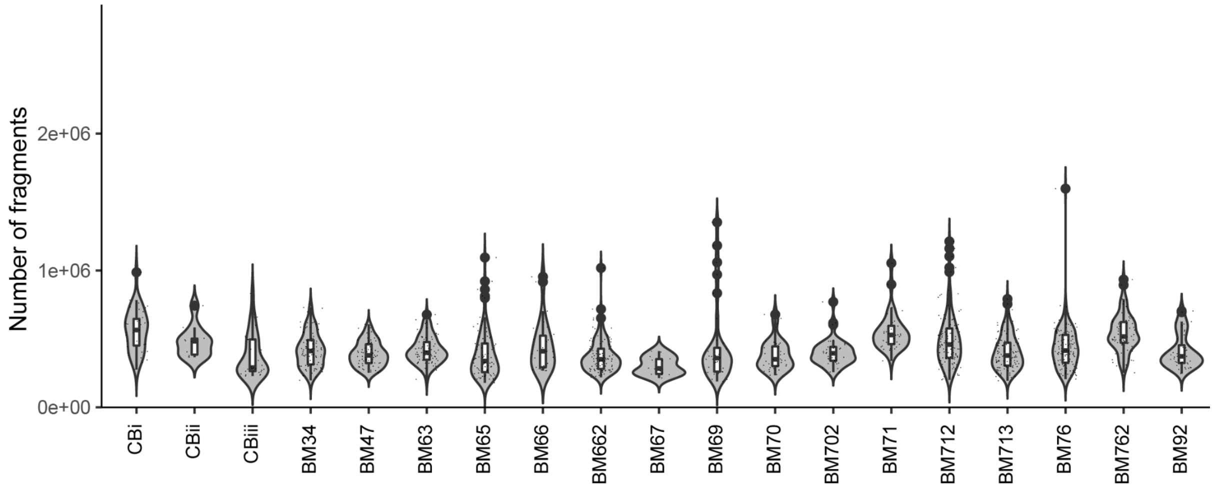


**Figure S2: Number of uniquely mapped fragments per Strand-seq library per donor across cohort.** The violin plots show the number of uniquely mapped fragments per cell profiled from 19 donors. Labels in the x-axis refer to sample names of samples, in which the tissue of origin (CB; cord blood, BM; bone marrow), and the age of each donor, is indicated.

.

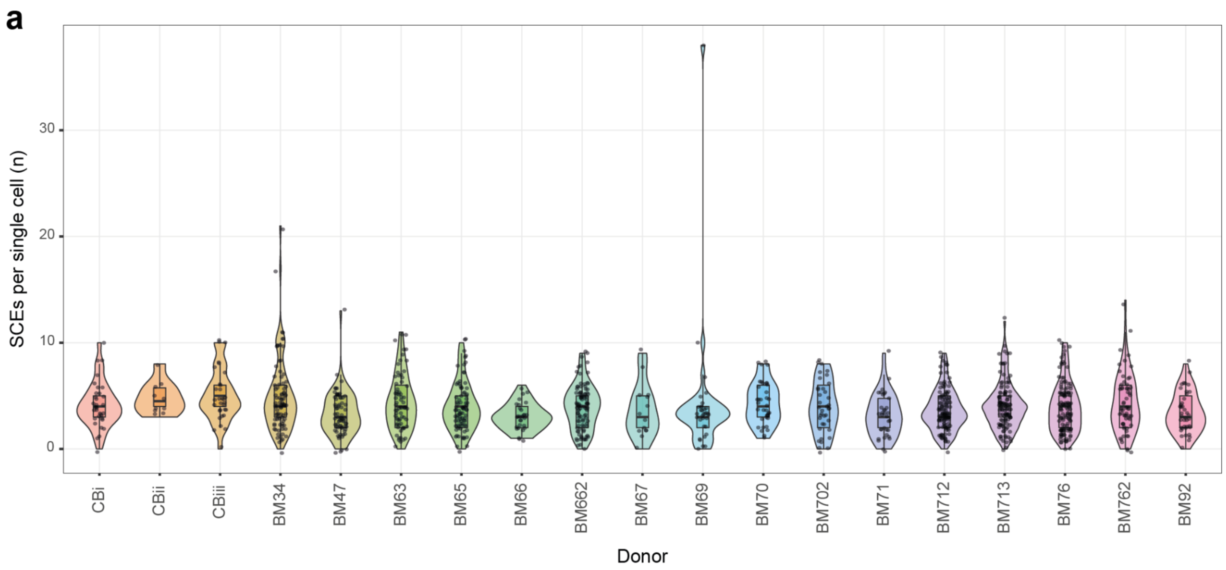


**Figure S3: SCEs per cell do not increase with age in human HSPCs.**
Violin plot showing the number of SCEs per single cell per donor, in order of increasing donor age.

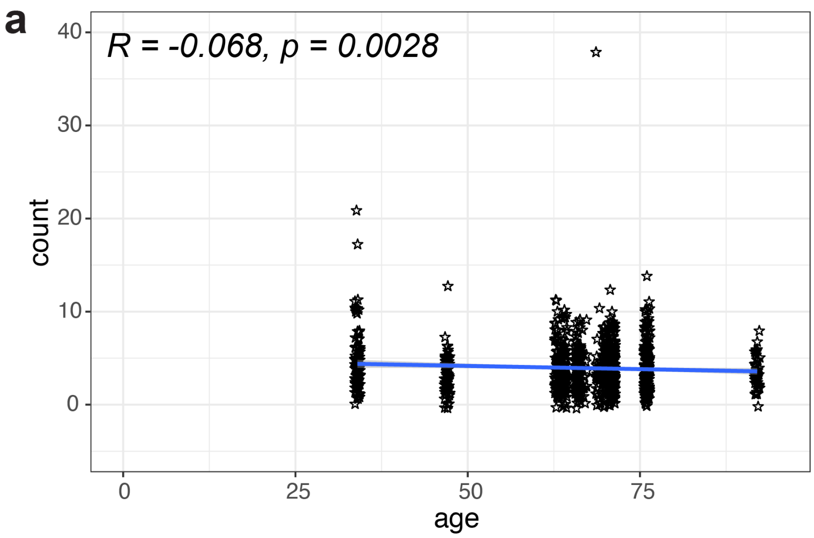


**Figure S4: Anticorrelation of SCE frequency with age is independent of tissue-of-origin.**
Whether UCB samples are included (**Fig. 1f**) or excluded (a), there remains a weak anticorrelation between the age of a donor and the number of SCEs seen per cell (R=-0.068; P=0.0028 excluding UCB).

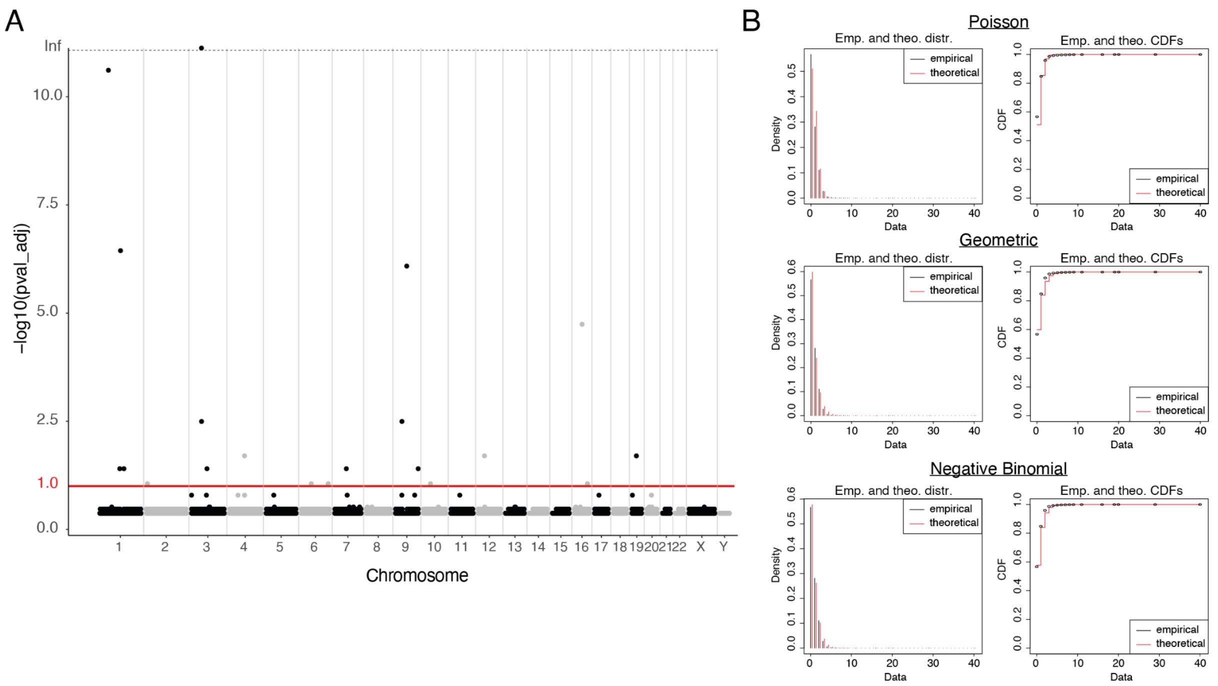


**Figure S5: SCEs occur non-randomly across the genome of human HSPCs.**

**A**) Manhattan plot showing the distribution of SCEs genome-wide (hg38, 500kb bins). Red line denotes the significance cutoff used (10% FDR). The significance of SCE counts per bin were calculated by fitting the permuted data of the number of overlaps per bin genome-wide (quantified using bedtools intersect) to a negative binomial distribution using the fitdist() function from the fitdistrplus package (https://cran.r-project.org/web/packages/fitdistrplus/index.html) (evaluation of distribution of empirical vs theoretical data shown in **B**), and then computing the p-value of the actual data using the fitted negative binomial distribution as a null distribution. The resulting size and mu (size = 1.4013518 (standard error = 0.08457053), mu = 0.6714258 (standard error = 0.01213493)) were used to transform SCE counts per bin into p-values, followed by Benjamini-Hochberg correction[1] to control the FDR.

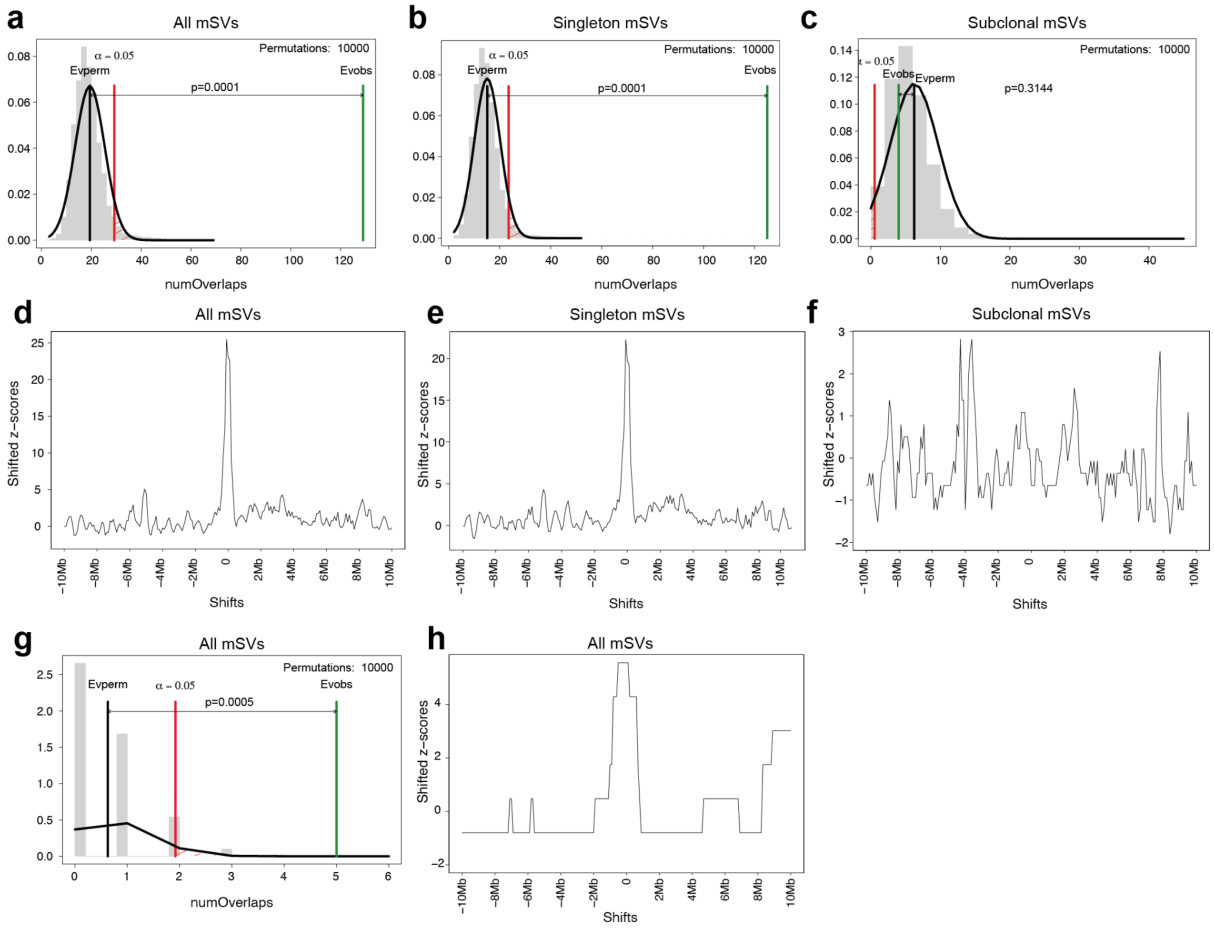


**Figure S6: Permutation summary and local Z-score plots for mSV breakpoints vs SCEs and chromosome fragile sites (CFSs).**

**a**-**c**) Permutation summary plots for 10,000 permutations of breakpoint regions from **a**) all mSVs, **b**) singleton mSVs and **c**) subclonal mSVs vs. all SCEs. **d-f**) Local Z-score plots showing enrichment Z-scores within a 20 Mb window, in 2 Mb bins, for **d**) all mSVs, **e**) singleton mSVs and **f**) subclonal mSVs. **g**) Permutation summary plots for 10,000 permutations of breakpoint regions from all mSVs vs. previously annotated CFSs[2]. **h**) Local Z-score plots showing enrichment Z-scores within a 20 Mb window, in 2 Mb bins, for all mSVs vs previously annotated CFS regions.

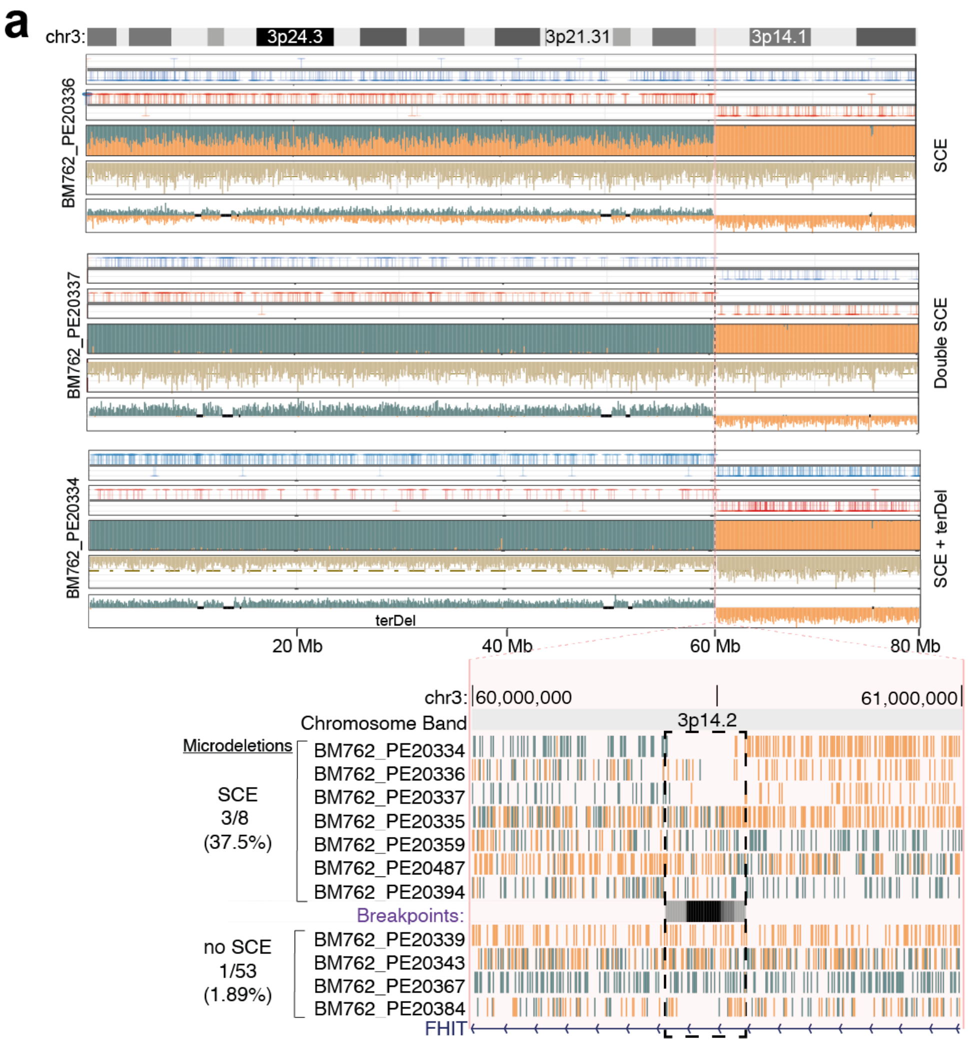


**Fig. S7: *De novo* DNA rearrangement acquisition at SCE hotspots.**

**a**) Upper: Strand-seq data showing recurrent SCE and mSV co-occurrence at the common fragile site (CFS) *FRA3B*, an SCE hotspot. Lower: Zoom in to the breakpoint region at *FRA3B*. Both cells with (top 8) and without (lower 4) SCEs at this CFS show potential deletions, evident at sub-200 kb resolution.

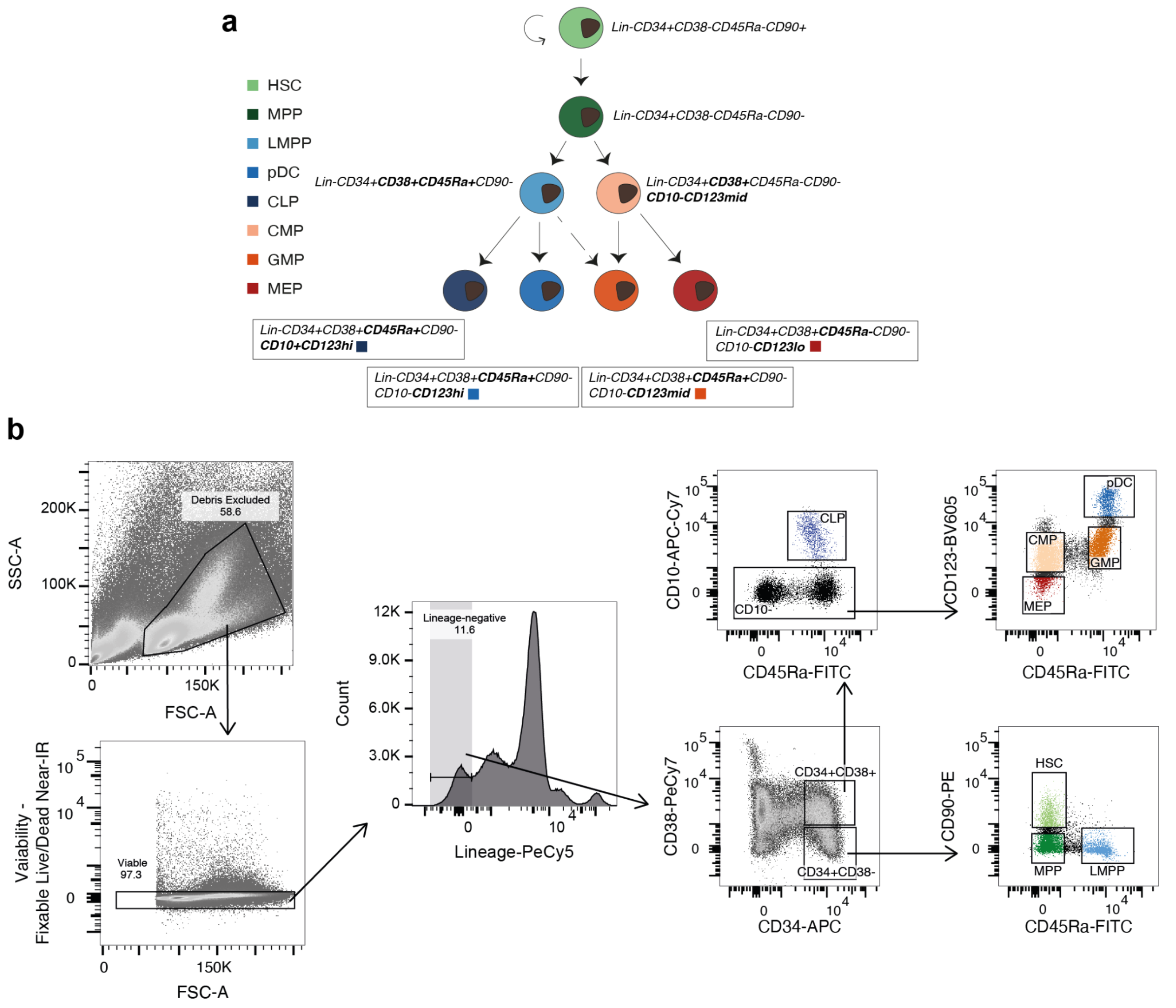


**Figure S8: Isolation of HSPCs for scMNase-seq reference dataset generation.**
**a**) Defining immunophenotypes for 8 HSPC cell types, as previously described in [3]. **b**) FACS gating strategy for isolating each of the cell types mentioned in **a**).

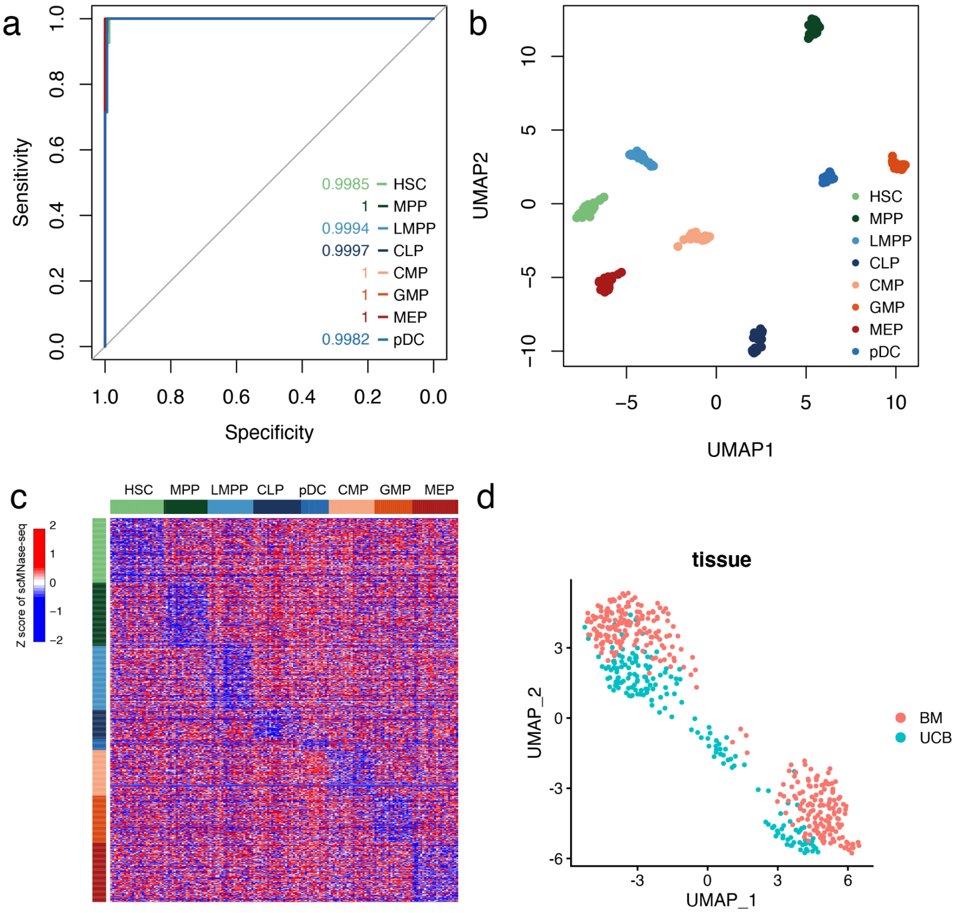


**Figure S9: Umbilical cord blood (UCB)-derived scMNase-seq-based cell-type classifier performance.**

**a**) ROC curve showing leave-one-out cross-validation of the cell-type classifier's performance using single cell NO patterns. **b**) UMAP projection of latent variables from the UCB HSPC cell type classifier. **c**) Heatmap of single cell NO of gene bodies of 175 single bone marrow HSPCs, generated using scMNase-seq. The 899 signature genes depicted (rows) allow for discrimination between 8 bone marrow HSPC cell types (columns). Cells are grouped and colour-coded by immunophenotyped cell-type identity, determined by FACS (**Fig. S8**). Differential NO of marker genes is represented by Z-scores. **d**) UMAP projection of scMNase-seq latent variables, coloured by tissue-of-origin. BM, bone marrow.

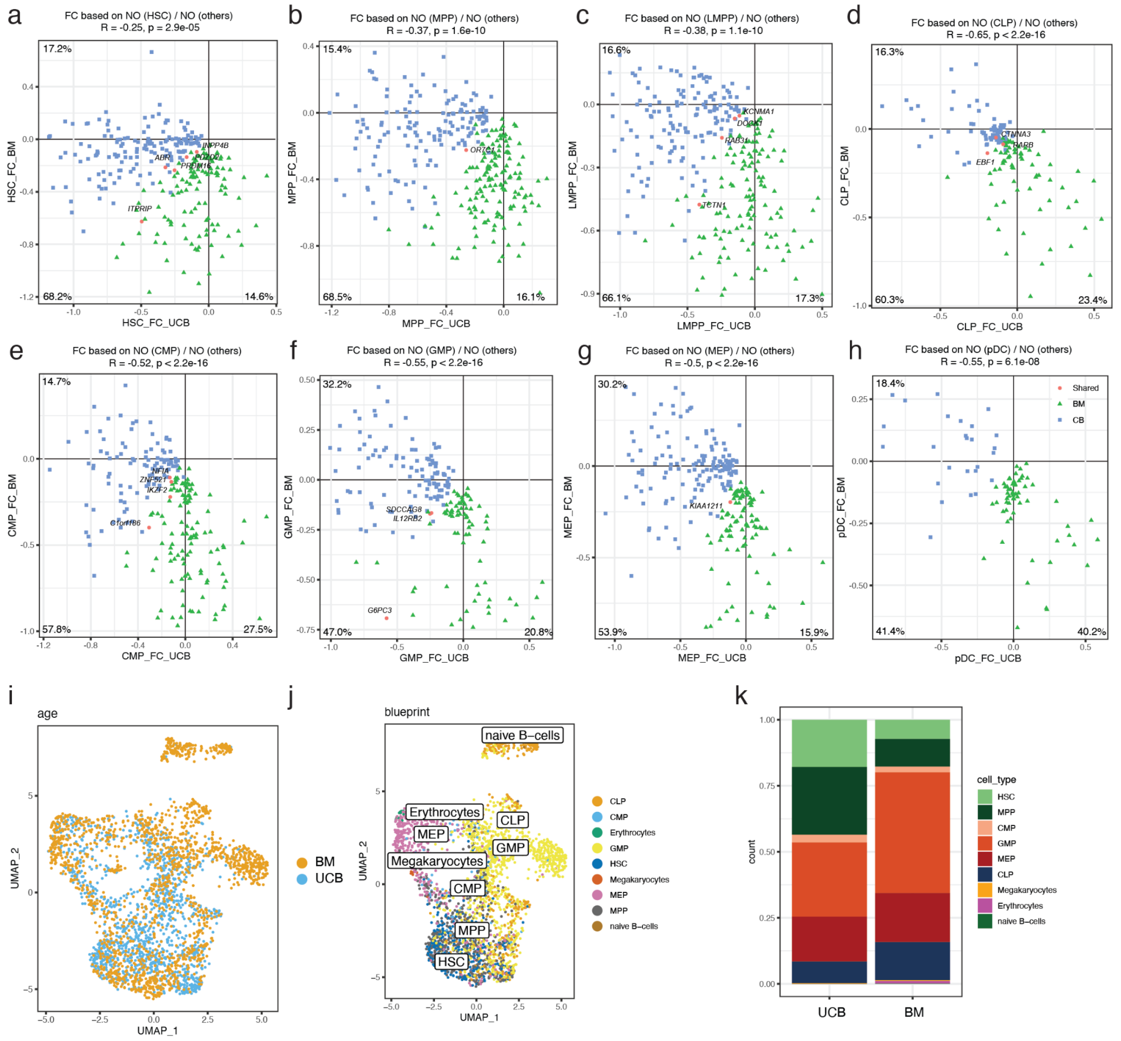


**Figure S10: Comparison of HSPCs from UCB and BM compartments. a-h**) Scatter plots comparing UCB (x-axis) and BM (y-axis) in terms of fold change of nucleosome occupancy (NO) by cell-type (HSC, MPP, LMPP, CLP, pDC, CMP, GMP, and MEP). Each dot represents the cell-type classifier genes selected from UCB (blue), BM (green), or both compartments (pink). The Names of classifier genes shared by both compartments (pink dots) are shown in the scatter plots. *PRDM16* shows significantly decreased NO in HSCs from both BM and UCB, in agreement with its previously determined role in HSC generation and maintenance[4]. However, many genes show distinct NO profiles in BM vs UCB, with only 21 genes appearing in both BM- and UCB-derived NO classifiers. **i**) UMAP of single-cell transcriptome data of HSPCs from UCB and adult bone marrow, obtained from [5]. **j**) Cell-types in the scRNA-seq were annotated using the singleR package[6] based on the blueprint reference[7]. **k**) Cell-type composition of scRNA-seq from UCB and BM shows that MPPs are highly enriched in UCB compared to BM HSPCs.

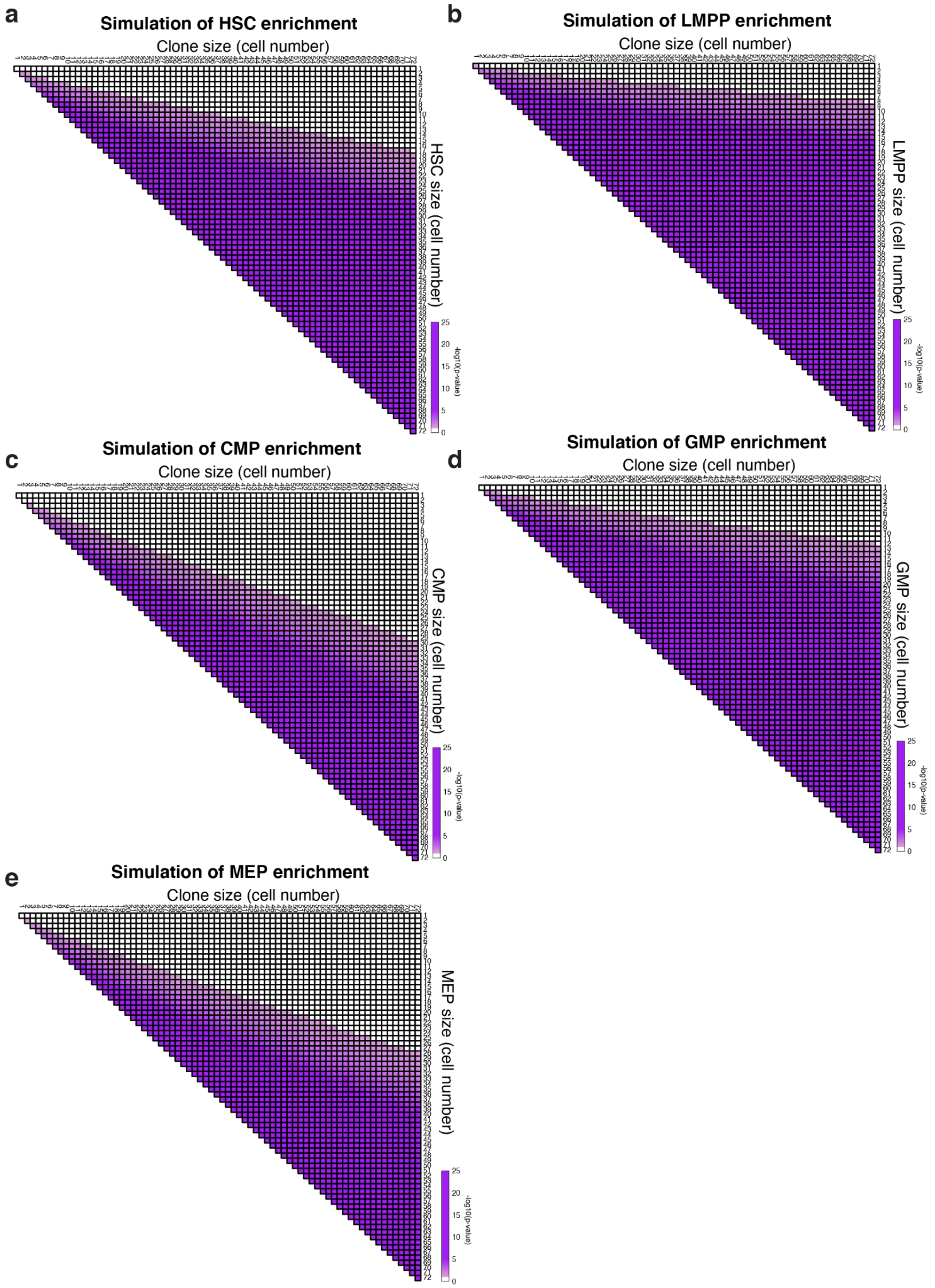


**Figure S11: Simulation analysis of cell-type bias for different subclone sizes.** Each pyramid plots show the number of cells in each cell-type (y-axis) required to statistically show the cell-type bias for given size of mSV clones (x-axis). This simulation analysis was performed for the mSV clone size range between 1 to 72 for five different cell-types we used for the cell-type enrichment analysis in **Fig. 3b**.

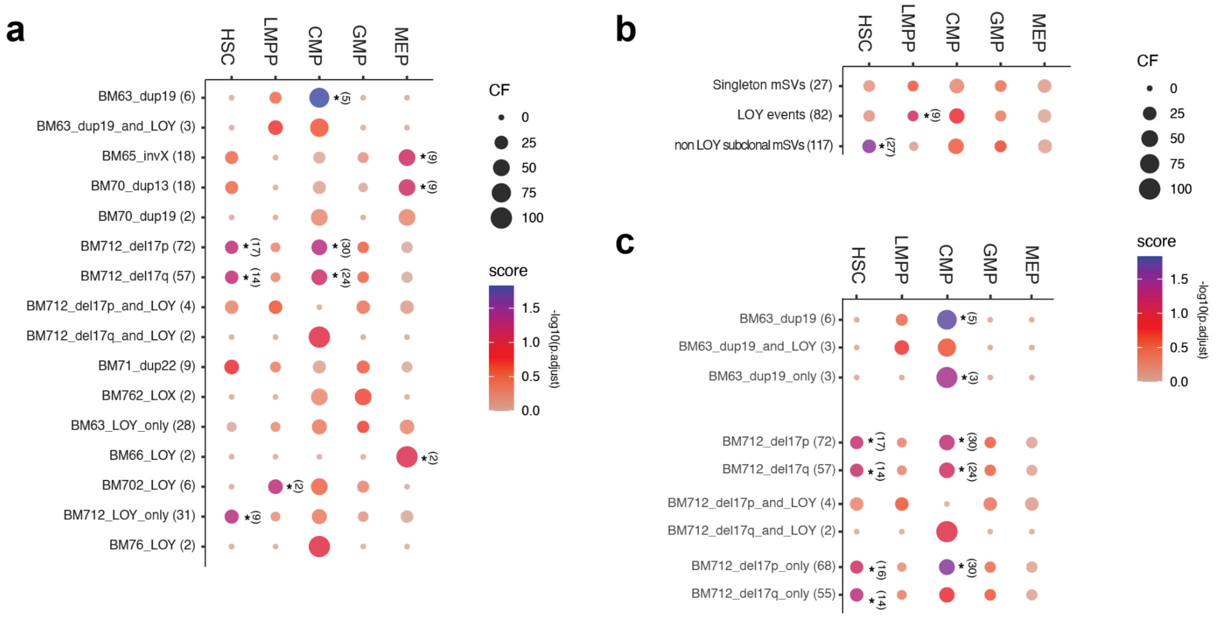


**Figure S12: Cell type enrichment across all unique genotypes at the a) single donor/single genotype and b) cross-donor/cross-genotype levels. a)** Extended dotplot of results of the cell-type enrichment analysis for each mSVs identified, showing the CF, enrichment and significance of enrichment in cell type per mSV sub-clone vs. an idealised control. This analysis was extended from the result in **Fig. 3b** to investigate the effect of whole chromosome losses (LOY, LOX), and the effect of newly arisen mosaicisms within subclones harboring mSVs. For instance, BM63_dup19_and_LOY denotes the LOY subclone originated from the bigger subclone harboring a duplication on chromosome 19. **b)** Dotplot of combined enrichments for the cells has singletons ('Singleton mSVs'), has LOYs ('LOY events'), or harboring subclonal mSVs that are not LOYs ('non LOY subclonal mSV'). **c)** Comparison of cell-type enrichment between subclones acquired secondary mSVs, and the bigger subclones they originated from.

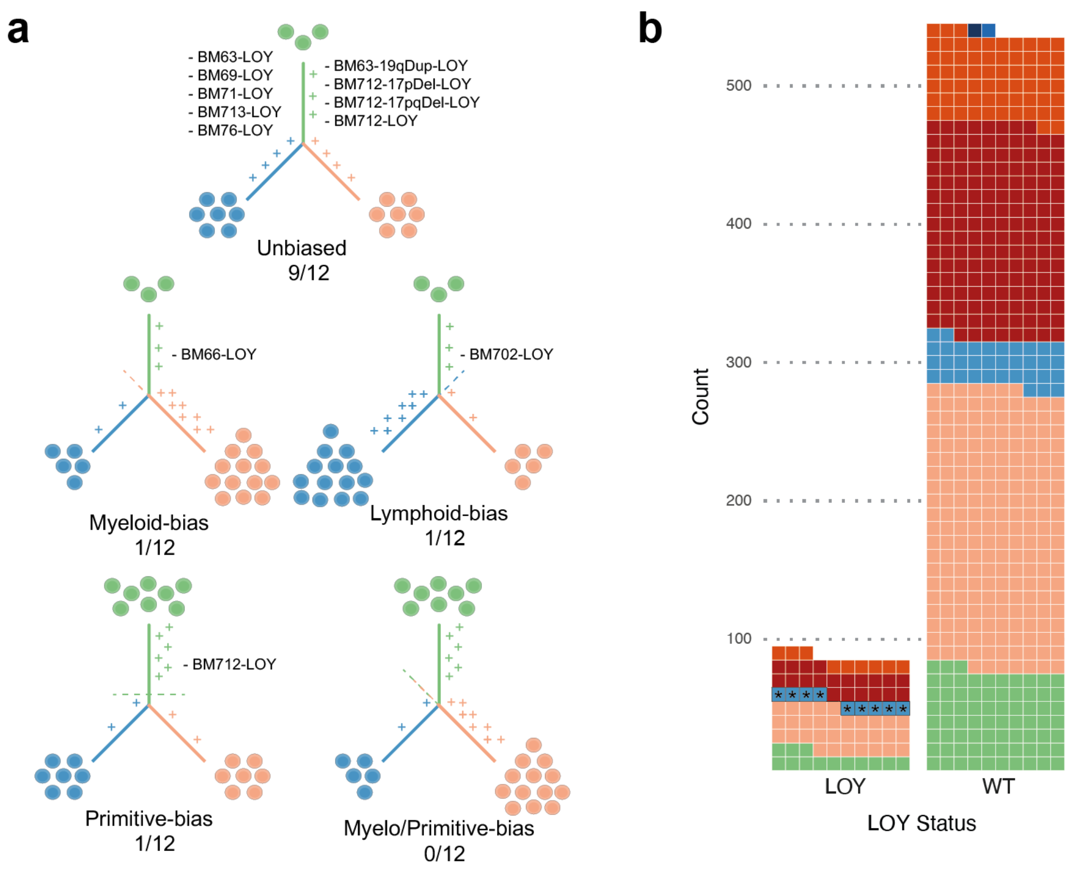


**Figure S13: Lineage biases observed for LOY.**

**a)** Lineage-level biases observed per donor for LOY subclones. **b**) Cohort-level bias observed for LOY. When compared to an idealised control population of HSPCs from male donors, we observe a significant enrichment of LOY in the LMPPs.

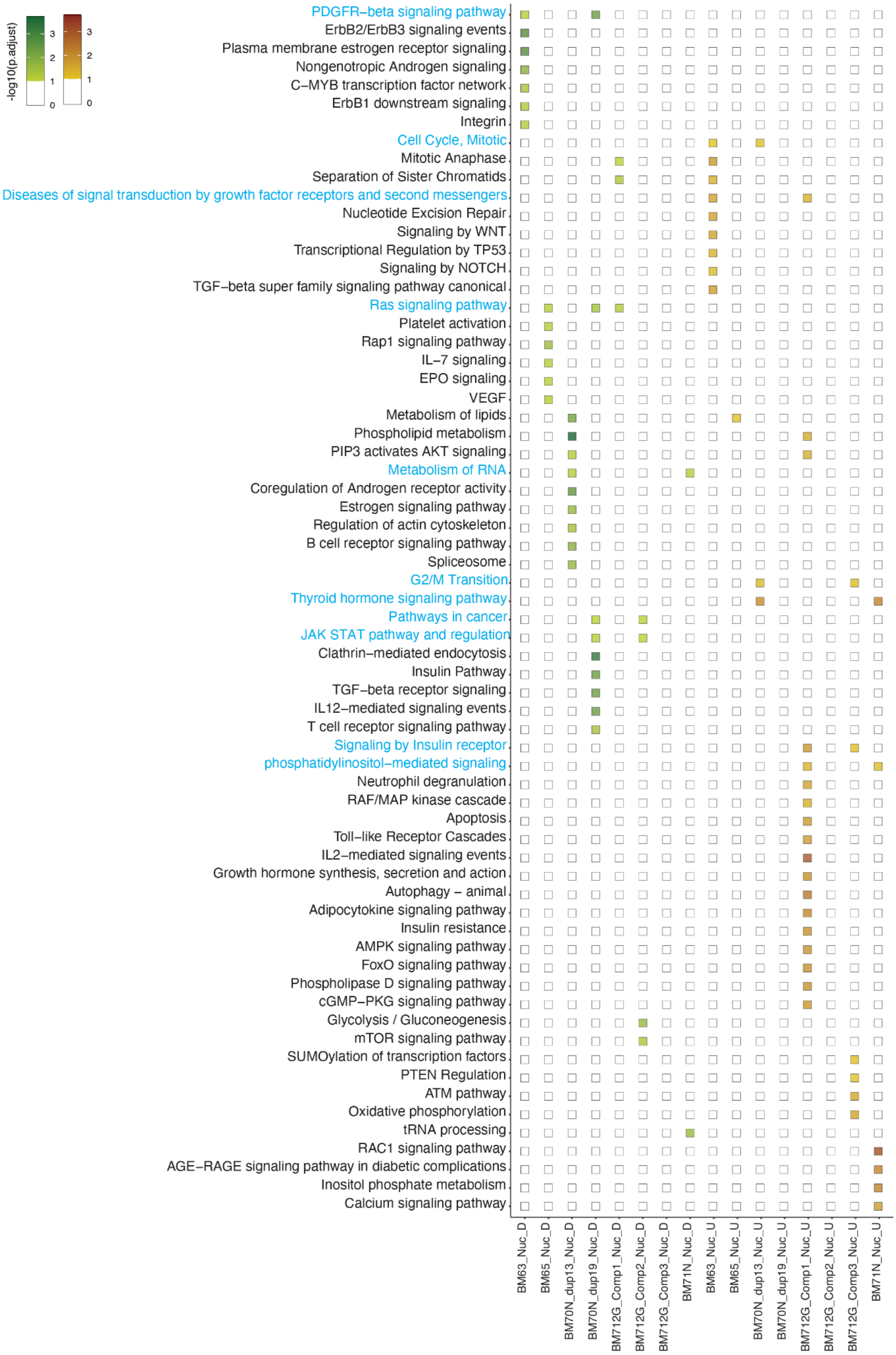


**Figure S14: Over-represented pathways of dysregulated genes in mSV subclones.** Plot of pathway enrichment analysis for subclonal mSVs across the cohort. Pathways showing enrichment in at least one mSV are shown here. Pathways enriched in 2 or more mSVs in the same direction are highlighted in blue font (those pathways are also depicted in **Fig. 1e**).

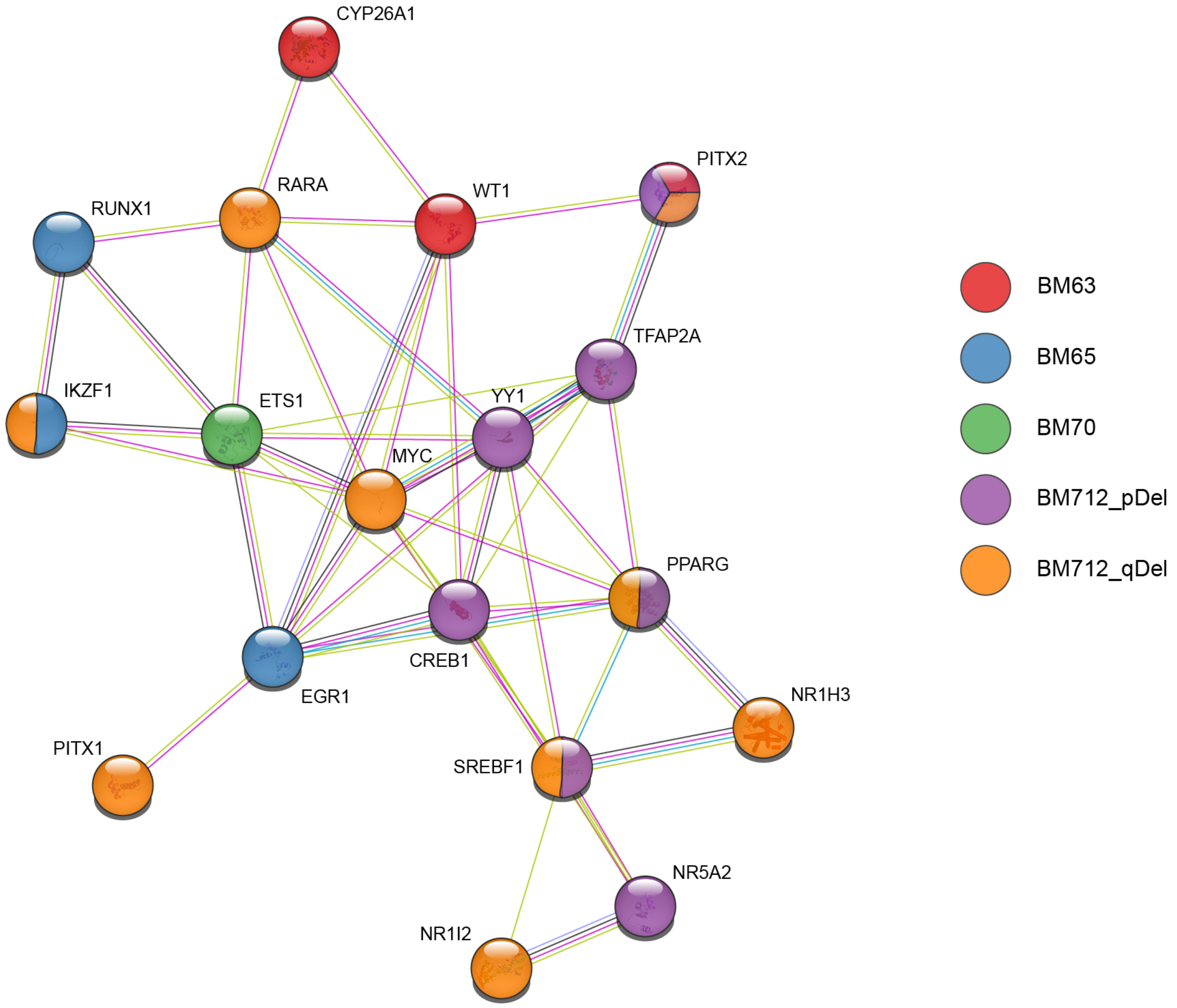


**Fig. S15: Protein-protein interaction network identified from differentially active TFs across all mSV subclones.**

STRING network of all differentially active TFs identified in samples exhibiting subclonal mSVs. TFs are coloured by sample and mSV. For a significantly enriched gene-sets, see **Table S20**.

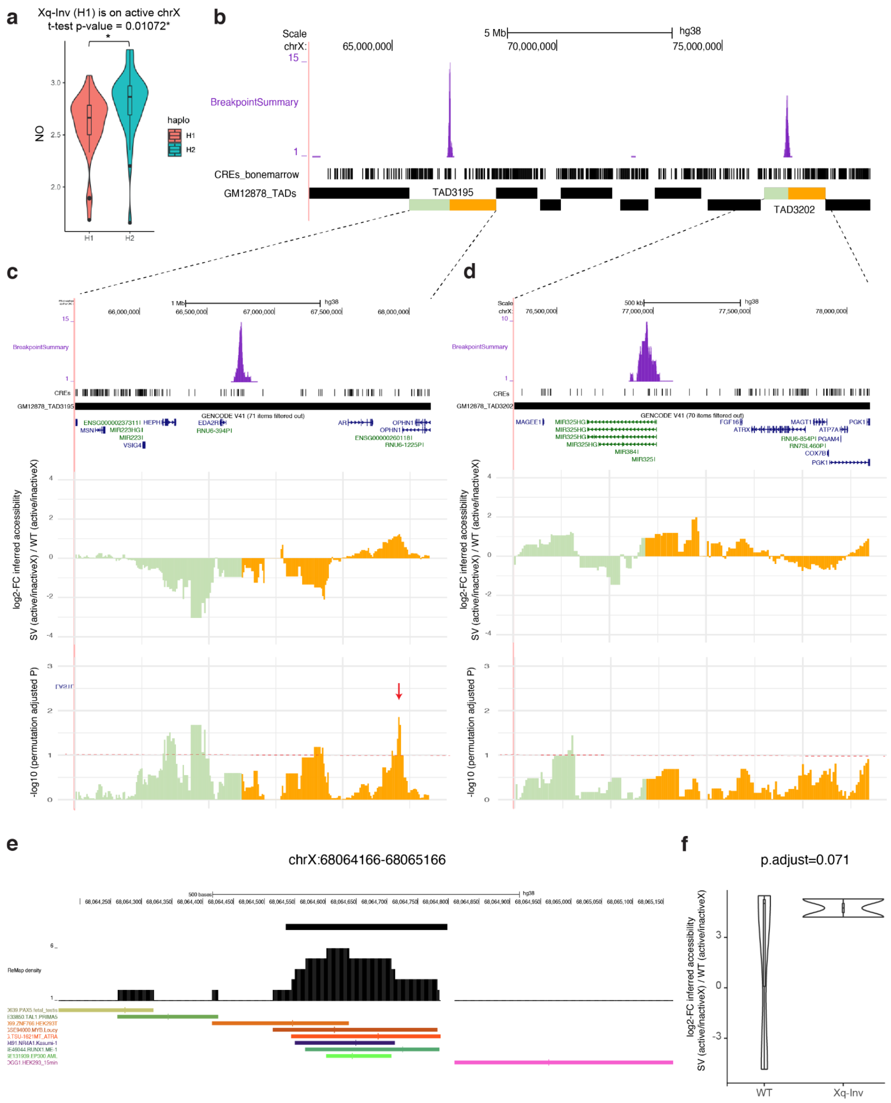


**Figure S16: Investigation of *cis* effects of the mosaic inversion in BM65.**
(**a**) The inversion detected in donor BM65 is mapped to homolog 1 (H1) of chromosome X, using the Strand-seq data generated in this sample. We first checked whether the inversion-affected homolog represents the epigenetically-active or -inactive X chromosome, utilising the haplotype-resolved NO profiles. The violin plot shows the average NO for H1 and haplotype 2 (H2) in each single-cell. Using a t-test to compare the two homologs indicates that the inversion harboring homolog (H1) represents the active X chromosome ($P$<0.01). (**b**) Browser track snapshot showing that two TADs (TAD3195, TAD3202) are disrupted as a result of the inversion. (**c-d**) Genome browser tracks showing log2-fold changes in haplotype-resolved NO (indicative of changes in chromatin accessibility[8]) between the mSV clone and WT cells on the active X chromosome, with permutation adjusted $P$-values (shown for TAD3195 (**c**) and TAD3202 (**d**), respectively). Red arrow indicates the sliding window showing the most significant difference between mSV clone and WT cells. (**e**) *Cis*-regulatory element located in the most significantly different sliding window highlighted in panel (**c**). NO of this individual CRE was significantly lower in the mSV subclone compared to WT cells (p.adjust < 0.071), indicating increased chromatin accessibility. TF motif sequences located in this region are depicted below. (**f**) Violin plot showing the single-cell level distribution of log2-fold changes in inferred chromatin accessibility[8] between the mSV subclone and WT cells.

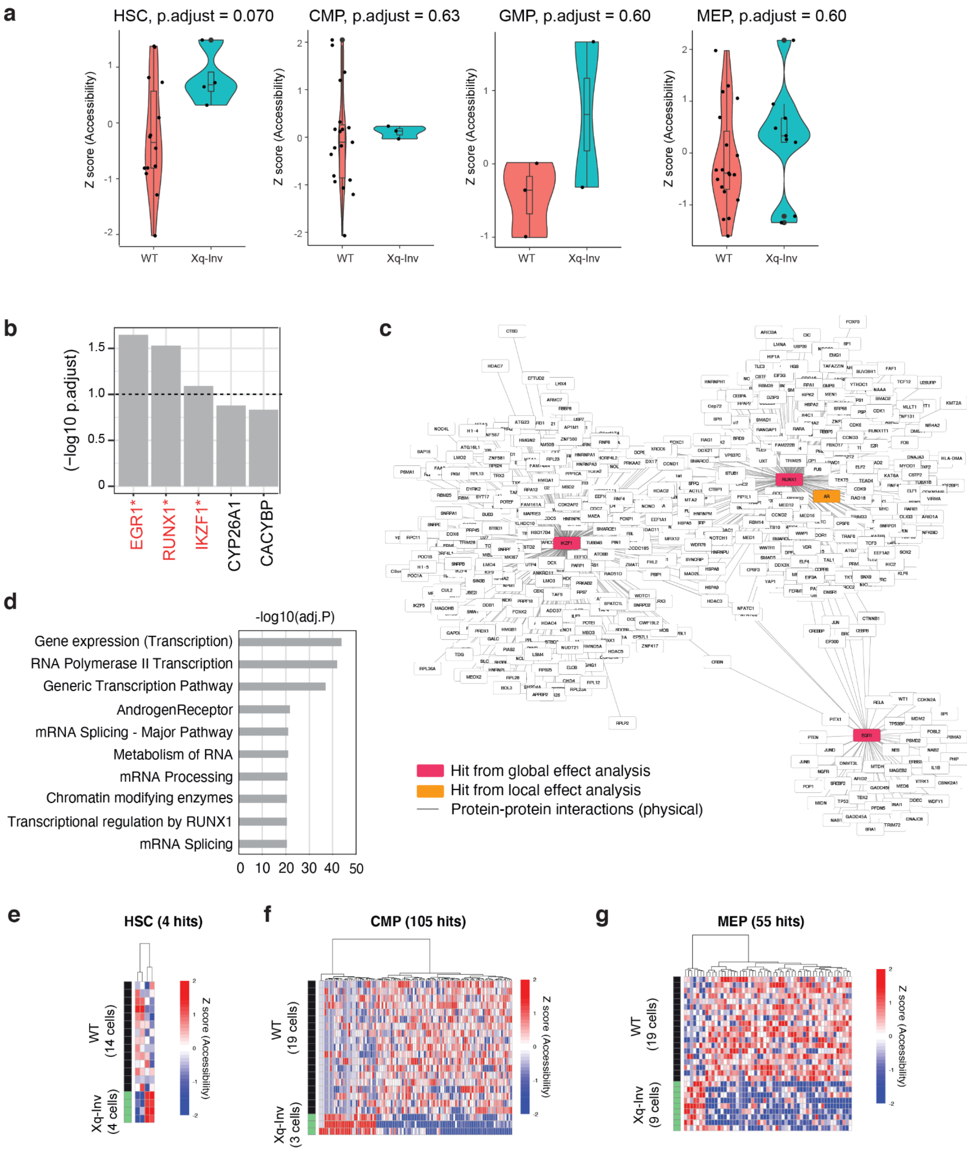


**Figure S17: Identification of global effects of Xq-Inv in BM65 using TF activity analysis (a)** Z-scores of the inferred activity of AR target genes, based on scNOVA, were compared for each cell-type separately. (**b**) TFs identified from the TF-target over-representation analysis[8–10] of deregulated genes in the inversion subclone. This analysis reveals 3 TFs with differential activity in Xq-Inv cells: EGR1, RUNX1, and IKZF1 – all of which are linked to AR signaling[11]. (**c**) Protein-protein interaction network representing the first interactors of all three significant TFs identified in (**b**), using an FDR of 10%. Protein-protein interactions were obtained from the NCBI gene resource (https://www.ncbi.nlm.nih.gov/gene/). The 3 TFs identified in (**b**) are highlighted in red. The AR protein, which in this network is a direct interactor of *RUNX1,* is highlighted in orange. (**d**) Pathways over-represented by the genes involved in the network in (**c**) based on the ConsensusPathDB[12]. (**e-g**) Dysregulated genes in the inversion subclone identified by scNOVA in a cell-type specific manner, for HSCs (**e**), CMPs (**f**), and MEPs (**g**). We find that 3/4 differential NO genes of HSCs are AR targets[13], while 23/105 and 12/55 differential NO genes of CMPs and MEPs are AR targets[13], respectively.

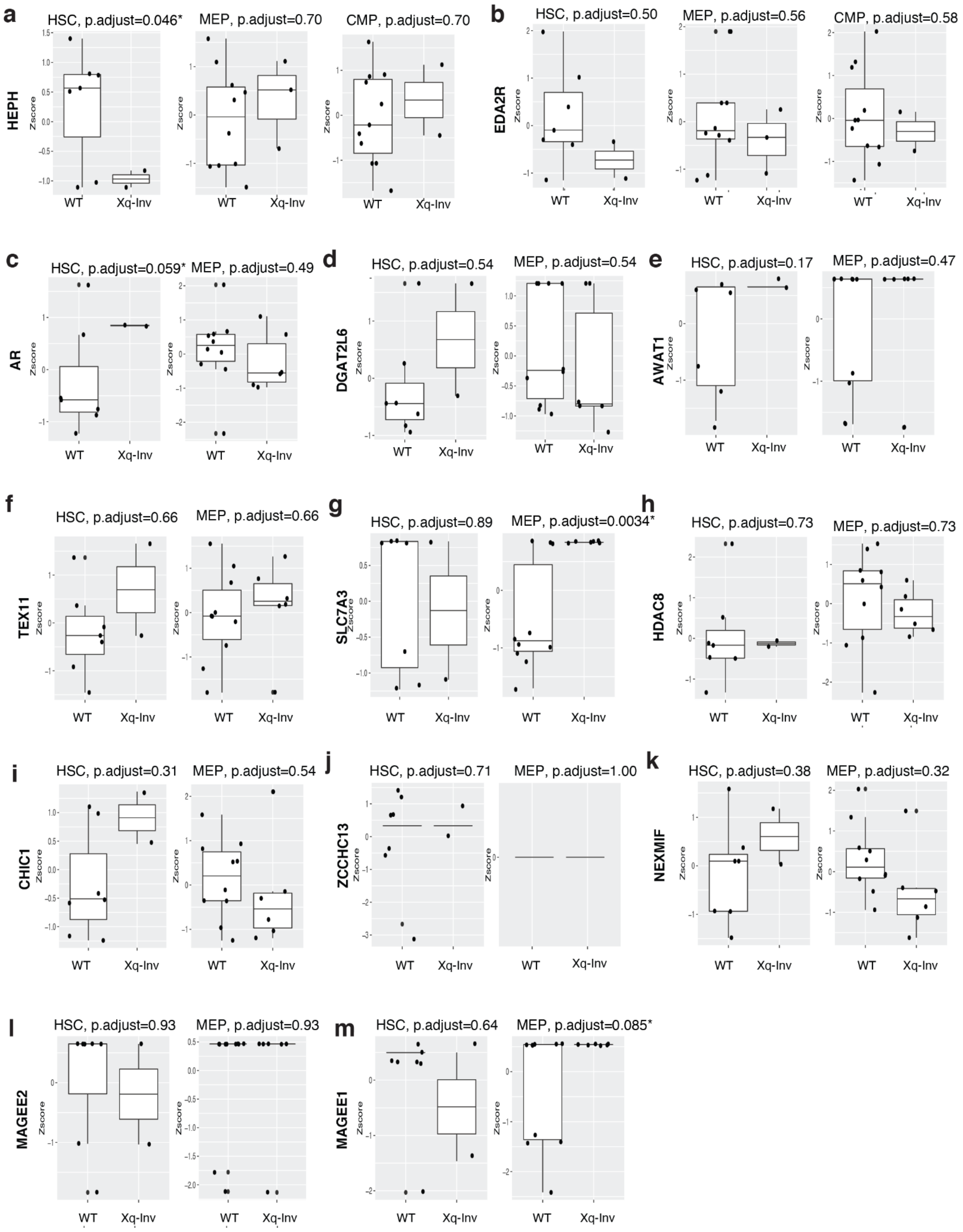


**Figure S18: Inferred haplotype-specific gene activity of genes located within the inverted region and affected TAD boundaries.** In total 13 genes are found in the TADs affected by inversion breakpoints or within the inverted segments (**Fig. 4a**). For those 13 genes, we compared the NO at gene bodies for the mSV clone and the WT cells in the active X homolog in a cell-type specific manner (**a-m**). Z-scores in the Y-axis indicate gene body-based inferred gene activities, inferred by scNOVA. This analysis requires at least two cells with WC configuration to resolve the haplotype into the active and inactive X homolog. Significance testing was achieved using t-tests. For the genes that reside in the TADs affected by the inversion breakpoints, but outside of the inverted region, we performed testing in HSCs, CMPs, and MEPs. For genes residing within the inverted segment, we performed testing in HSCs and MEPs (as the segregation pattern for the inverted segment and the rest of the chromosome is different, explaining differences in our capacity to test for different cell-types). Cell-type and adjusted p-values of genes showing significant difference of NO are indicated with asterisks (FDR 10%).

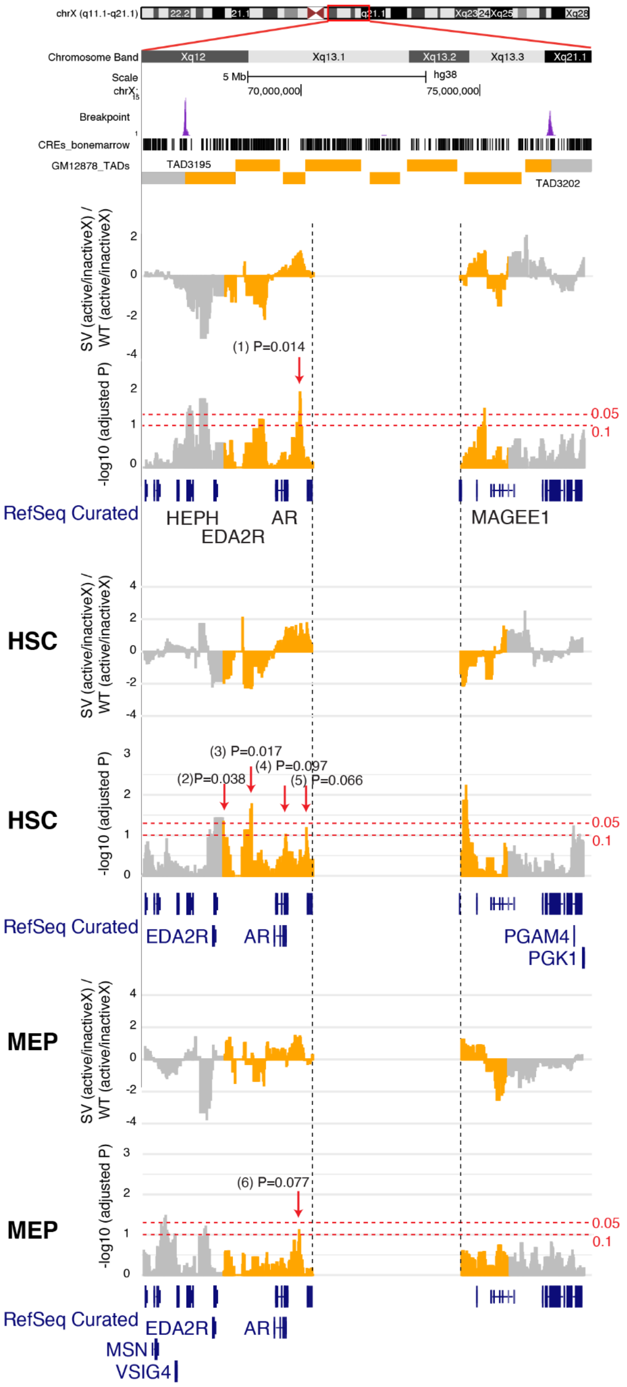


**Figure S19: Investigation of cell-type resolved local effects of the mosaic inversion on chromatin accessibility.** For the mSV affected TAD boundaries (TAD3195, TAD3202), we compared the NO profiles of the active X homolog between the mSV clone and the WT cells in a cell-type resolved fashion. We performed this analysis for HSCs and MEPs, as the cell count of those cell-types fulfilled the minimum required cell count needed for statistical testing. For each cell type, log2 fold changes of the mSV (active/inactiveX) subclone compared to WT (active/inactiveX) cells, as well as permutation adjusted p-values are shown in the browser tracks. Additionally, names of the nearest genes adjacent to significant peaks (FDR 10%) are highlighted in the RefSeq genes track. Genomic coordinates of inferred differentially accessible peak regions are as follows: (1) chrX:67730000-68090000 (2) chrX:66400000-66910000 (3) chrX:66940000-67350000 (4) chrX:67580000-67880000 (5) chrX:67900000-68200000 (6) chrX:65640000-66150000.

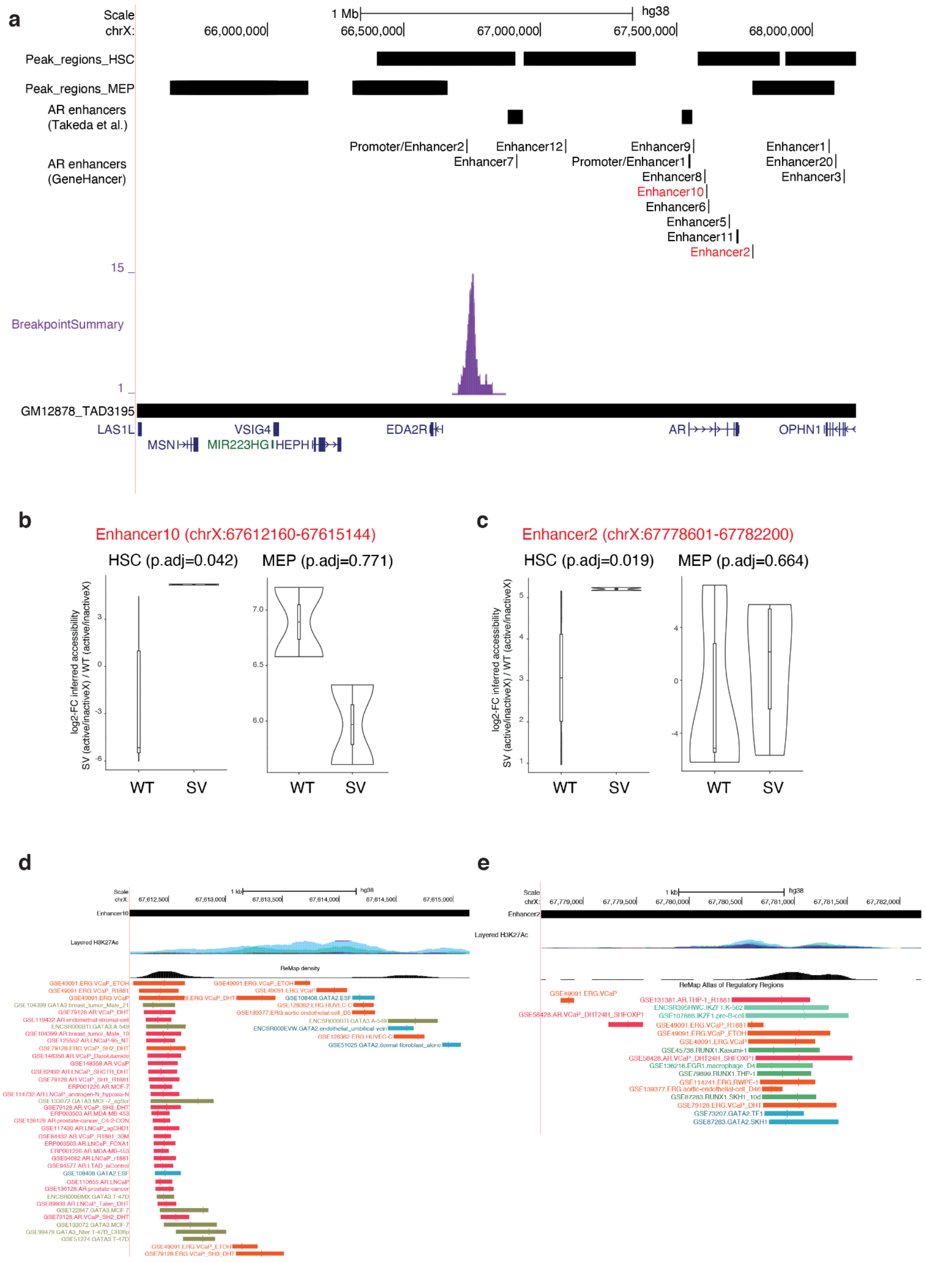


**Figure S20: Cell-type specific inferred accessibility of individual AR enhancers. (a)** A browser track shows the genomic locations of peak regions with differential accessibility for HSCs and MEPs based on analyzing Strand-seq data, and the annotated AR enhancers based on GeneHancer[14] and prior literature[15]. Individual enhancers showing significant differential accessibility inferred by NO are highlighted in red. (**b-c**) Violin plots show the inferred haplotype-resolved accessibility for WT cells (left) and the mSV subclone (right) for enhancer 10 (**b**) and enhancer 2 (**c**). Significance was established using the Wilcoxon rank-sum test. (**d-e**) Browser track snapshots depicting TF binding sites within enhancer 10 (**d**) and enhancer 2 (**e**) according to the ReMap Atlas[16].

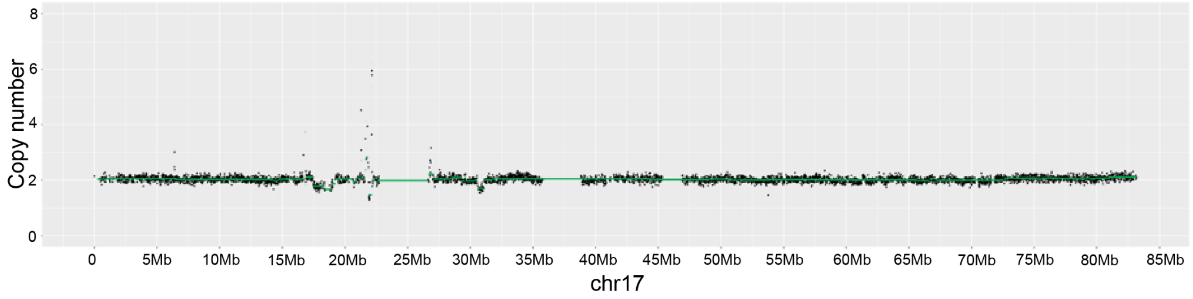


**Figure S21: Verification of mosaic deletions in BM712 using whole genome sequencing (WGS)** Chromosome 17 read coverage plot of whole genome sequencing (WGS) data obtained from BM712, revealing the presence of the mosaic 17p-Del and 17q-Del events in sorted CD34-cells. Split read and paired-end analysis using the Delly2 tool[17] identify the breakpoints of the 17q-Del event with base pair resolution (chr17:30602839-31097552), and validate the exon 1 deletion of *NF1*.

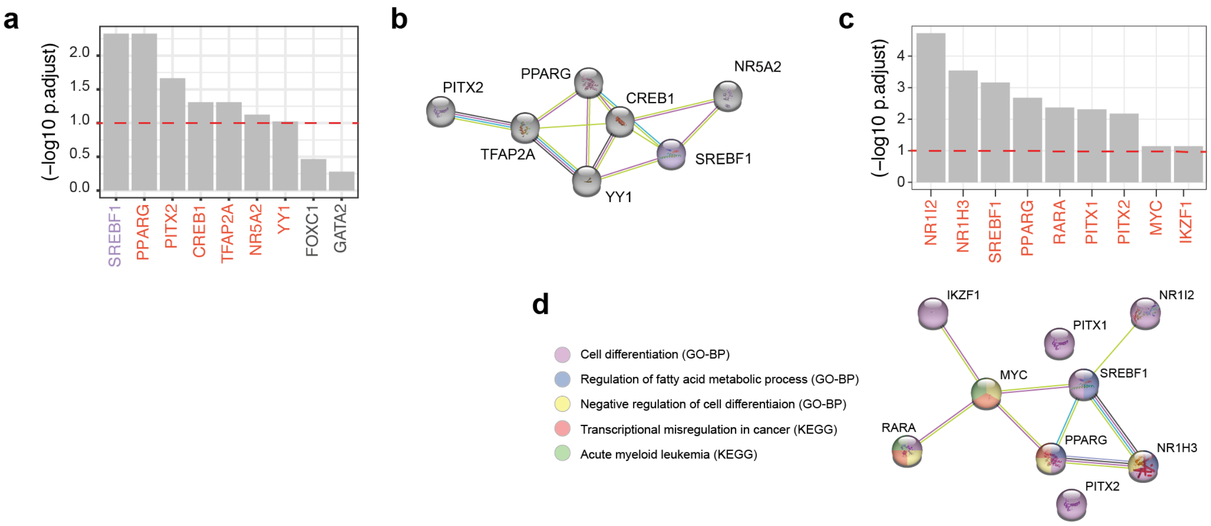


**Figure S22: TF activity and interaction analysis for the 17p- and 17q-Del subclones.**
**a,c**) TFs whose targets show significantly differential NO in 17p-Del cells (**a**) and 17q-Del cells (**c**) vs. WT cells. Significant TFs are coloured (10% FDR threshold); and genes within the specific deleted locus are coloured in purple. **b**) STRING network of TFs in (**a**) (PPI enrichment $P$=3.57e-08, hypergeometric test[18]). **d**) STRING network of TFs in (**c**) (PPI enrichment $P$=0.00838, hypergeometric test[18]). Proteins are coloured based on their contribution to relevant significant genesets listed in the legend.

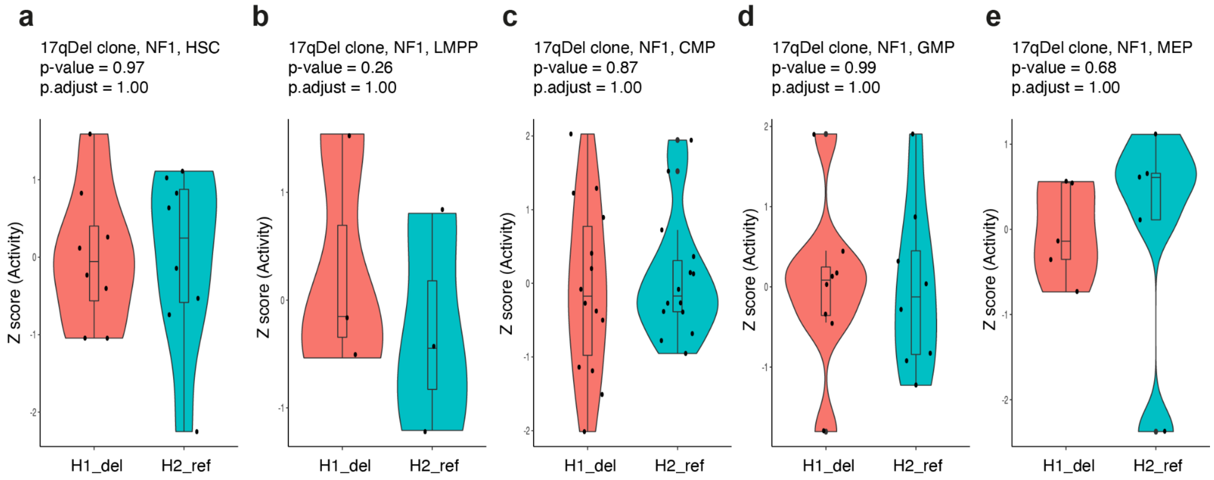


**Figure S23: Haplotype-resolved *NF1* activity based on NO in specific cell types**

Using scNOVA, we extracted the haplotype-resolved NO at the *NF1* gene body for the 17q Del clone and calculated the Z-score of the inferred gene activity for H1 (the rearranged homolog) and H2 (the WT homolog). The difference between two homologs was evaluated using t-test followed by Benjamini-Hochberg multiple testing correction. We performed this analysis in a cell-type specific manner for **(a)** HSC, **(b)** LMPP, **(c)** CMP, **(d)** GMP, and **(e)** MEP. There is no significant difference in NO between the two haplotypes at the *NF1* gene body, suggesting that the *NF1* gene fragment, which is truncated at its 5`-end (exon 1 deletion), is expressed from the rearranged homolog.

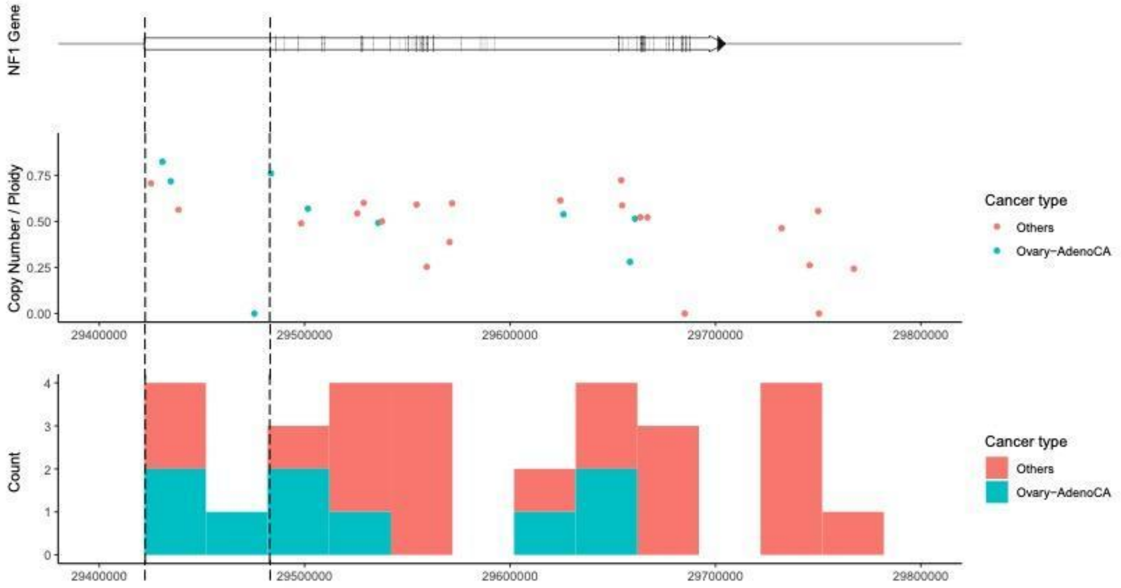


**Figure S24: Somatic deletion breakpoints in intron 1 of the *NF1* gene in PCAWG samples**

Somatic deletion data were downloaded from the PCAWG resource of 2,583 whole cancer genomes[19]. Only deletions fully spanning the first exon of *NF1* are shown. The top panel shows the *NF1* gene structure; the middle panel shows the estimated copy number normalized by ploidy with each point representing a PCAWG donor; the bottom panel shows the distribution of breakpoints. The X-axis depicts the breakpoint coordinates in chromosome 17 (hg19). Notably, 3 out of 5 samples exhibiting an intron 1 deletion breakpoint are ovarian cancer samples – with such deletions seen in 3% of ovarian cancers (3 out of 110 ovarian cancer cases in the whitelisted PCAWG resource).

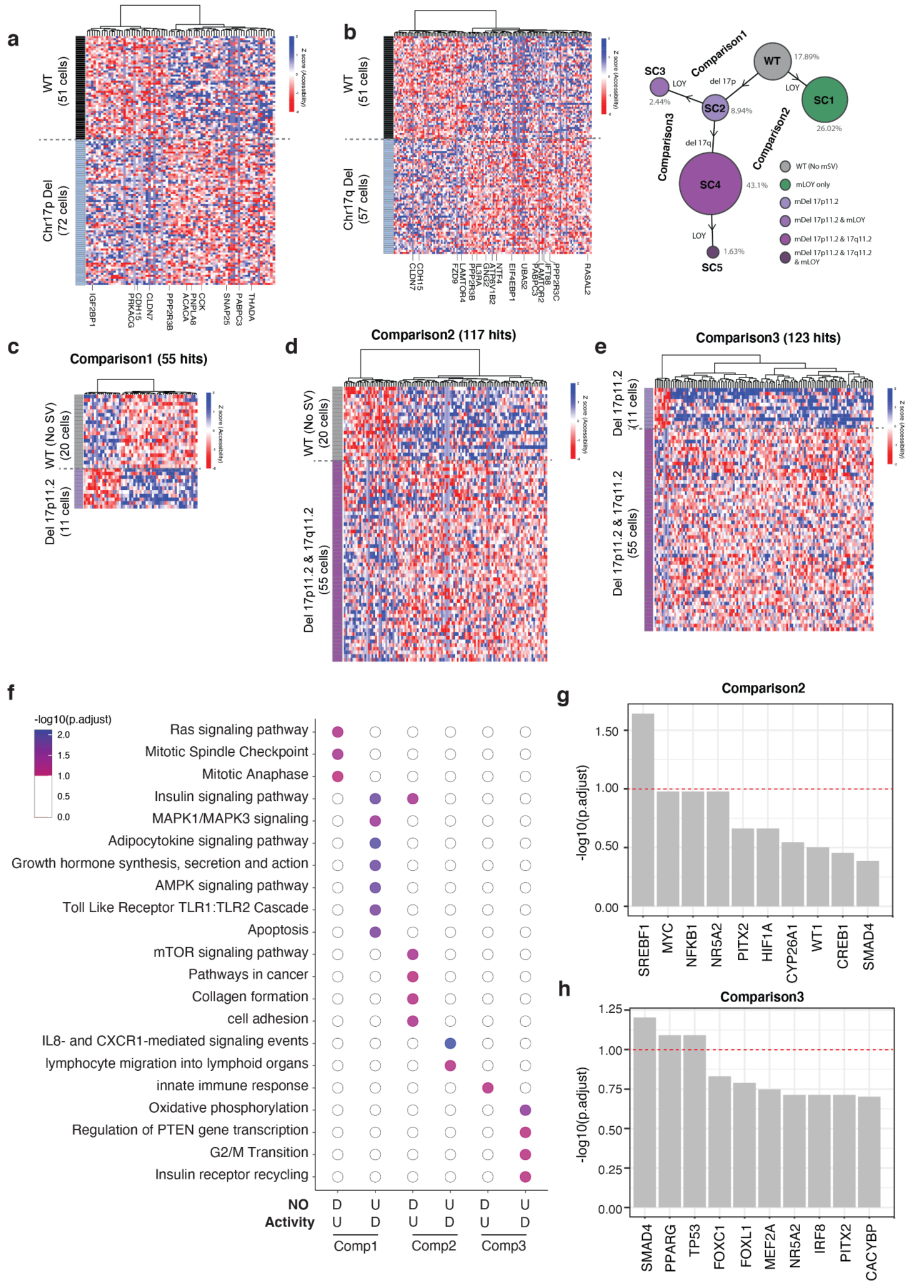


**Figure S25: Inference of altered gene activity in mSV subclones detected in BM712**

(**A**) Differentially active genes between cells harboring 17p mDel and the cells without 17p-Del (WT (no 17p-Del)), using scNOVA[8]. (**B**) Differentially active genes between cells harboring 17q-Del and WT cells (no 17p-Del). (**C-E**) Pairwise comparison of differential gene activity between cells bearing 17p11.2-Del (11 cells), cells bearing 17p11.2-Del and 17q11.2-Del (55 cells), and WT cells (no mSV, 20 cells). (**F**) Pathway over-representation analysis using ConsensusPathDB[12] for the genes identified in the pairwise comparisons in (**C-E**). Significant pathways were identified by controlling the FDR at 10%. In the x-axis, U and D indicate inferred Up and Down-regulation, respectively. Comp1: 17p11.2-Del (11 cells) vs. WT (no mSV, 20 cells); Comp2: Del 17p11.2-Del & 17q11.2-Del (55 cells) vs. WT (no mSV, 20 cells); Comp3: 17p11.2-Del & 17q11.2-Del (55 cells) vs. 17p11.2-Del (11 cells). (**G-H**) TF-target over-representation analysis for the dysregulated genes identified in the Comp2 (**G**) and Comp3 (**H**). For Comp1, no significant TF was identified.

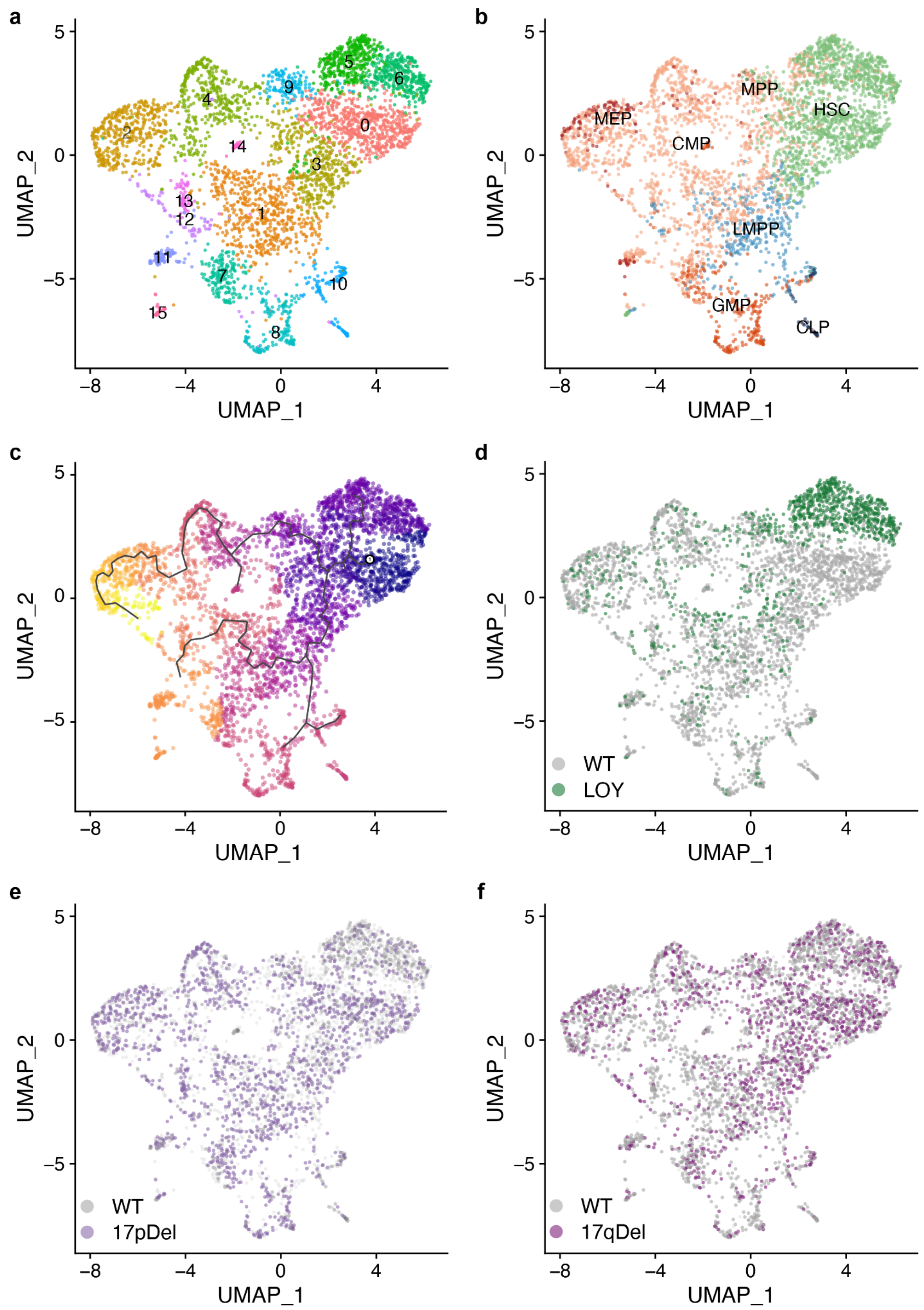


**Figure S26: Labeled UMAP projections of scRNA-seq data from BM712.**

**a)** Unsupervised clustering using Seurat[20]. **b)** Referenced-based cell-type labeling using SingleR[6], based on data from[21]. **c)** Pseudotime analysis of single-cell libraries using Monocle3[22]. Cells are ordered along the trajectory based on the decreasing number of HSCs (coloured from dark blue to yellow). **d-f)** Overlay of LOY (**d**), as well as 17p-Del (**e**) and 17q-Del (**f**) re-calls made using CONICSmat[23].

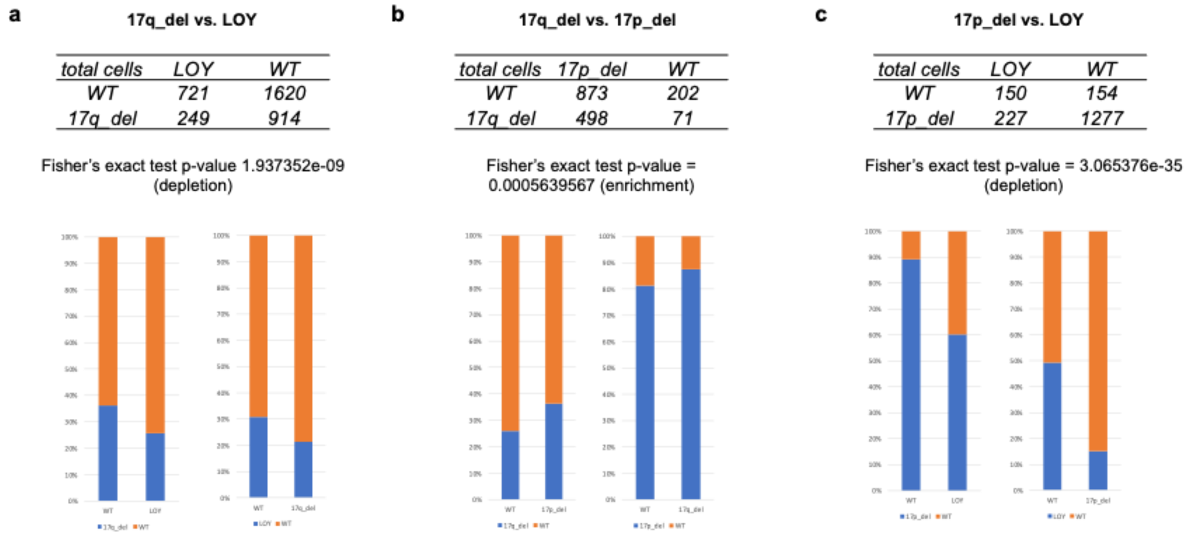


**Figure S27: Confirmation of Strand-seq-inferred clonal architecture in BM712 by targeted CNA re-calling using CONICSmat.**

**(a)** Contingency table showing the association between 17q-Del and LOY calls generated by targeted CNA recalling[23] in scRNA-seq data from BM712. Bar graphs depicting the proportion of 17q-Del calls within the WT and LOY calls (left panel), and the proportion of LOY calls within the WT and 17q-Del cells (right panel) are shown. Similar analyses were performed for the association between 17q-Del calls and 17p-Del calls **(b)**, as well as 17p-Del calls and LOY calls **(c)**. Whereas the Del events tend to co-occur, we find that LOY is inversely correlated with the presence of each deletion, confirming the BM712 clonal structure inferred based on Strand-seq (**Fig. 5c**).

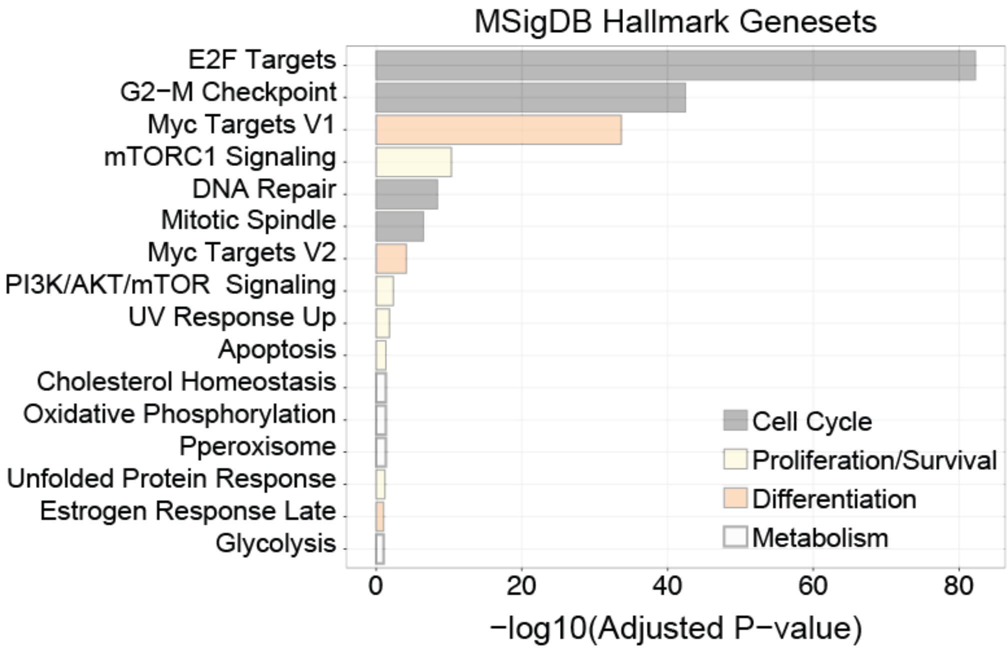


**Figure S28: Pathway enrichment analysis for 17q-Del cells from BM712.**

MSigDB Hallmarks gene set enrichment analysis for differentially expressed genes in 17q-Del cells vs WT cells, identified in scRNA-seq.

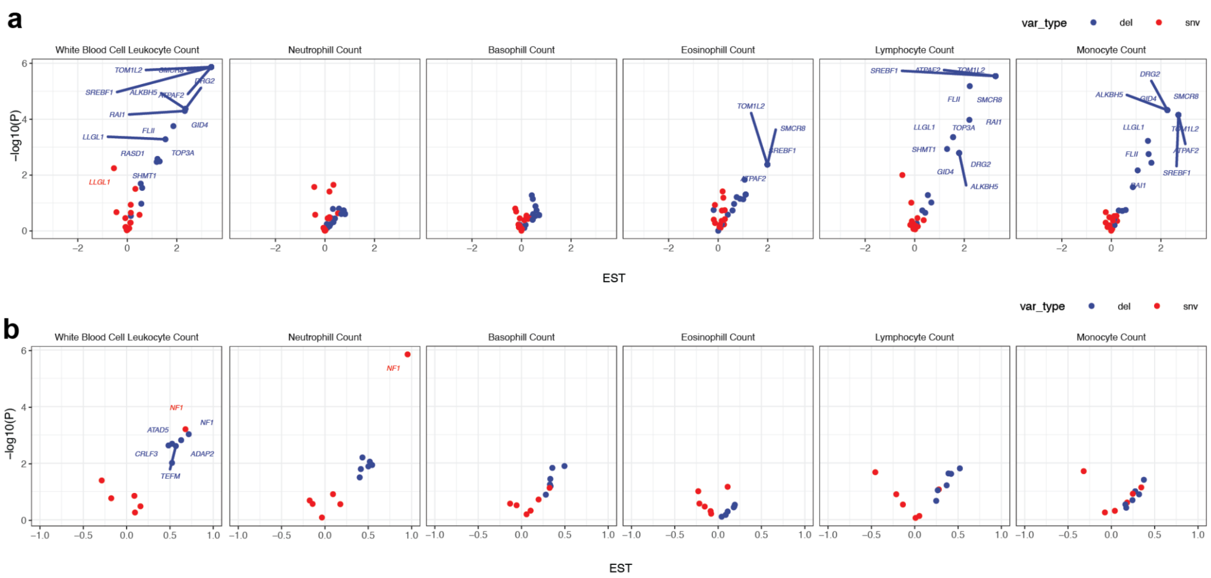


**Fig S29: Blood genotype-phenotype analysis using UK Biobank data for 17p- and 17q-Del regions.**

Volcano plots summarize burden test results for 17p-Del (**a**) and 17q-Del (**b**) based on blood count traits and genotype data from the UK Biobank (see also **Fig. 6b,c**).

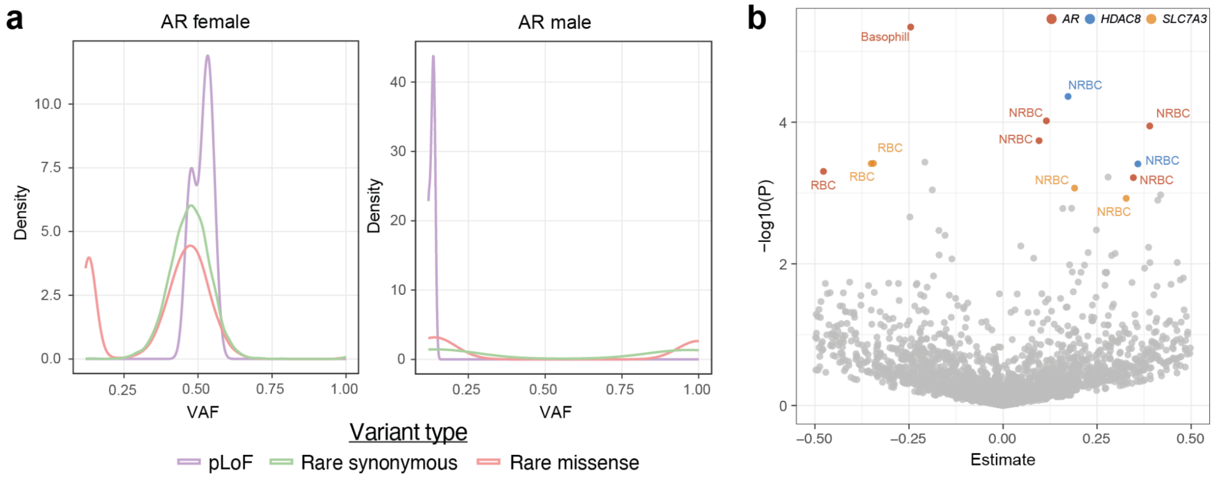


**Figure S30: Blood genotype-phenotype analysis using UK Biobank data for Xq-Inv region.**

**a**) Variant allele frequency plot for mutations in *AR*, separated by mutation type and sex. **b**) Volcano plot showing association test results of single rare missense variant at the Xq-Inv locus for all 11 blood count traits. The full respective list of missense variants analyzed is available from **Table S18**. Variants with $P_{adj}$<0.05 are colored by gene and labeled by trait: NRBC, nucleated red blood cell count; RBC, red blood cell count; basophil, basophil count. Variants with $P_{adj}$ >= 0.05 are colored in gray. Y-axis depicts nominal *P*-values.

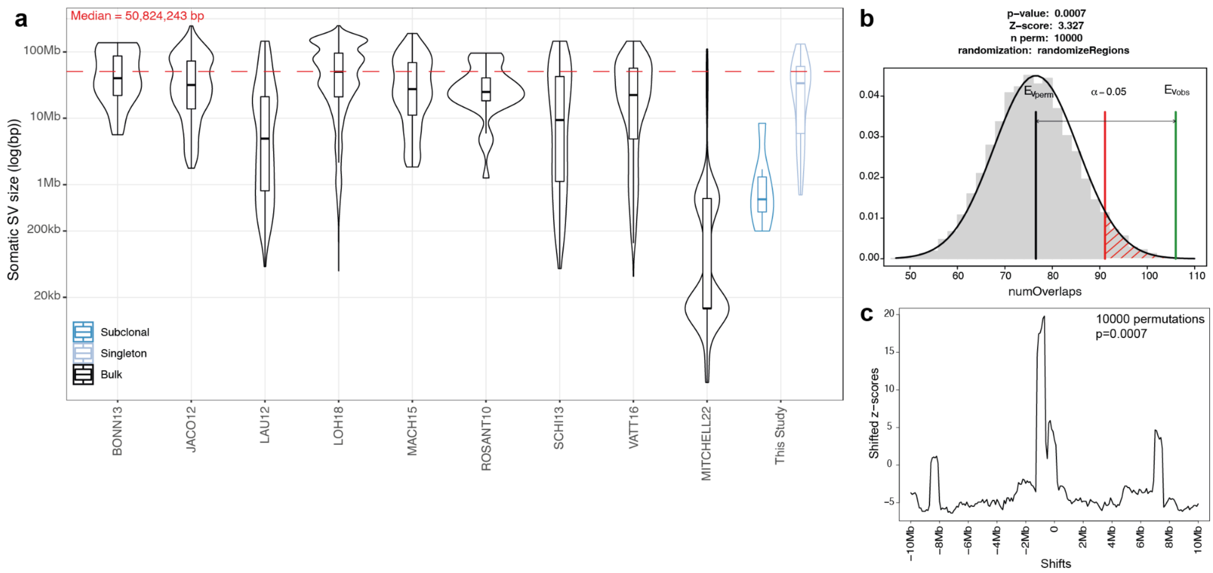


**Figure S31: Comparing previously published mosaic copy-number alteration (mCNA) data to this study.**
**a**) Violin plots showing the size distribution of mCNAs identified per study[24–32]. **b**) Permutation plot of overlaps between 200 kb regions around interstitial mCNA breakpoints from a), and SCE hotspots from this study. **c**) Local enrichment plot of breakpoints from **a-b**).

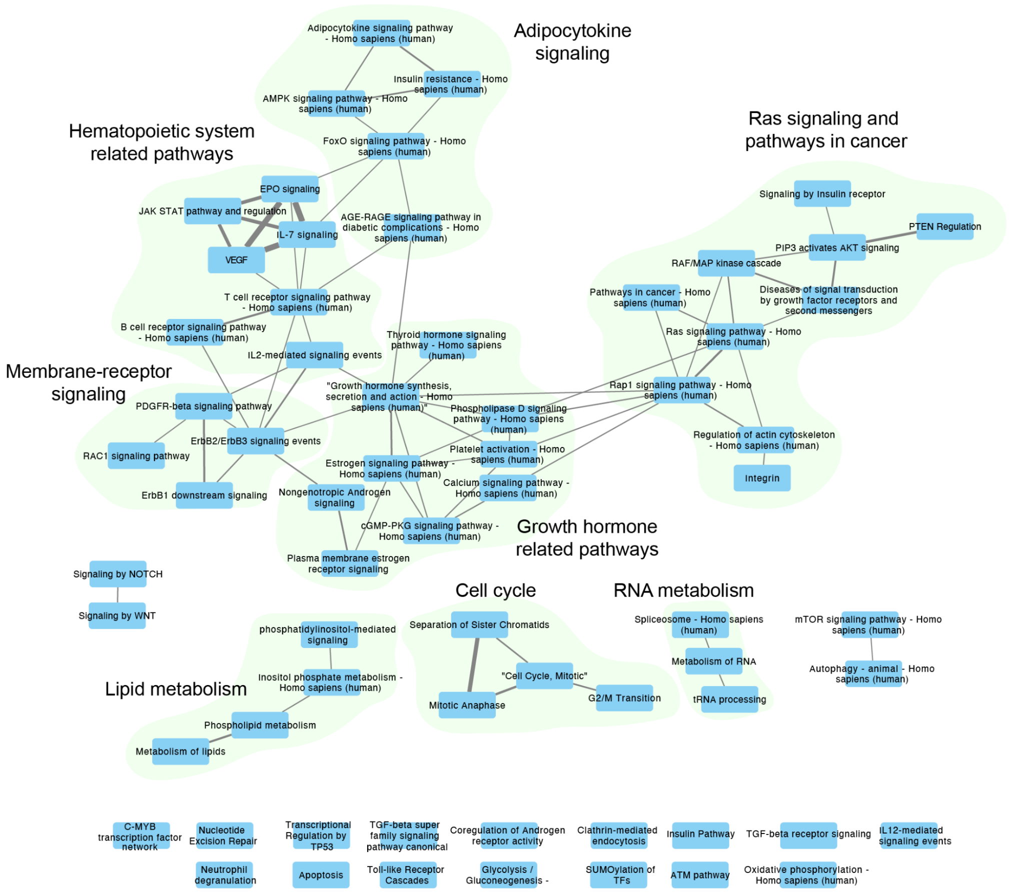


**Figure S32: Similarity analysis of 66 pathways reported in Figure S14.** In **Fig. S14**, over-represented pathways of dysregulated genes in mSV subclones are reported. To reduce the redundancy between the biological pathways and comprehensively understand the underlying hierarchical structure of the over-represented pathways, the similarity between pathways are shown as a network format in this figure. In this network, each node represents one of 66 over-represented pathways in **Fig. S14**, and the thickness of edges between the nodes represent the Jarccard similarity between them. Edges are shown when the Jaccard similarity is larger than 0.16. It resulted in the bigger category of pathways as represented in a light green background behind the nodes. We annotated the name of those bigger categories based on the common characteristics of nodes belonging to that category.

#### Supplementary Tables

**Table S1:** Donor information and fragment size distribution of Strand-seq genomic libraries generated and in this study. (Table accompanying the submission as spreadsheet).

**Table S2:** Summary of mosaicisms discovered using Strand-seq across the cohort of 19 donors. (Table accompanying the submission as spreadsheet).

**Table S3:** SCE hotspots identified in HSPCs. (Table accompanying the submission as spreadsheet).

**Table S4:** Previously described common fragile sites (CFS) in the human genome (from Palin et al., 2018). (Table accompanying the submission as spreadsheet).

**Table S5:** Immunophenotypes defining the 8 HSPC cell types represented in this study. (Table accompanying the submission as spreadsheet).

**Table S6:** scMNase-seq libraries generated for each of 8 HSPC cell types. (Table accompanying the submission as spreadsheet).

**Table S7A,B:** A) 899 genes with significant differential NO in one of the cell-types in scMNase atlas of cord blood HSPCs. B) 819 genes with significant differential NO in one of the cell-types in scMNase atlas of bone marrow HSPCs. (Tables accompanying the submission as spreadsheet).

**Table S8:** Inferred cell-type composition of Strand-seq libraries. (Table accompanying the submission as spreadsheet).

**Table S9:** Genes identified by scNOVA which have differential nucleosome occupancy between mSV-harbouring cells and WT cells, per mSV per donor. (Table accompanying the submission as spreadsheet).

**Table S10:** Genes identified by scNOVA which have differential nucleosome occupancy between mSV-harbouring cells and WT cells, per HSPC cell type with sufficient testing power per mSV per donor. (Table accompanying the submission as spreadsheet).

**Table S11:** Nearest Genes of sliding windows showing differential NO between WT and Xq-Inv cells in BM65. (Table accompanying the submission as spreadsheet).

**Table S12:** Pathway enrichment analysis for diffNO genes identified in Xq-Inv cells in BM65. (Table accompanying the submission as spreadsheet).

**Table S13:** AR target genes. (Table accompanying the submission as spreadsheet).

**Table S14:** List of genes included in the CONICSmat analysis to infer each genotype in the scRNA-seq from donor BM712. (Table accompanying the submission as spreadsheet).

**Table S15:** Differentially expressed genes per subclone from scRNA-seq of BM712. (Table accompanying the submission as spreadsheet).

**Table S16:** Geneset enrichment analysis for subclones from scRNA-seq of donor BM712. (Table accompanying the submission as spreadsheet).

**Table S17:** Over-represented pathways of dysregulated genes in mSV clones. (Table accompanying the submission as spreadsheet).

**Table S18:** Results of the UKBiobank analysis (including burden tests for genes from the 17p-Del and 17q-Del regions, and missense mutation testing from the Xq-Inv region). (Table accompanying the submission as spreadsheet).

**Table S19:** Antibody panel used to identify 8 distinct HSPCs using FACS. (Table accompanying the submission as spreadsheet).

**Table S20:** Cohort-wide enrichment analysis of dysregulated transcription factors. (Table accompanying the submission as spreadsheet).

### Supplementary Methods

##### *1.* Identification of active X chromosome from the female genome

To identify the transcriptionally active X chromosomal homolog in the cells from the female donor BM65, we performed haplotype-specific NO analysis using scNOVA, as previously described[8]. Firstly we analyzed single-cells from the Xq-Inv subclone, and resolved the reads by haplotype for chromosome X in those cells. We then extracted the haplotype-resolved NO for each gene body and calculated the mean of this value for each cell. Finally, we compared the single-cell NO from haplotype 1 and haplotype 2 using a t-test, to identify the transcriptionally active X chromosome among the two homologous chromosomes.

##### *2.* Protein-protein interaction (PPI) network analysis using STRING

To reconstruct the network between dysregulated TFs inferred by scNOVA, we used the STRING[33] multiple protein search algorithm. To evaluate the enrichment of direct interaction between dysregulated TFs, we utilized network statistics from STRING, based on the PPI enrichment *P*-value. For the analysis of BM65, we extended the network of dysregulated TFs by including their first neighbors based on the physical binding partners reported in the NCBI gene pages (Available from: https://www.ncbi.nlm.nih.gov/gene/) (**Fig. S13**). To reconstruct the network between dysregulated TFs and their first neighbors, we utilized STRING[33] as mentioned above. Pathways enriched for members of the PPI network were also identified by STRING[33].

##### *3.* Similarity analysis for over-represented pathways of dysregulated genes in mSV subclones

To reduce the redundancy between the biological pathways and comprehensively understand the underlying hierarchical structure of the over-represented pathways, we built a network model showing the links among the 66 pathways enriched by the dysregulated genes in mSV subclones (**Figure. S32**). For each pair of the 66 pathways, we computed the Jaccard coefficient and connected the two pathways with the Jaccard coefficient >= 0.16. This analysis resulted in 6 connected subnetworks. Among the connected subnetworks, the biggest subnetworks were further categorized into five network modules based on their connectivity and biological implications.

### Supplementary Notes

#### *1.* Characteristics of de novo mSVs in HSPCs

In 1 out of every 43 cells, regardless of donor age, we identify a singleton mSV. We scrutinized these singleton mSVs by utilizing single-cell tri-channel processing (scTRIP), the underlying principle of which is that each structural variant is characterized and discerned by a specific 'diagnostic footprint'[34]. The footprint encapsulates the co-segregation patterns of rearranged DNA segments, identified by sequencing single strands of each chromosome in each cell in a haplotype-resolved manner, using Strand-seq[8,34].

Our analysis revealed that singleton mSVs, but not subclonal mSVs, bear the following characteristics indicative for *de novo* DNA rearrangement:

(1) Amongst the 32 singleton mosaicisms we identify in our HSPC single-cell genomic dataset, 21 (66%) display terminal gains or losses confined to a single haplotype. In contrast, subclonal mSVs lack terminal rearrangements entirely and instead exhibit a significant enrichment for interstitial rearrangements ($P$=0.0004; Fisher's exact test) when compared to singleton mSVs. These terminal rearrangement footprints of singleton mSVs are depicted, amongst all singleton mSVs events, in **Supplemental Data File 1** as well as in **Fig. 1c**. The frequent occurrence of terminal losses and gains in singleton mSVs suggests that the derivative chromosomes emerging from these rearrangements often lack telomeric stabilization events, which may potentially increase the likelihood that more DNA rearrangements accumulate in these cells (**Fig. 1c,f**)[35].

(2) Three singleton mSVs exhibit characteristics of complex chromosomal rearrangements, encompassing mSVs triggered by breakage-fusion-bridge (BFB) cycles[34,36] and an amplification induced by terminal sister chromatid fusion, leading to a sevenfold increase in copy number (**Fig. 1c**). The latter rearrangement event could stem from a BFB process occurring in the absence of telomere stabilization[35].

(3) Our observations, as delineated in **Fig. 1j, Fig. S7** and in the main text, show instances of SCEs occurring in the same cell and haplotype directly at the breakpoints of singleton mSVs (we did not, by comparison, observe significant colocalisation with SCEs for the breakpoints of subclonal mSVs). This indicates a link between SCE formation and DNA rearrangement processes resulting in mSV formation[37–39].

(4) On average, singleton mSVs are ~17.6 times larger than subclonal mSVs (mean size of 36.9 and 2.1 Megasbasepairs (Mb), respectively; $P$=0.0009, Wilcoxon rank-sum test; **Fig. 1d**). Due to their substantial size, these mSVs result in significant autosomal aneuploidy, often tens of megabasepairs in length, which is likely to be detrimental for their clonal expansion given the adverse effects of autosomal aneuploidy in normal cells[40].

We therefore infer that the singleton mSVs identified in our HSPC dataset predominantly represent *de novo* mSV formation events. Considering their distinct characteristics, such as substantial regions of autosomal aneuploidy and likely lacking telomeric functionality on the affected homolog, it is plausible that most newly formed mSVs are incapable of reaching considerable subclonal frequencies in normal HSPCs.

#### *2.* Comparison with prior surveys of mosaic copy-number alterations and mSVs

Placing our findings into the larger scope of research on clonal hematopoiesis and its previously known association with mosaic CNAs (*i.e.* copy-imbalanced mSVs), we find that the subclonal mSVs identified in our study through single-cell genomic sequencing (Strand-seq) are significantly smaller (**Fig. S31**) than CNAs detected in surveys based on utilizing blood cells in bulk (primarily pursued using microarray based hybridization)[24–32]. This observation is consistent with the high genomic resolution of Strand-seq, which exceeds bulk approaches for detecting certain subclonal structural variant classes, particularly in the case of sub-Megabase sized somatic variants[34]. By comparison, the subclonal mSVs detected in our study fall into the same size range as for one study, by Mitchell and colleagues[25], who undertook WGS of clones derived from single HSPCs. These data suggest that the mSVs identified in our study as well as by Mitchell et al. may have escaped detection in prior CNA-focused studies using bulk hybridization-based assays. We note that Mitchell et al. and our study are the only two surveys specifically examining HSPCs, rather than the peripheral blood, leaving the possibility that different cell types are differentially impacted by mSVs.

To bolster our findings relating to common fragile sites (CFSs) and their association with mSVs, we performed additional analyses on the aforementioned prior data on mosaic CNAs[24–32]. We permuted breakpoints from previously reported mosaic CNAs against the SCE hotspots identified in our study, identifying a similar trend to that in our own data, whereby mosaic CNA breakpoints demonstrate significant local enrichment at SCE hotspots (**Fig. S31**). This observation suggests that genomic loci in HSPCs, predisposed to mSV formation, also denote fragile regions prone to CNAs in peripheral blood.

#### *3.* Investigation of genes associated with local effect of subclonal inversion in BM65

Amongst the genes our analysis associates with a local effect of the subclonal inversion in BM65, 3 genes – which include *AR*, the top hit, as well as *HDAC8* and *MAGEE1* – are inferred to be more active in mSV cells, whereas the remaining 10 genes are inferred to be downregulated in mSV cells. 5/13 genes including *AR*, *EDA2R*, *SLC7A3*, *HDAC8*, and *CHIC1* show expression in HSPCs according to previously published bulk RNA-seq data from HSPCs[41].

#### *4.* Investigation of functional links between dysregulated TFs in the 17p-Del subclone in BM712

Prior reports have tied *SREBF1* knockout in murine blood cells to an increase in primitive HSPCs[42], and our findings closely mirror these data, with 17p-Del cells showing enrichment for both HSCs and CMPs (**Fig. 5a**). Protein-protein interaction mapping of the dysregulated TFs using STRING[33] (**Supplementary Methods)** revealed significant functional associations between *SREBF1* and six additional TFs, which suggests that *SREBF1* cooperates with functionally-related TFs that become dysregulated in association with 17p-Del ($P$=3.57e-08; **Fig. S22**). Collectively, these data implicate the somatic hemizygous loss of *SREBF1* as a putative driver of gene dysregulation in 17p-Del-bearing cells. SREBF1 (also known as SREBP1) has been reported to induce PPARG expression[43,44]. Furthermore, PPARG activates CREB1 expression by binding its promoter[45,46], in line with the tight functional connections between the TFs identified in the STRING based PPI analysis, and in support of a possible causal role of *SREBF1* deletion in mediating the molecular phenotype seen in BM712.

#### *5.* Potential small deletions at regions of recurrent SCE/mSV formation

We observed marked localized 'fragility' at the *FRA3B* locus in donor BM762 (**Fig. 1j**). Although this donor showed similar SCE counts to the other samples (**Fig. S3**), nine single cells exhibit an SCE, an mSV, or both within a 500 kb region of this CFS. One cell shows two SCEs at *FRA3B*, one on each homolog, while another harbors a terminal deletion originating from the same locus (**Fig. 1j, S7**). A closer look at *FRA3B* at sub-Mb resolution reveals potential small deletions (< 200 kb), which are below our mSV discovery resolution[34], aligning with SCEs in the same cells (**Fig. 1j; S7**). Manual inspection shows that these putative small deletions arise in 37.5% (3/8) of cells with an SCE versus 1.89% (1/53) without an SCE ($P$=0.0055; Fisher's exact test).

##### *6.* Analysis of somatic SNVs from the IntoGen Clonal Hematopoiesis Mutation Browser

We analysed data from the IntOGen Clonal Hematopoiesis Mutation Browser[47], which at the time of analysis (9th June 2023) reported on 175 mosaic SNVs falling into the *NF1* gene. We find a marked abundance of potentially deleterious variants (27% [47/175]) amongst these SNVs. This includes predicted loss-of-function (pLoF) SNVs (frameshift variant: 5.1% [9/175]; stop gained: 9.7% [17/175]) and SNVs predicted to affect splicing and thus potentially resulting in *NF1* transcripts generated from an aberrant open reading frame[48] (splice donor variant: 6.3% [11/175]; splice acceptor variant: 3.4% [6/175]; splice region variant: 2.3% [4/175]). These data suggest that *NF1* hemizygous loss could fuel clonal hematopoiesis.

##### *7.* Analysis of somatic SNVs affecting the AR gene in the UK Biobank

The *AR* gene has a sex-specific molecular biology[49], and shows highly sex-specifc SNV distributions in the UK Biobank cohort (for example, *AR* pLoF SNVs are seen in females (**Fig. 6d**), whereas they are essentially absent in males from the UK Biobank cohort). Therefore, our comprehensive analysis of presumed mosaic SNVs within the *AR* gene employed a sex-specific approach. We utilized sex as a covariate in the employed multiple linear regression model; in addition, we built two separate models for males and females, respectively. We focused our analysis of the *AR* gene on females, as the mosaic inversion was identified in a female (BM65).

#### References


1. Benjamini, Y. & Hochberg, Y. Controlling the False Discovery Rate: A Practical and Powerful Approach to Multiple Testing. *J. R. Stat. Soc. Series B Stat. Methodol.* **57**, 289–300 (1995).
2. Palin, K. *et al.* Contribution of allelic imbalance to colorectal cancer. *Nat. Commun.* **9**, 3664 (2018).
3. Buenrostro, J. D. *et al.* Integrated Single-Cell Analysis Maps the Continuous Regulatory Landscape of Human Hematopoietic Differentiation. *Cell* **173**, 1535–1548.e16 (2018).
4. Gudmundsson, K. O. *et al.* Prdm16 is a critical regulator of adult long-term hematopoietic stem cell quiescence. *Proc. Natl. Acad. Sci. U. S. A.* **117**, 31945–31953 (2020).
5. Bunis, D. G. *et al.* Single-Cell Mapping of Progressive Fetal-to-Adult Transition in Human Naive T Cells. *Cell Rep.* **34**, 108573 (2021).
6. Aran, D. *et al.* Reference-based analysis of lung single-cell sequencing reveals a transitional profibrotic macrophage. *Nat. Immunol.* **20**, 163–172 (2019).
7. Martens, J. H. A. & Stunnenberg, H. G. BLUEPRINT: mapping human blood cell epigenomes. *Haematologica* **98**, 1487–1489 (2013).
8. Jeong, H. *et al.* Functional analysis of structural variants in single cells using Strand-seq. *Nat. Biotechnol.* (2022) doi:10.1038/s41587-022-01551-4.
9. Wingender, E., Dietze, P., Karas, H. & Knüppel, R. TRANSFAC: a database on transcription factors and their DNA binding sites. *Nucleic Acids Res.* **24**, 238–241 (1996).
10. Sandelin, A., Alkema, W., Engström, P., Wasserman, W. W. & Lenhard, B. JASPAR: an open-access database for eukaryotic transcription factor binding profiles. *Nucleic Acids Res.* **32**, D91–4 (2004).
11. Sharma, N. V. *et al.* Identification of the Transcription Factor Relationships Associated with Androgen Deprivation Therapy Response and Metastatic Progression in Prostate Cancer. *Cancers* **10**, (2018).
12. Kamburov, A. & Herwig, R. ConsensusPathDB 2022: molecular interactions update as a resource for network biology. *Nucleic Acids Res.* **50**, D587–D595 (2022).

13. Lachmann, A. *et al.* ChEA: transcription factor regulation inferred from integrating genome-wide ChIP-X experiments. *Bioinformatics* **26**, 2438–2444 (2010).

14. Fishilevich, S. *et al.* GeneHancer: genome-wide integration of enhancers and target genes in GeneCards. *Database* **2017**, (2017).

15. Takeda, D. Y. *et al.* A Somatically Acquired Enhancer of the Androgen Receptor Is a Noncoding Driver in Advanced Prostate Cancer. *Cell* **174**, 422–432.e13 (2018).

16. Hammal, F., de Langen, P., Bergon, A., Lopez, F. & Ballester, B. ReMap 2022: a database of Human, Mouse, Drosophila and Arabidopsis regulatory regions from an integrative analysis of DNA-binding sequencing experiments. *Nucleic Acids Res.* **50**, D316–D325 (2022).

17. Rausch, T. *et al.* DELLY: structural variant discovery by integrated paired-end and split-read analysis. *Bioinformatics* **28**, i333–i339 (2012).

18. Rivals, I., Personnaz, L., Taing, L. & Potier, M.-C. Enrichment or depletion of a GO category within a class of genes: which test? *Bioinformatics* **23**, 401–407 (2006).

19. ICGC/TCGA Pan-Cancer Analysis of Whole Genomes Consortium. Pan-cancer analysis of whole genomes. *Nature* **578**, 82–93 (2020).

20. Butler, A., Hoffman, P., Smibert, P., Papalexi, E. & Satija, R. Integrating single-cell transcriptomic data across different conditions, technologies, and species. *Nat. Biotechnol.* **36**, 411–420 (2018).

21. Xie, X. *et al.* Single-cell transcriptomic landscape of human blood cells. *Natl Sci Rev* **8**, nwaa180 (2021).

22. Trapnell, C. *et al.* The dynamics and regulators of cell fate decisions are revealed by pseudotemporal ordering of single cells. *Nat. Biotechnol.* **32**, 381–386 (2014).

23. Müller, S., Cho, A., Liu, S. J., Lim, D. A. & Diaz, A. CONICS integrates scRNA-seq with DNA sequencing to map gene expression to tumor sub-clones. *Bioinformatics* **34**, 3217–3219 (2018).

24. Loh, P.-R. *et al.* Insights into clonal haematopoiesis from 8,342 mosaic chromosomal alterations. *Nature* **559**, 350–355 (2018).

25. Mitchell, E. *et al.* Clonal dynamics of haematopoiesis across the human lifespan. *Nature* **606**, 343–350 (2022).

26. Vattathil, S. & Scheet, P. Extensive Hidden Genomic Mosaicism Revealed in Normal Tissue. *Am. J. Hum. Genet.* **98**, 571–578 (2016).

27. Machiela, M. J. *et al.* Characterization of large structural genetic mosaicism in human autosomes. *Am. J. Hum. Genet.* **96**, 487–497 (2015).

28. Bonnefond, A. *et al.* Association between large detectable clonal mosaicism and type 2 diabetes with vascular complications. *Nat. Genet.* **45**, 1040–1043 (2013).

29. Schick, U. M. *et al.* Confirmation of the reported association of clonal chromosomal mosaicism with an increased risk of incident hematologic cancer. *PLoS One* **8**, e59823 (2013).

30. Jacobs, K. B. *et al.* Detectable clonal mosaicism and its relationship to aging and cancer. *Nat. Genet.* **44**, 651–658 (2012).

31. Laurie, C. C. *et al.* Detectable clonal mosaicism from birth to old age and its relationship to cancer. *Nat. Genet.* **44**, 642–650 (2012).

32. Rodríguez-Santiago, B. *et al.* Mosaic uniparental disomies and aneuploidies as large structural variants of the human genome. *Am. J. Hum. Genet.* **87**, 129–138 (2010).

33. Szklarczyk, D. *et al.* The STRING database in 2021: customizable protein-protein networks, and functional characterization of user-uploaded gene/measurement sets. *Nucleic Acids Res.* **49**, D605–D612 (2021).

34. Sanders, A. D. *et al.* Single-cell analysis of structural variations and complex rearrangements with tri-channel processing. *Nat. Biotechnol.* **38**, 343–354 (2020).

35. Cosenza, M. R., Rodriguez-Martin, B. & Korbel, J. O. Structural Variation in Cancer: Role, Prevalence, and Mechanisms. *Annu. Rev. Genomics Hum. Genet.* **23**, 123–152 (2022).

36. McClintock, B. The Stability of Broken Ends of Chromosomes in Zea Mays. *Genetics* **26**, 234–282 (1941).

37. Dillon, L. W., Burrow, A. A. & Wang, Y.-H. DNA instability at chromosomal fragile sites in cancer. *Curr. Genomics* **11**, 326–337 (2010).

38. Glover, T. W. & Stein, C. K. Induction of sister chromatid exchanges at common fragile sites. *Am. J. Hum. Genet.* **41**, 882–890 (1987).

39. Liu, P., Carvalho, C. M. B., Hastings, P. J. & Lupski, J. R. Mechanisms for recurrent and

complex human genomic rearrangements. *Curr. Opin. Genet. Dev.* **22**, 211–220 (2012).

40. Tang, Y.-C. & Amon, A. Gene copy-number alterations: a cost-benefit analysis. *Cell* **152**, 394–405 (2013).

41. Corces, M. R. *et al.* Lineage-specific and single-cell chromatin accessibility charts human hematopoiesis and leukemia evolution. *Nat. Genet.* **48**, 1193–1203 (2016).

42. Lu, Y. *et al.* Srebf1c preserves hematopoietic stem cell function and survival as a switch of mitochondrial metabolism. *Stem Cell Reports* **17**, 599–615 (2022).

43. Teresi, R. E., Planchon, S. M., Waite, K. A. & Eng, C. Regulation of the PTEN promoter by statins and SREBP. *Hum. Mol. Genet.* **17**, 919–928 (2008).

44. Kim, J. B. & Spiegelman, B. M. ADD1/SREBP1 promotes adipocyte differentiation and gene expression linked to fatty acid metabolism. *Genes Dev.* **10**, 1096–1107 (1996).

45. Rudko, O. I., Tretiakov, A. V., Naumova, E. A. & Klimov, E. A. Role of PPARs in Progression of Anxiety: Literature Analysis and Signaling Pathways Reconstruction. *PPAR Res.* **2020**, 8859017 (2020).

46. Mäkelä, J. *et al.* Peroxisome proliferator-activated receptor-γ (PPARγ) agonist is neuroprotective and stimulates PGC-1α expression and CREB phosphorylation in human dopaminergic neurons. *Neuropharmacology* **102**, 266–275 (2016).

47. Pich, O., Reyes-Salazar, I., Gonzalez-Perez, A. & Lopez-Bigas, N. Discovering the drivers of clonal hematopoiesis. *Nat. Commun.* **13**, 4267 (2022).

48. Anna, A. & Monika, G. Splicing mutations in human genetic disorders: examples, detection, and confirmation. *J. Appl. Genet.* **59**, 253–268 (2018).

49. Bennett, N. C., Gardiner, R. A., Hooper, J. D., Johnson, D. W. & Gobe, G. C. Molecular cell biology of androgen receptor signalling. *Int. J. Biochem. Cell Biol.* **42**, 813–827 (2010).
